# A multispecies foundation model for genome function prediction, gene annotation and sequence design

**DOI:** 10.64898/2025.12.22.695963

**Authors:** Sam Boshar, Benjamin Evans, Ziqi Tang, Yanis Adel, Yohan Bornachot, Shawn Sidbon, Franziska K. Lorbeer, Armand Picard, Chandana Rajesh, Evan Seitz, David Emms, Javier Mendoza-Revilla, Ariana Hernandez, Fatimah Al-Ani, Tristan Karch, Yair Schiff, Marie Lopez, Alexandre Laterre, Karim Beguir, Peter Koo, Volodymyr Kuleshov, Alexander Stark, Bernardo P. de Almeida, Thomas Pierrot

## Abstract

Comparative genomics and synthetic biology require models that can interpret, annotate, and design DNA sequences across species, yet genomic models have not combined these capabilities beyond a few well-characterized organisms. Here we introduce Nucleotide Transformer v3 (NTv3), a long-context cross-kingdom foundation model that provides a common backbone for functional prediction, gene annotation and controllable sequence generation across phylogenetically diverse species. NTv3 is pretrained on more than 128,000 genomes using sequences up to 1 Mb and post-trained on 16,262 functional tracks across 15 animal and plant genomes. NTv3 performs comparably to AlphaGenome on matched human assays and matches or exceeds evaluated published models for functional-track prediction across 19 species spanning mammals, birds, fish, plants, fungi and protists. Ablation experiments show that both genome pre-training and expanding functional supervision from human and mouse to 15 species improve track prediction. Leveraging these learned functional representations, NTv3 outperforms leading *ab initio* gene predictors across mammals, plants and insects, with fewer spurious gene models and improved recovery and reduced fragmentation of long genes. Finally, the same representations can be used with discrete diffusion to design regulatory DNA sequences with specific activity levels. We demonstrate this in plant and fly promoters and enhancers, and experimentally validate the activity and promoter selectivity of the generated fly enhancers. Together, NTv3 unifies genome interpretation, gene annotation and regulatory sequence design across the animal and plant kingdoms, with direct relevance to agriculture, biodiversity research and biotechnology.

## 1. Introduction

Comparative genomics and synthetic biology pose complementary challenges: interpreting natural genomes across diverse species and designing new DNA with specified functions. Four modeling paradigms address connected aspects of these challenges: 1) genome pre-training learns sequence features from large collections of unlabeled genomes, 2) functional-track prediction grounds those representations in experimentally measured transcriptional and regulatory activity, 3) genome annotation assembles sequence signals into complete gene models, and 4) regulatory-sequence design inverts sequence–function relationships to produce new DNA sequences. Although biologically connected, these paradigms have largely been developed separately, limiting the transfer of information between genome collections, functional measurements, genome annotations and sequence-design experiments.

Functional-track prediction provides a natural bridge between genome-scale representation learning and biological interpretation. Supervised sequence-to-function models learn to predict measurements of transcription, chromatin accessibility, histone modifications, and protein binding directly from DNA [1–14]. Joint training on human and mouse established that regulatory information can transfer between genomes, while Enformer, Borzoi and AlphaGenome subsequently increased sequence context and the breadth or resolution of the predicted modalities [5, 6, 15, 16]. However, training on large, diverse collections of functional tracks remains concentrated principally in human and mouse, and it remains unclear whether functional measurements from phylogenetically diverse genomes can provide transferable regulatory information and improve prediction in well-characterized species while enabling regulatory interpretation in species with limited functional data.

Self-supervised genomic foundation models address a complementary limitation by learning from unlabeled genomes at much greater phylogenetic scale [17–24]. Nucleotide Transformer showed that such masked language modeling yields transferable representations of genomic sequence [19], and Evo models extended long-context genome modeling to prediction and generation [21, 25]. These approaches can capture sequence regularities across broad evolutionary diversity, but sequence pre-training alone does not directly connect those regularities to measured genome function. Conversely, supervised functional models learn from measurements of genome function but are trained on few species. It therefore remains unknown whether the two kinds of training are complementary when combined in one long-context model, and whether functional data from distant taxa reveal regulatory principles that transfer across species.

Gene annotation has followed a largely separate trajectory, progressing from probabilistic models such as AUGUSTUS [26] to deep-learning gene callers including Helixer and Tiberius [27–29]. Pre-trained representations have also been adapted to nucleotide-level semantic segmentation with SegmentNT [30], while PlantCAD2 and GeneCAD extended foundation-model approaches to gene annotation in plants [31, 32]. Nevertheless, existing *ab initio* gene predictors learn principally from genome sequence and reference annotations and have not been combined with models first trained to predict measured transcriptional and regulatory activity across species. Regulatory-sequence design is similarly disconnected from genome-scale functional prediction and annotation. Most approaches use task-specific sequence-to-activity models, optimize candidates against external predictors or generate sequences using general genome language models [3, 25, 33]. Consequently, information learned through functional-track prediction rarely transfers directly to complete gene annotation or experimentally validated regulatory-sequence design.

Comparative genomics and synthetic biology therefore lack a common framework connecting genome pretraining, multispecies functional-track prediction, structural genome annotation and regulatory-sequence design. To our knowledge, no study has tested whether broad genome self-supervision can be grounded through functional measurements from phylogenetically diverse species, whether the resulting functional representation improves *ab initio* prediction of complete coding gene models, and whether the same model lineage can be adapted to design regulatory DNA with experimentally verified properties.

Here, we introduce *Nucleotide Transformer v3* (NTv3), a cross-kingdom foundation model family designed to connect all four paradigms, from 1) genome pre-training and 2) functional-track prediction, to 3) genome annotation and 4) regulatory-sequence design, through a common model trunk and successive training stages (Fig. 1). NTv3 is first pre-trained by masked language modeling on more than 128,000 genomes with sequence contexts up to 1 Mb, followed by post-training on 16,262 functional genomic tracks across fifteen animal and plant genomes. This functional representation can subsequently be adapted to *ab initio* prediction of complete gene models from DNA alone or converted into a conditional regulatory-sequence generator through masked diffusion language modeling [34].

**Figure 1.**
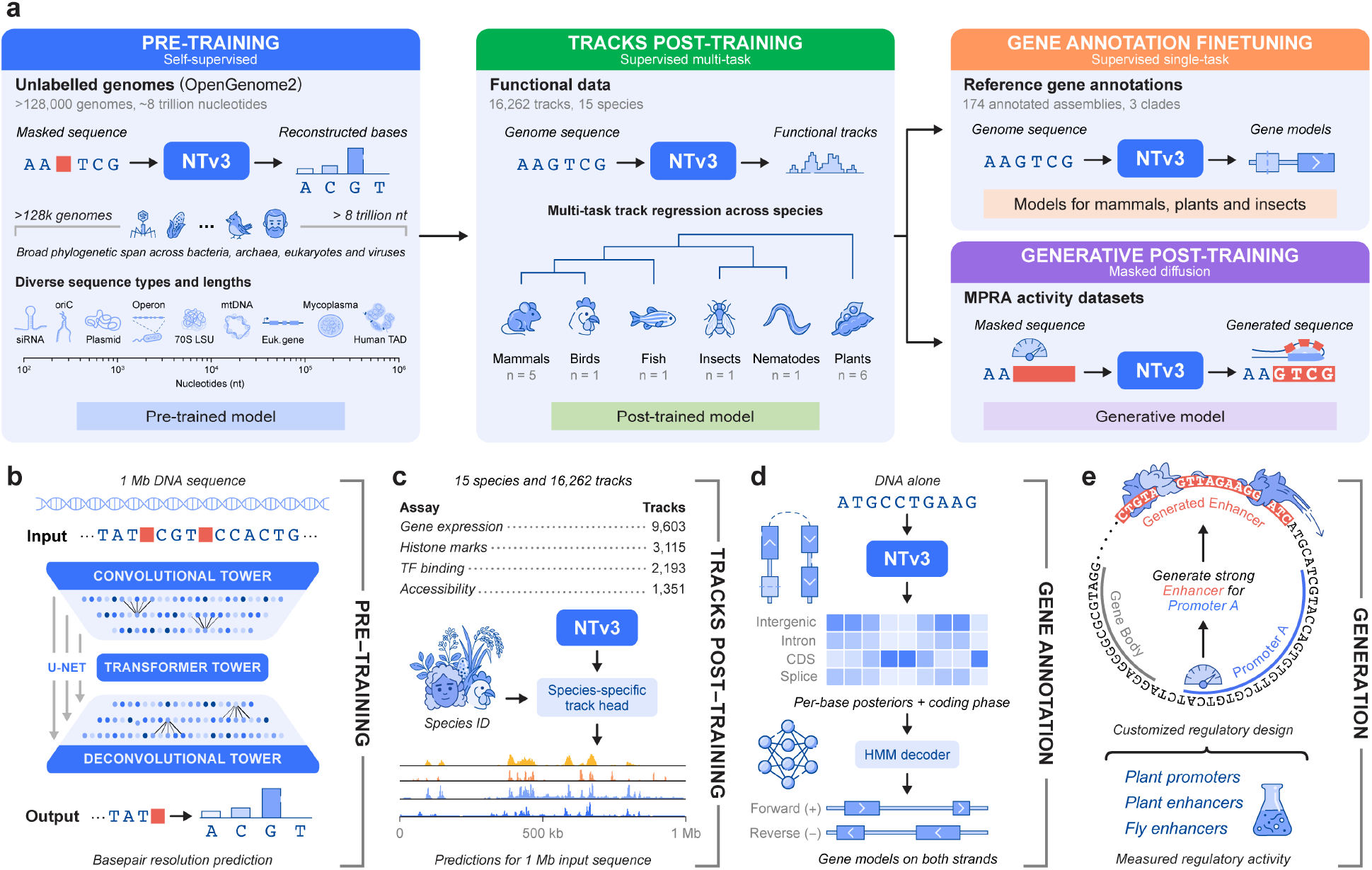
NTv3 connects genome pre-training, multispecies functional-track prediction, gene annotation and regulatory-sequence design. **a** Training lineage. Masked language modeling on more than 128,000 genomes is followed by supervised post-training on 16,262 functional tracks across 15 species. The resulting checkpoint is adapted separately for gene annotation using reference annotations and for sequence generation using masked diffusion and massively parallel reporter assay activity data. **b** NTv3 is composed of a long-context U-Net architecture that combines convolutional towers with a transformer bottleneck and produces base-pair-resolution outputs from sequences up to 1 Mb. Pre-training is done through masked language modeling. **c** NTv3 predicts functional genomic tracks across gene-expression, chromatin and transcription-factor-binding assays on 15 species. **d** NTv3 predicts per-base annotation states from DNA alone and decodes complete gene models on both strands into an output GFF3 file, with models trained for mammals, plants and insects. **e** NTv3 designs regulatory sequences conditioned on activity and sequence context in plant-promoter, plant-enhancer and fly-enhancer reporter systems.

We demonstrate that NTv3 achieves the highest performance on functional tracks across 19 species spanning mammals, birds, fish, plants, fungi and protists, while matching AlphaGenome [6] on matched human assays. Controlled ablations show that genome self-supervision and multispecies functional supervision each improve functional-track prediction, demonstrating that broader species coverage contributes transferable regulatory information. On genome annotation, NTv3 improves gene prediction across mammals, plants and insects, including the recovery of long mammalian genes. Finally, masked-diffusion adaptation enables conditional design of plant and fly regulatory elements. In both plants and flies, independent predictive models indicate that the designs follow the requested activity. We experimentally tested the activity of the fly enhancers in *Drosophila* cells and show that they reproduce the requested ordering and preference for the intended promoter. Together, these results connect phylogenetic-scale pre-training, multispecies regulatory interpretation, structural genome annotation and experimentally validated sequence design within a common foundation-model framework.

## 2. Results

### 2.1. Genome pre-training and multispecies supervision improve functional-track prediction

NTv3 brings two innovations to previous sequence-to-function models: it combines self-supervised pre-training on phylogenetically diverse genomes with supervised post-training on functional data from 15 species, adding 13 species to the human–mouse pair used by most large sequence-to-function models [5, 6, 15, 16]. We demonstrate below with different ablations that both additions improve track prediction in human as well as in the other species, in particular on gene expression.

NTv3 accepts DNA sequences of up to 1 Mb and produces base-pair-resolution predictions over the target region using a U-Net-inspired architecture (Fig. 1b; Methods 4.1.1) [5, 6]. A convolutional tower progressively compresses the input sequence, a transformer bottleneck integrates information across the full context, and a symmetric upsampling tower restores the original resolution. This architecture combines local sequence processing with the long-range context required to model distal regulatory dependencies. Species identity enters through adaptive layer normalization in the prediction module, whereas the shared model trunk receives sequence alone. Although U-Net-style architectures have previously been used for supervised sequence-to-function modeling, here we additionally pre-train the model trunk at genome scale.

Pre-training uses OpenGenome2 [25], comprising more than 8 trillion nucleotides from over 128,000 genomes across all domains of life. We pre-train NTv3 by masked language modeling (MLM) [35] using a length curriculum that begins with 1–8-kb windows and progressively doubles the maximum context to 1 Mb while continuing to sample shorter sequences (Methods 4.2.1, 4.2.6). We subsequently post-train the model on 16,262 functional genomic tracks from 15 animal and plant genomes (human, mouse, pig, sheep, horse, chicken, zebrafish, fly, worm, *Arabidopsis Thaliana*, rice, maize, wheat, soybean and cotton) (Fig. 2b; Methods 4.3, 4.3.1). The dataset spans chromatin accessibility, histone modifications, transcription-factor binding and transcriptional activity. These two training stages and multispecies datasets allow NTv3 to learn shared regulatory principles while remaining sensitive to species-specific genome organization. We also release a variant of NTv3 further fine-tuned on a mixture of input lengths from 16 kb to 1 Mb to facilitate its use on shorter sequences (Methods 4.3.9; Supplementary Fig. A.4).

**Figure 2.**
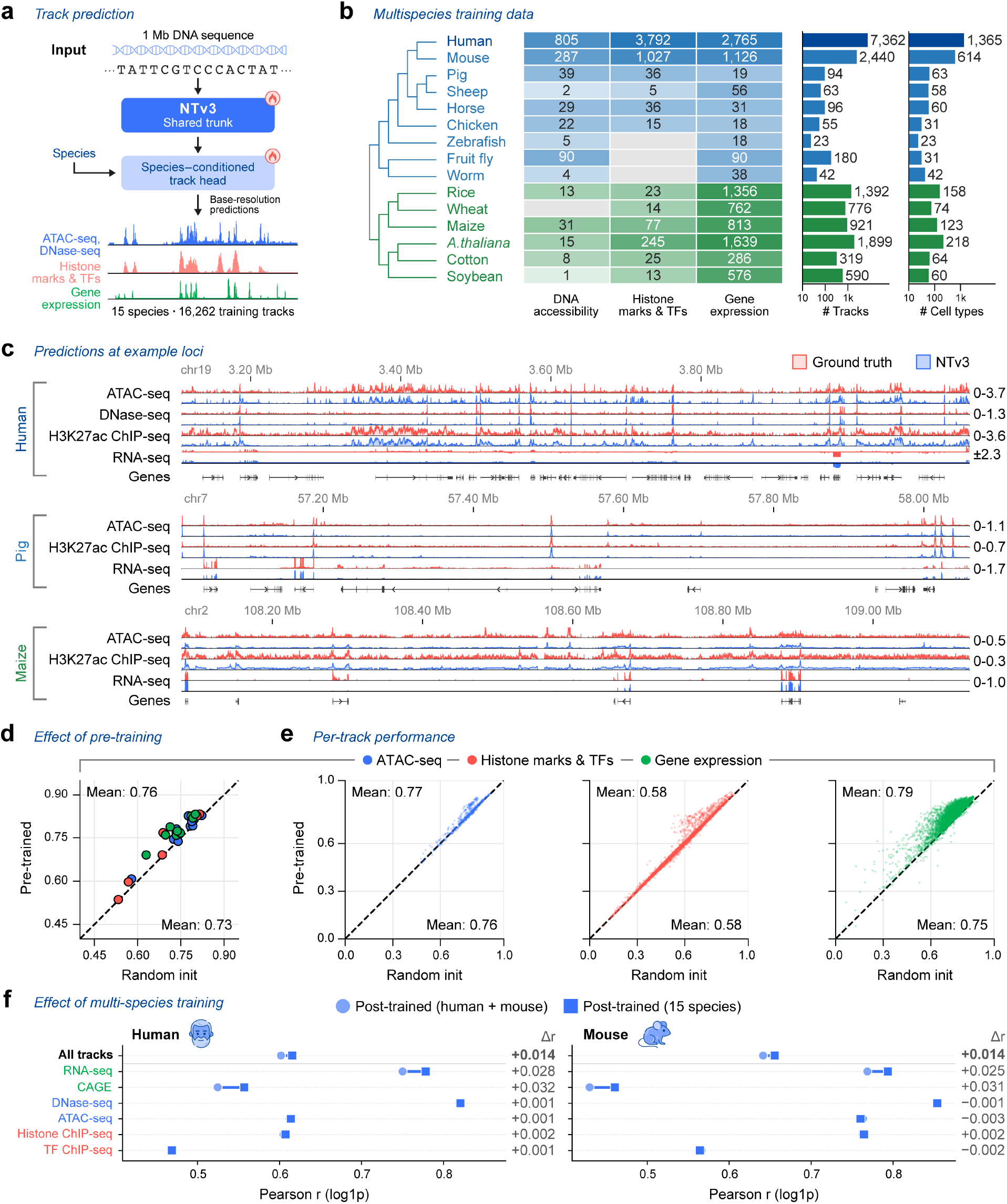
NTv3 architecture and effects of genome pre-training and multispecies supervision. **a** NTv3 architecture. A shared U-Net trunk maps a 1 Mb DNA sequence to base-resolution functional-track predictions. The trunk comprises a convolutional downsampling tower, a transformer bottleneck and a convolutional upsampling tower. Species identity conditions the prediction head through adaptive layer normalization. **b** Composition of the post-training dataset, showing the 16,252 tracks in the three assay classes plotted here, of the 16,262 used for post-training (mouse GRO-seq tracks not shown). The matrix shows the number of tracks by species and assay class; bars show the total numbers of tracks and cell types for each species. Species are ordered phylogenetically, with animals in blue and plants in green. **c** Observed signal (red) and NTv3 predictions (blue) at representative 1 Mb loci from human liver, pig liver and maize endosperm. Tracks are displayed after log-transformation and aggregation into 96-bp bins; the plotted ranges are indicated beside each track. Ensembl gene annotations are shown below. **d** Effect of genome pre-training across species and assay classes. Each point represents the mean Pearson correlation for one species–assay combination (*n* = 41); the dashed line denotes equal performance. **e** Track-level comparison of pre-trained and randomly initialized models, stratified by assay class. Each point represents one functional track, annotations report the mean across tracks and dashed lines denote equal performance. **f** Effect of multispecies supervision on held-out human and mouse test windows. Paired points compare models trained on human and mouse with models trained on all 15 species. Values at right report the difference in Pearson correlation. Panels **d**–**f** report Pearson correlations on held-out test windows.

NTv3 recovered assay- and cell-type-specific functional profiles across species (Fig. 2c). Across 41 species–assay groups, initialization from the pre-trained model improved performance over random initialization in 40 groups, increasing the mean Pearson correlation by 0.03 (Fig. 2d). The gain was greatest for RNA-seq tracks (0.04), compared with 0.01 for ATAC-seq and 0.008 for histone-modification and transcription-factor-binding tracks (Fig. 2e; Supplementary Table C.5). Because RNA-seq and CAGE together represented 59% of the post-training corpus, this assay-specific difference may partly reflect their greater representation during training.

Expanding supervised post-training from human and mouse to all 15 species produced an additional improvement in expression-related predictions. On held-out human windows, the 15-species model increased Pearson correlation by 0.032 for CAGE and 0.028 for RNA-seq relative to the human-and-mouse model (Fig. 2f; Supplementary Table C.6). By contrast, the differences for DNase-seq, ATAC-seq, histone ChIP-seq and transcription-factor ChIP-seq were all within 0.003. The same qualitative pattern was observed in mouse. Thus, the benefit previously observed from joint human-and-mouse training [15] extends to supervised training across a broader and more phylogenetically diverse set of species. We confirmed that this improved performance is not related to sequence homology between the added genomes and the held-out human windows (Supplementary Fig. A.1).

Together, these ablations show that genome pre-training and the expansion of supervised training beyond human and mouse both improve functional-track prediction, with the largest gains observed for gene expression. Both are also changes to the training recipe rather than to the architecture and therefore are complementary to existing sequence-to-function models rather than alternatives to them.

### 2.2. NTv3 matches AlphaGenome on human functional-track prediction

We next asked whether the phylogenetic breadth of NTv3 comes at a cost on densely characterized human functional tracks. We evaluated the model on AlphaGenome’s released human datasets and held-out evaluation split [6]. We fine-tuned NTv3 on all five assay tasks released at single-base resolution with available signal and outputs compatible with one-dimensional track prediction: ATAC-seq, DNase-seq, CAGE, PRO-cap and RNA-seq (Methods 4.5). NTv3 achieved higher performance than AlphaGenome on CAGE, PRO-cap and RNA-seq and remained within 0.023 Pearson correlation on ATAC-seq and DNase-seq (Fig. 3a; Supplementary Table C.8). The three improvements therefore occurred across all transcription-related modalities, consistent with the preferential gains in gene-expression prediction observed in our pre-training and multispecies-supervision ablations. These comparisons used five assay-specific NTv3 fine-tuning runs, since the model was already post-trained on thousands of human tracks, whereas the AlphaGenome values are those reported for its final jointly trained model, in which these assays formed part of the full multi-assay training corpus.

**Figure 3.**
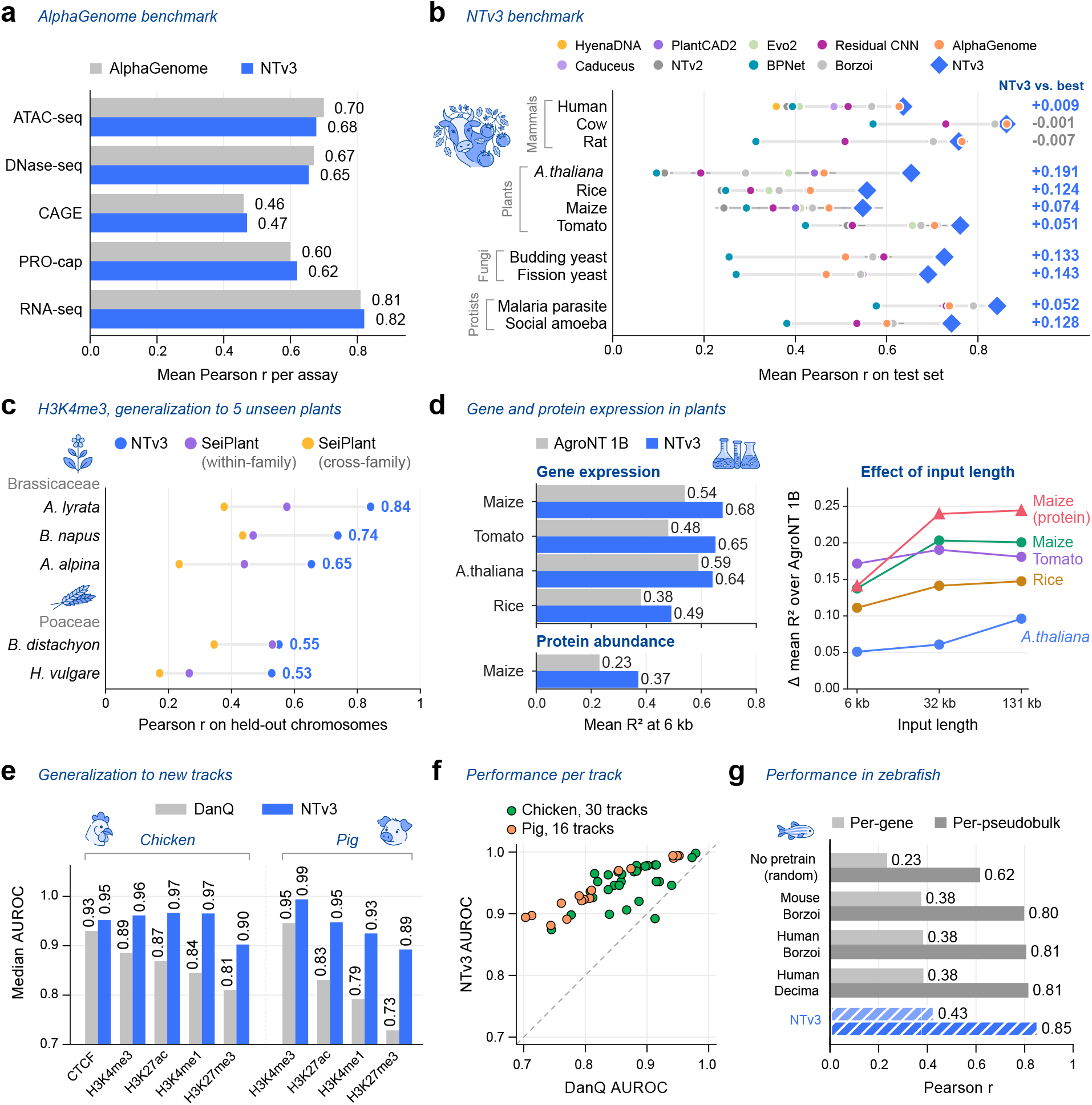
Benchmarking NTv3 across human and multispecies functional-track prediction tasks. **a** Mean Pearson correlation across human evaluation tracks for five AlphaGenome assays. AlphaGenome values are taken from its published evaluation. ATAC-seq and DNase-seq were evaluated on untransformed signal; CAGE, PRO-cap and RNA-seq after log1p transformation. **b** Test-set Pearson correlation across 11 species. Points and error bars show the mean and s.e.m. across three fine-tuning seeds for each model–species combination. Evo2 was evaluated with one seed due to compute limitations. Not every model was evaluated on every genome, and a model is drawn only on the species it has a run on: HyenaDNA and Caduceus on human only, PlantCAD2 on the four plants, and NTv2 and Evo2 on five of the eleven. BPNet, Residual CNN, Borzoi, AlphaGenome and NTv3 cover all eleven, so every row carries at least four published baselines and human carries eight. Values at right denote the difference between NTv3 and the strongest published baseline for each species. **c** H3K4me3 Pearson correlation on held-out chromosomes of five plant species absent from both NTv3 post-training and SeiPlant’s training set, scored against both released SeiPlant arms, within-family and cross-family. Both models are evaluated zero-shot. **d** Left, mean *R*^2^ across tissues at 6 kb input length for gene-expression prediction in four plant species and for protein abundance in maize. Right, the margin over AgroNT as the input context grows from 6 to 131 kb. **e** Median auROC per assay group for chicken and pig, compared with DeepFARM’s DanQ. **f** The same livestock tracks scored individually, NTv3 against DanQ, with the identity line shown. **g** Zebrafish gene-expression prediction on the DanioDecima benchmark: per-gene Pearson correlation across 304 pseudobulks, and per-pseudobulk Pearson correlation across genes. NTv3 is evaluated zero-shot in panels **c, e** and **f**, and after task-specific fine-tuning in panels **a, b, d** and **g**.

For completeness, we independently evaluated NTv3 on Borzoi’s published benchmark [5]. In the resolution-matched comparison, NTv3 was fine-tuned on the same training data and evaluated at Borzoi’s native 32-bp resolution, improving mean Pearson correlation across every assay group by 1.2–3.2%, with the largest gain on RNA-seq (Supplementary Fig. A.2a,b,e; Supplementary Table C.7). In a complementary zero-shot comparison on 1-bp targets, NTv3 also outperformed Borzoi across all assay groups (Supplementary Fig. A.2a–c). Together, these benchmarks show that NTv3 reaches AlphaGenome-level performance on matched human assays while consistently outperforming Borzoi on its own benchmark.

### 2.3. NTv3 accurately predicts functional tracks across 19 species

To evaluate cross-species transfer under standardized conditions, we developed the NTv3 Benchmark, a suite of 155 functional-track prediction tasks spanning 11 species, on which we compared convolutional models trained from scratch, pre-trained genomic foundation models and sequence-to-function models such as Borzoi and AlphaGenome (Methods 4.6). To compare model families at a common context length, every model receives the same 32-kb input, a length that all included models can process, together with the same task-specific 1-bp prediction head and per-nucleotide Poisson–multinomial loss (Supplementary Fig. A.3a). We note that all models were evaluated under a shared fine-tuning protocol to allow direct comparisons, but further model-specific tuning could improve performance and change the relative ranking. The benchmark evaluates two forms of generalization. For human, *Arabidopsis thaliana*, rice and maize, which were represented during NTv3 post-training, we selected assays and tracks absent from the post-training dataset. The remaining mammalian, plant, fungal and protist genomes were entirely absent from post-training, enabling to evaluate species-level transfer.

Across three independent fine-tuning runs per model and species, NTv3 achieved the highest performance of any published model, including AlphaGenome, on nine of the eleven species (Fig. 3b, Supplementary Fig. A.3; Supplementary Table C.4). The two exceptions were the two non-human mammals in the panel (cattle and rat), where AlphaGenome was 0.001 and 0.007 Pearson correlation ahead respectively, although both differences were within one standard error. This result is consistent with AlphaGenome showing its strongest cross-species performance in mammals closely related to its training species. Compared with the other eight models, NTv3 exceeded the strongest on all species, and across the eight plant and microbial genomes its margin over the strongest published baseline ranged from 0.05 to 0.19. Averaged over the eleven species, NTv3 reached a mean Pearson correlation of 0.71, against 0.60 for AlphaGenome and 0.58 for Borzoi, with the gains holding for both assays and species absent from NTv3 post-training.

This generalization extended to existing published plant benchmarks. In a zero-shot evaluation of H3K4me3 prediction across five species absent from NTv3 post-training, NTv3 outperformed both released SeiPlant models in every species (Fig. 3c; Supplementary Table C.9) [36]. We also fine-tuned NTv3 on the gene-expression and protein-abundance tasks introduced with AgroNT [37], in which a gene’s promoter-proximal sequence is used as input to predict the respective gene or protein levels. At AgroNT’s original 6-kb input length, NTv3 achieved higher mean *R*^2^ across all tasks, with absolute improvements ranging from 0.05 to 0.17. Increasing the NTv3 input context to 131 kb produced further gains (Fig. 3d; Supplementary Table C.9). Because inputs are centered on the transcription start site, longer windows add potential distal regulatory elements but also more of the transcribed gene body, neighboring genes and the overall composition of the region, such as its GC content.

We obtained similar results in livestock and zebrafish. On the DeepFARM benchmark of CTCF binding and histone modifications, NTv3 zero-shot increased median auROC (area under the receiver operating characteristic curve) relative to DanQ [38] across all nine chicken and pig assay groups, with improvements ranging from 0.022 to 0.164 (Fig. 3e; Supplementary Table C.9) [39]. In addition, NTv3 outperformed DanQ on 45 of the 46 individual functional tracks (Fig. 3f). On the DanioDecima benchmark, fine-tuned NTv3 scored highest for zebrafish gene-expression prediction among models initialized from human- and mouse-trained Borzoi and Decima checkpoints (Fig. 3g; Supplementary Table C.9) [40].

Across the NTv3 Benchmark and additional external evaluations, NTv3 achieved the highest performance among the evaluated models in 19 species spanning mammals, birds, fish, plants, fungi and protists. These results establish NTv3 as a broadly transferable model for functional-track prediction in both zero-shot and fine-tuned settings.

### 2.4. NTv3 leverages functional-track representations to improve *ab initio* gene prediction

We next asked whether representations learned through multispecies functional-track prediction transfer to structural genome annotation. Traditional *ab initio* gene predictors learn gene structure from genomic sequence and reference annotations, either through probabilistic models such as AUGUSTUS [26, 41] or through deep neural networks coupled to structured decoding, as in Helixer [27, 42] and Tiberius [28, 29]. Pre-trained sequence representations have also been applied to nucleotide-level genomic annotation, including SegmentNT and GeneCAD [32]. By contrast, NTv3 has additionally been trained to predict experimentally measured functional tracks across multiple assays and phylogenetically diverse species. We therefore fine-tuned its representation to produce complete coding gene models. To our knowledge, existing *ab initio* gene predictors have not been initialized from a representation trained on experimental functional-genomics tracks. The distinction concerns the learned representation rather than the inference inputs: like the other *ab initio* predictors evaluated here, NTv3 predicts from DNA sequence alone and does not receive functional tracks at inference.

We trained separate annotation models for mammals, plants and insects (three clades represented in functional-track post-training) using 1-Mb genomic windows and clade-specific genome splits derived from those of Tiberius (Methods 4.4, Supplementary Table C.2) [28, 29]. An HMM decoder converts per-base posteriors over intergenic, intronic, coding and splice-site states, together with coding phase, into complete coding transcripts. Joint prediction of gene states and coding phase distinguishes this task from nucleotide-level segmentation approaches such as SegmentNT [30]. We evaluated predictions on held-out species at the CDS-exon and gene levels, requiring exact coordinate matches to the longest annotated isoform of each gene and excluding untranslated regions, and we report precision, recall and F1 throughout (Methods 4.4.3, Supplementary Fig. A.8, Supplementary Table C.3). We also provide results using all annotated isoforms in Supplementary Fig. A.6a.

NTv3 achieved higher CDS-exon and gene-level F1 than competitors Helixer and Tiberius in all three clades (Fig. 4a). At gene level, the mean F1 scores for Helixer, Tiberius and NTv3 were respectively 28.7%, 52.3% and 56.4% across two held-out mammals; 55.8%, 67.0% and 73.4% across five plants; and 34.3%, 50.6% and 60.7% across ten insects. NTv3 had the highest gene-level F1 of all callers in each of the 17 test species (two-sided sign test against Tiberius, *P* = 1.5 × 10^−5^; Methods 4.4.3). On plants, we additionally compared with the plant-specialist model GeneCAD [32]. GeneCAD led Tiberius at gene level on this clade, with a mean F1 of 68.9% against 67.0%, but trailed it at CDS-exon level, 86.3% against 88.1%. Still, NTv3 exceeded GeneCAD in all five plant species at both levels. The improvement in insects differed markedly between evaluation levels: NTv3 exceeded Tiberius by 10.1 percentage points at gene level and performed better in all ten species, whereas their mean CDS-exon-level gap was far smaller, 86.2% against 85.1%, with Tiberius ahead in three of the ten (*Zerene cesonia, Colias croceus* and *Danaus plexippus*) (Fig. 4a). In this clade NTv3 was trained on 93 of the 100 assemblies that Tiberius itself was trained on, with the same BRAKER annotations, so its margin there does not depend on a larger or differently labeled training set. The gene-level advantage held in all three clades under either the longest- or the any-isoform matching rule (Supplementary Fig. A.6a). The predicted protein sets from NTv3 also had higher BUSCO completeness than those from Tiberius, recovering 98.1% versus 97.0% of BUSCO markers in cow and 96.0% versus 95.6% in beluga whale (Fig. 4b) [43]. Both predictors remain below the unfiltered reference proteome, which recovers 98.9% in cow and 97.7% in beluga whale, so the interpretable quantity is each caller’s shortfall against that ceiling rather than an absolute completeness. Further evaluations on protein-identity recovery, open-reading-frame validity and intron-chain matching are given in Supplementary Fig. A.7.

**Figure 4.**
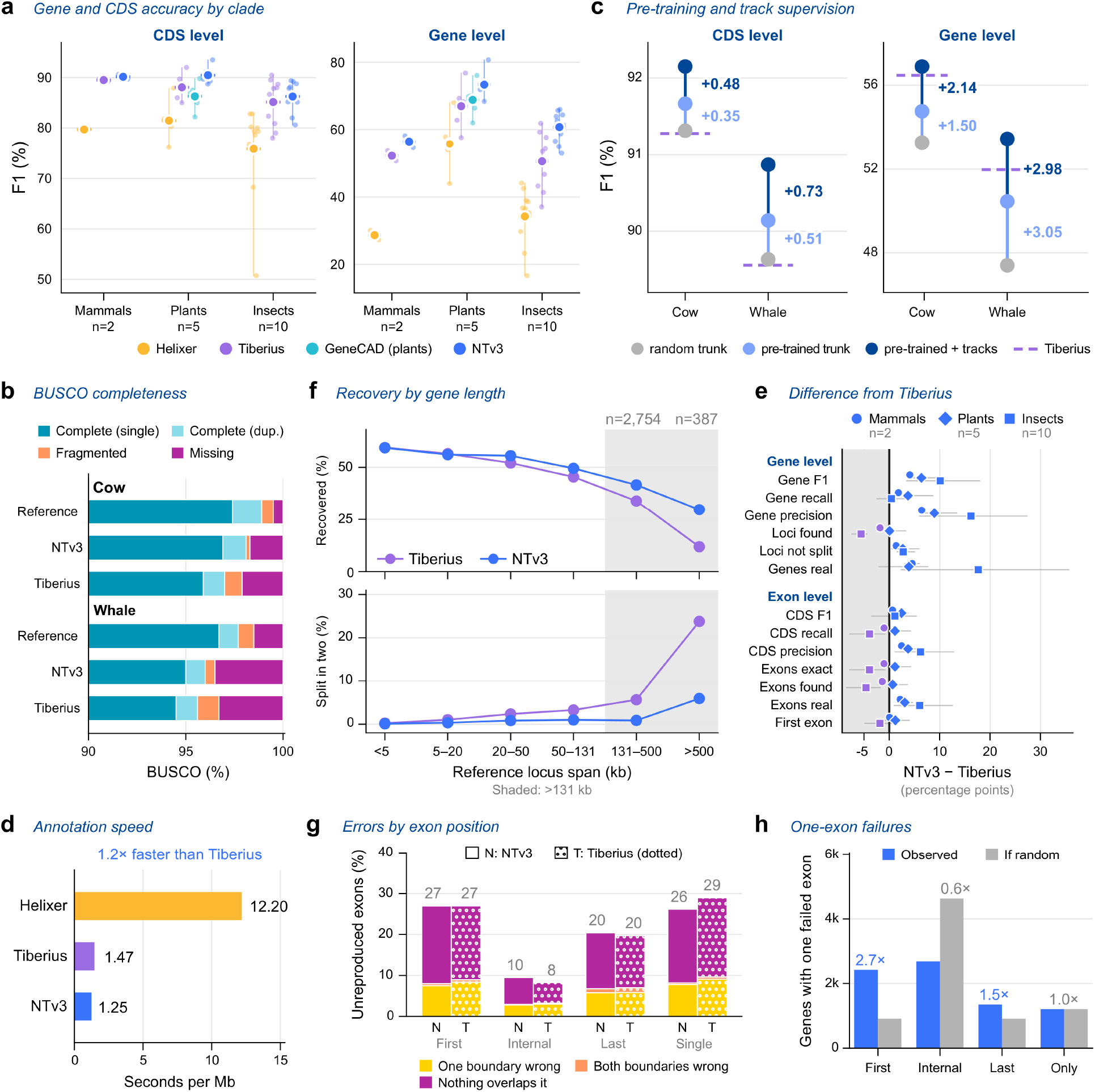
Evaluation of NTv3 for *ab initio* gene prediction across mammals, plants and insects. **a** CDS- and gene-level exact-match F1 by clade for each caller. Small points denote individual species, large filled circles denote clade means and vertical lines span the species within each clade. GeneCAD is shown only for plants. **b** BUSCO completeness of predicted proteomes for the two held-out mammals, compared with the corresponding unfiltered reference proteomes. Bars partition BUSCO markers into complete single-copy, complete duplicated, fragmented and missing categories. **c** The separate contributions of genome pre-training and of functional-track post-training, per held-out genome, from the 131-kb initialization ablation, scored on matched regions (the five largest cow chromosomes and the five largest beluga whale scaffolds). The values are not comparable with (**a**) (Methods 4.4). The dashed line is Tiberius on the same regions. **d** Wall-clock annotation time per Mb on one H100 80-GB GPU, measured end to end for each tool as its authors ship it, unweighted mean over the 13 benchmark assemblies whose mean contig exceeds the 1-Mb decoding window. GeneCAD is measured but not shown, being plant-specific and the slowest of the four. **e** Each structural aspect as a signed difference, NTv3 minus Tiberius, oriented so that right is always better, summarized to one marker per clade with the range across its species behind it. **f** Loci recovered exactly, and loci split into two or more predictions, against the span of the reference locus, pooled across the two mammals. Counts are summed and rates re-derived rather than averaging percentages, so every bin is weighted by the genes it contains. The shaded band marks spans above 131 kb. **g** Exon outcome by position within the transcript, for internal, last and first coding exons and for single-exon genes, under the longest-isoform rule. **h** Among genes that fail the exact-match criterion by exactly one exon, which exon position was incorrect, relative to a uniform expectation.

To distinguish the contribution of learned representations from that of architecture and sequence length, we trained otherwise identical annotation models initialized randomly, from genome pre-trained or from tracks post-trained NTv3 models. Relative to random initialization, genome pre-training increased CDS-exon-level F1 by 0.4 and 0.5 percentage points in cow and beluga whale, respectively, and gene-level F1 by 1.5 and 3.1 points. Initialization from the tracks post-trained model produced further gains of 0.5 and 0.7 points at CDS-exon level and 2.1 and 3.0 points at gene level (Fig. 4c). At gene level, our primary metric, only the model combining genome pre-training and functional-track post-training exceeded Tiberius in both species.

NTv3 was also among the fastest of the evaluated methods. Every caller was timed end to end on a single NVIDIA H100 80-GB GPU using the respective default settings. On the 13 benchmark assemblies whose mean contig exceeds the 1-Mb window, NTv3 ran at 1.25 s/Mb against 1.47 s/Mb for Tiberius and 12.20 s/Mb for Helixer, a 1.2-fold advantage over Tiberius and a 9.8-fold advantage over Helixer (Fig. 4d). Because each contig is padded to whole 1-Mb windows, NTv3’s cost rises on fragmented assemblies, and on the remaining four Tiberius was faster, at 1.91 against 2.54 s/Mb for NTv3 (Supplementary Table C.3). GeneCAD annotates plants only, so it was timed on the five plant assemblies rather than on this set, achieving 39.3 s/Mb, the slowest of the four callers. Thus, among the evaluated *ab initio* predictors, NTv3 achieved the highest accuracy across mammals, plants and insects, at a runtime comparable to Tiberius and about tenfold faster than Helixer.

### 2.5. NTv3 improves precision and long-gene recovery but remains limited at first exons

To identify the sources of the improvement over Tiberius, we compared the two models across 13 measures of gene and exon structure in all three clades (Fig. 4e, Supplementary Fig. A.8, Supplementary Fig. A.9) [28, 29]. The main difference was increased precision, with NTv3 improving gene-level precision by 6.4 percentage points in mammals, 8.9 points in plants and 16.2 points in insects. The fraction of predicted genes overlapping a reference locus increased by 4.5, 3.9 and 17.6 points, respectively. Recall gains were smaller and not uniform: gene-level recall rose by 1.8 points in mammals and 3.7 in plants but by only 0.4 in insects, and several recall-type measures moved against NTv3. The fraction of reference loci overlapped by any prediction fell by 1.8 points in mammals and 5.6 in insects, where the fractions of recovered exons and coding sequences also fell, by 3.9 points each. This follows from how many genes each caller predicts: NTv3 predicted close to the reference gene count in every clade (0.94, 0.98 and 1.03 predicted genes per reference locus in mammals, plants and insects), whereas Tiberius over-predicted, most strongly in insects (1.02, 1.06 and 1.45, up to 1.99 in *Leptidea sinapis*). Tiberius’s additional models overlap more reference loci without reproducing them, and split more of them. The 10.1-point improvement in insect gene-level F1 therefore reflected substantially higher precision despite little change in exact-match gene recall and lower locus-, exon- and coding-sequence recovery, indicating that NTv3 produces fewer false-positive gene models. Results for every reference locus in all three clades are given in Supplementary Fig. A.10.

The improvement also varied with gene length. NTv3 recovered more reference genes and fragmented fewer loci than Tiberius as gene length increased, with the difference becoming most apparent above 131 kb for mammals (Fig. 4f, Supplementary Fig. A.9b, Supplementary Fig. A.11). Such loci are a minority, 3,141 of the 39,966 mammalian reference loci (7.9%; Fig. 4f, Supplementary Fig. A.8), so the long-gene advantage adds to, rather than explains, the gene-level margin, which comes mainly from precision (Fig. 4e). Most plant and insect species showed the same length dependence, at the shorter spans typical of their genes (Supplementary Fig. A.9b). Among the 387 reference loci longer than 500 kb (200 in cow and 187 in beluga whale), NTv3 recovered 33.0% against 11.5% for Tiberius in cow and 26.2% against 12.3% in beluga whale, and fragmented 6.0% against 28.0% and 5.9% against 19.3% into two or more predictions. Fragmentation is insensitive to the matching rule: across all reference loci, recomputed against the best-matching annotated isoform, NTv3 fragments 85 loci against Tiberius’s 323 in cow and 93 against 311 in beluga whale, 3.8-fold and 3.3-fold fewer, close to the 3.7-fold and 3.2-fold advantages measured genome-wide under the longest-isoform rule. Both predictors tile the genome rather than decoding it in one pass, but they tile it differently. Tiberius partitions the sequence into seamless 500,004 bp tiles and decodes each in a single Viterbi parse; where a tile terminates inside a gene, it re-decodes the two adjacent tiles concatenated and keeps the longer coding structure. NTv3 instead decodes overlapping 1-Mb windows and retains the central 75% of each (786,432 bp), so that consecutive decoded regions abut while every emitted base carries at least 131 kb of flanking context. Its decoded span per parse is therefore about 1.6 times longer, giving roughly a third fewer decode boundaries per genome, although unlike Tiberius it does not re-decode across them. The reduced fragmentation is consistent with this longer decode span and the larger context available at every emitted position, although a context-length ablation would be required to establish that explanation directly.

We next examined errors according to exon position, pooling the two held-out mammals. For NTv3, internal coding exons had an error rate of 9.5%, compared with 20.4% for last coding exons, 26.9% for first coding exons and 26.2% for single-exon genes (Fig. 4g, Supplementary Fig. A.13c). NTv3 reduced the error rate for single-exon genes from 28.9% with Tiberius to 26.2%, whereas on first coding exons, the least accurately predicted class of multi-exon genes for both callers, the two were level (26.9% for NTv3 and 27.0% for Tiberius). Among failed gene models with at least one incorrect coding exon, 43.2% differed from the reference by exactly one exon (45.3% in cow and 40.9% in beluga whale; Supplementary Fig. A.12c, Supplementary Fig. A.13b). Among these one-exon errors, the incorrect exon was enriched 2.7-fold at the first coding position relative to its expected frequency, whereas internal exons were depleted to 0.6-fold (Fig. 4h, Supplementary Fig. A.13). Both patterns hold when any annotated isoform is accepted as a match (Methods 4.4.3).

Together, these analyses show that the gains from NTv3 arise primarily from fewer false-positive gene models and reduced fragmentation of long loci, whereas, among multi-exon genes, the first coding exon remains the class both callers identify least reliably, and the position most over-represented among the genes they fail. We release the NTv3 model checkpoints together with an end-to-end inference pipeline, enabling users to apply the corresponding clade-specific model to unannotated genomes.

### 2.6. NTv3 enables multiscale interpretation of regulatory genomic features

We investigated whether NTv3’s functional representations could support regulatory interpretation across sequence scales. We combined assay-specific functional-track predictions, base-resolution gradient attributions, transformer attention maps and controlled sequence perturbations (Fig. 5a; Methods 4.8.1). These complementary post-hoc readouts respectively describe predicted regulatory activity, the local sequence features contributing to a selected prediction and model dependencies across longer genomic distances. To make these capabilities accessible as a unified analysis workflow, we additionally developed an LLM-guided interpretation pipeline that leverages NTv3 track prediction, gene annotation, perturbation, attribution and locus-context tools to examine genomic loci (Methods 4.8.3; Supplementary Fig. A.20).

**Figure 5.**
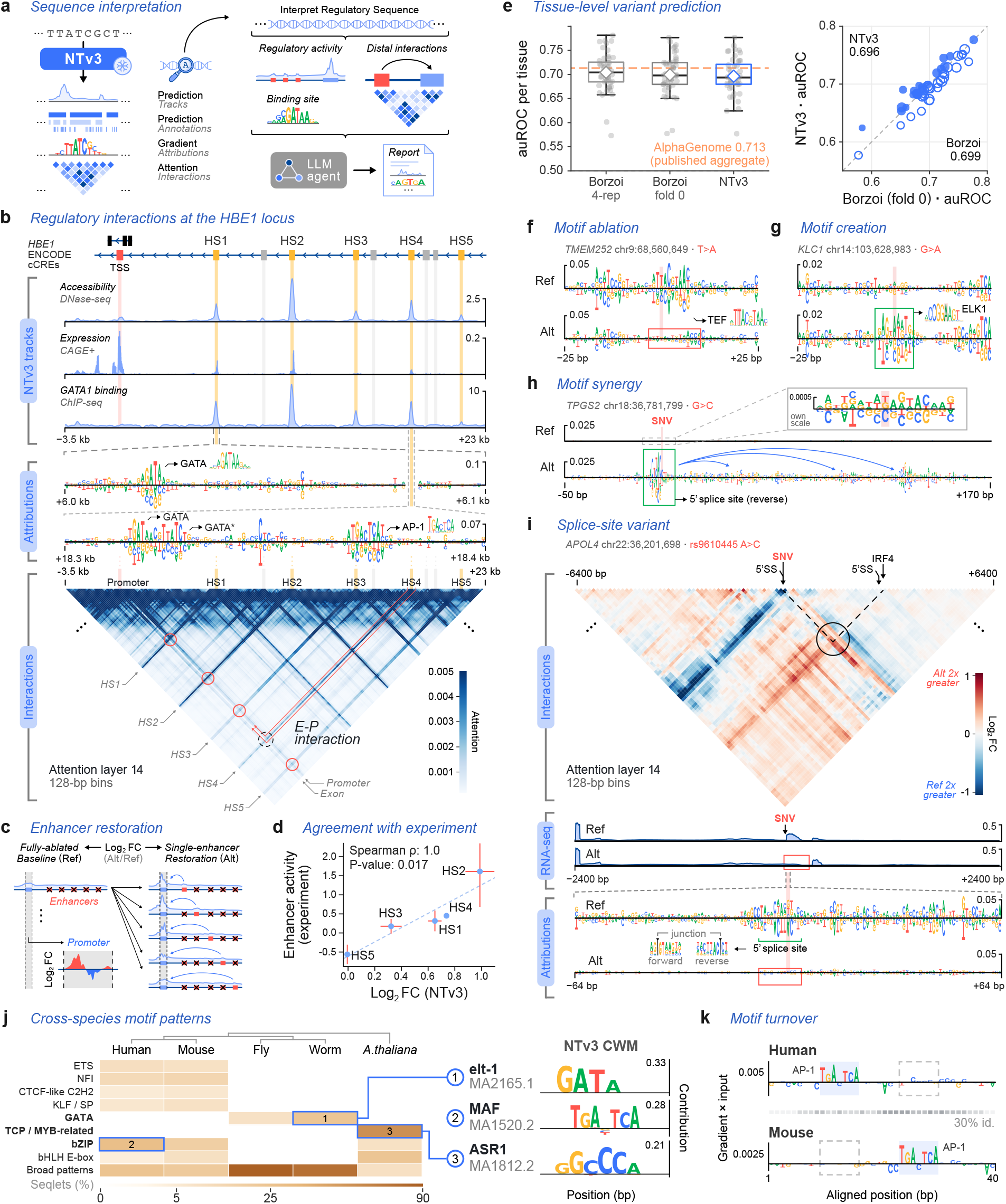
NTv3 enables multiscale interpretation of regulatory features across loci, variants and species. Functional-track predictions, gradient-based attribution maps, attention maps and targeted sequence perturbations provide complementary views of regulatory activity from local sequence features to long-range interactions. **a** Schematic of complementary NTv3 readouts for regulatory-sequence interpretation, including functional-track predictions, gradient-based attributions and long-range attention interactions. **b** Analysis of the human *HBE1* locus (−3.5 kb to +23 kb from the TSS). ENCODE candidate cis-regulatory elements provide reference context; K562 DNase-seq, CAGE and GATA1 ChIP-seq tracks localize regulatory activity, attribution maps identify sequence features, and layer-14 attention highlights promoter interactions with enhancers HS1–HS5. **c, d** *In silico* enhancer-restoration strategy (**c**) and comparison of predicted K562 CAGE effects with measured enhancer activity (**d**; Spearman *ρ* = 1.0, *P* = 0.017). **e** Quantitative eQTL variant-effect benchmark across 48 GTEx tissues. Left, per-tissue auROC distributions for NTv3, Borzoi fold 0 and the four-replicate Borzoi ensemble; white diamonds denote the unweighted mean per-tissue auROC and the dashed orange line denotes the published AlphaGenome aggregate. Right, per-tissue auROC for NTv3 versus Borzoi fold 0, with point area scaling with variant count. **f–h** Representative allele-specific responses illustrating motif ablation (**f**; TEF, MA0843.2), motif creation (**g**; ELK1, MA0028.2), and a splice-associated response at a *TPGS2* variant (**h**; chr18:36,781,799, G>C), with the latter showing coordinated attribution changes around a reverse-strand 5^*′*^ splice site together with altered long-range interactions. **i** Reconstruction of the HIGH-impact *APOL4* splice variant rs9610445, showing disruption of splice-donor attribution together with reduced predicted RNA-seq signal. **j** Representative TF-MoDISco motif families recovered from independently defined native-enhancer cohorts in human, mouse, fly, worm and *Arabidopsis* [44], with contribution weight matrices matched to one pooled JASPAR 2026 collection (vertebrates, insects, nematodes and plants) [45] and grouped by DNA-binding domain. **k** Attribution on one published human–mouse enhancer ortholog [46] localizes an AP-1 motif (MA0099.2) in both sequences at different aligned positions. Where shown, attention maps are from layer 14 in 128-bp bins.

For illustration, we examined the well-characterized human *HBE1* locus, where five DNase hypersensitive enhancers (HS1–HS5) have experimentally measured activities [14, 47]. Functional-track predictions and sequence attributions localized the promoter and hypersensitive elements, while long-range attention high-lighted the expected dependencies between promoter-proximal and enhancer-containing regions (Fig. 5b; Methods 4.8.2). In a complementary perturbation analysis, restoring each enhancer individually from a common shuffled background produced the predicted hierarchy HS2 > HS4 > HS1 > HS3 > HS5, the same order as the measured hierarchy (Spearman *ρ* = 1.0, *P* = 0.017; Fig. 5c,d).

To test whether this signal extended beyond a single locus, we next evaluated enhancer–gene linking on the independent ENCODE-rE2G K562 CRISPR benchmark (Methods 4.8.2). NTv3 and Borzoi produced similar auPRCs (area under the precision–recall curve) for pairs separated by less than 10 kb (all/matched pairs: 0.86/0.87 versus 0.89/0.89) and 10–100 kb (0.53/0.55 versus 0.59/0.60), whereas NTv3 was numerically higher in the 100–2,500 kb bin (0.24/0.41 versus 0.21/0.37), which contained 82 positive pairs (42 matched). Published AlphaGenome results were higher in all three bins, with auPRCs of 0.91/0.91, 0.77/0.78 and 0.35/0.64, respectively. The complete comparison, including precision–recall curves, auROC, precision at 70% recall and a reciprocal-distance baseline, is reported in Supplementary Fig. A.21 and Supplementary Note B.7 [5, 6, 48]. Thus, NTv3 recovers enhancer–gene regulatory signal across genomic distances, although it does not outperform the strongest model on this benchmark.

### 2.7. NTv3 reveals regulatory sequence features across species

The multispecies training of NTv3 brings genomes from diverse phylogenetic groups into a common model, providing an opportunity to interpret local regulatory sequence features across species. We first asked whether the model captures the effects of sequence perturbations in human, where genetic variants and their functional consequences are comparatively well characterized. On the established benchmark of 21,241 eQTLs from 48 GTEx tissues [6, 49], per-tissue auROC for NTv3 ranged from 0.577 to 0.777, while tissue-weighted mean auROC was 0.701 for NTv3, 0.704 for Borzoi fold 0 and 0.709 for the four-replicate Borzoi ensemble (Fig. 5e; Supplementary Table C.10). The corresponding tissue-weighted auPRCs were 0.710, 0.704 and 0.712. Relative to Borzoi fold 0, NTv3 did not differ significantly by auROC (Δ = −0.0029, *P* = 0.20) and was modestly higher by auPRC (Δ = +0.0062, *P* = 0.038); relative to the four-replicate ensemble, NTv3 had lower auROC (Δ = −0.0075) but no significant difference in auPRC (Δ = −0.0025, *P* = 0.27). AlphaGenome reports an aggregate auROC of 0.713 for this benchmark, although per-tissue results and eQTL auPRC were not available. Thus, although NTv3 does not establish an advantage over existing variant-effect models, these results show that its functional representation retains regulatory-variant signal.

We next examined what regulatory features underlie these responses by contrasting reference and alternative alleles using NTv3’s complementary interpretable readouts (Fig. 5f–i; Methods 4.8.2). Representative variants illustrate three qualitatively distinct responses: motif ablation, motif creation and a splice-associated response. The first two localized effects to altered TF-binding grammar, whereas the third, at a *TPGS2* variant near a reverse-strand 5^′^ splice site, was associated with coordinated attribution changes together with altered long-range interactions (Fig. 5f–h). At rs9610445 in *APOL4*, the alternative allele sharply attenuated splice-donor attribution at the variant and reduced predicted RNA-seq signal by approximately 5-fold (Fig. 5i). Together, the loss of splice-donor attribution and the concordant reduction in predicted expression provide a coherent model-based mechanism for a variant classified as HIGH-impact across 15 *APOL4* transcripts [50]. These examples illustrate how complementary model readouts can generate sequence-level hypotheses about predicted variant effects without, by themselves, establishing causal mechanisms.

Comparable perturbation datasets are substantially more limited across many of the other species represented by NTv3. We therefore asked whether the same attribution framework recovers recognizable regulatory sequence features across species. We analyzed independently defined native-enhancer cohorts from human, mouse, fly, worm and *Arabidopsis*, rather than orthologous elements, and computed accessibility-directed attribution maps using the corresponding species-conditioned NTv3 outputs. TF-MoDISco [44] patterns were matched with TOMTOM-lite [51, 52] against one pooled JASPAR 2026 collection spanning vertebrates, insects, nematodes and plants [45], so that motif labels were not restricted by species, and grouped by DNA-binding domain. Representative matches included a GATA motif in worm (elt-1, MA2165.1), a Maf-related MAF motif in human (MA1520.2), and an *Arabidopsis*-derived pattern matching the plant MYB-related ASR1 motif (MA1812.2; Fig. 5j; Methods 4.8.2). These results show that base-resolution accessibility attributions recover recognizable enhancer-associated motif families across phylogenetically diverse genomes.

As a complementary example, we examined one published human–mouse enhancer ortholog with each sequence conditioned on its own species token [46]. NTv3 localized an AP-1 motif match in both sequences, but at different aligned positions (Fig. 5k; Methods 4.8.2). This example illustrates how attribution maps can identify a shared motif class despite positional turnover within an orthologous enhancer pair. Together, these analyses show that NTv3 supports regulatory interpretation from individual nucleotides to distal enhancer–gene relationships and across phylogenetically diverse genomes.

### 2.8. NTv3 enables conditional regulatory sequence design

Finally, we tested whether the representations learned by NTv3 through genome pre-training and functional-track prediction could support regulatory-sequence design. Starting from NTv3 weights, we adapted the model to masked diffusion language modeling (MDLM) [34]. Whereas conventional masked language modeling uses a fixed masking rate, MDLM uses variable noise levels, training the model to recover the original bases across a range of corruption levels. At inference, the model generates sequences through iterative bidirectional denoising, starting from a span of all masked tokens that are gradually filled in by the model. Unlike the left-to-right decoding used by autoregressive genomic models [21, 25], our MDLM adaptation of NTv3 permits both *de novo* generation and infilling between fixed flanking sequences.

The MDLM adaptation retains the NTv3 backbone and nucleotide-prediction head. We provided functional conditions through adaptive layer normalization and used classifier-free guidance to steer sampling toward the requested activity [53, 54]. The masked-diffusion training, iterative sampling and guidance procedure are illustrated in detail for the *Drosophila* enhancer-design setting in Supplementary Fig. A.15 (Methods 4.9).

We evaluated conditional generation using three high-throughput reporter-assay datasets: plant promoters measured in maize protoplasts and tobacco leaves [55], plant enhancers measured under four conditions in tobacco leaves [56], and *D. melanogaster* S2-cell enhancers measured with two core promoters, the developmental *Drosophila* synthetic core promoter (DSCP) and the housekeeping RpS12 promoter [3, 57] (Fig. 6a). For each dataset, we fine-tuned NTv3 on regulatory sequences embedded in their reporter context. Measured activities were discretized into conditioning bins; for promoter-selective fly enhancer design, the condition jointly specified activities with the two core promoters. During sampling, the surrounding reporter sequence was held fixed and the regulatory element was generated by iterative denoising.

**Figure 6.**
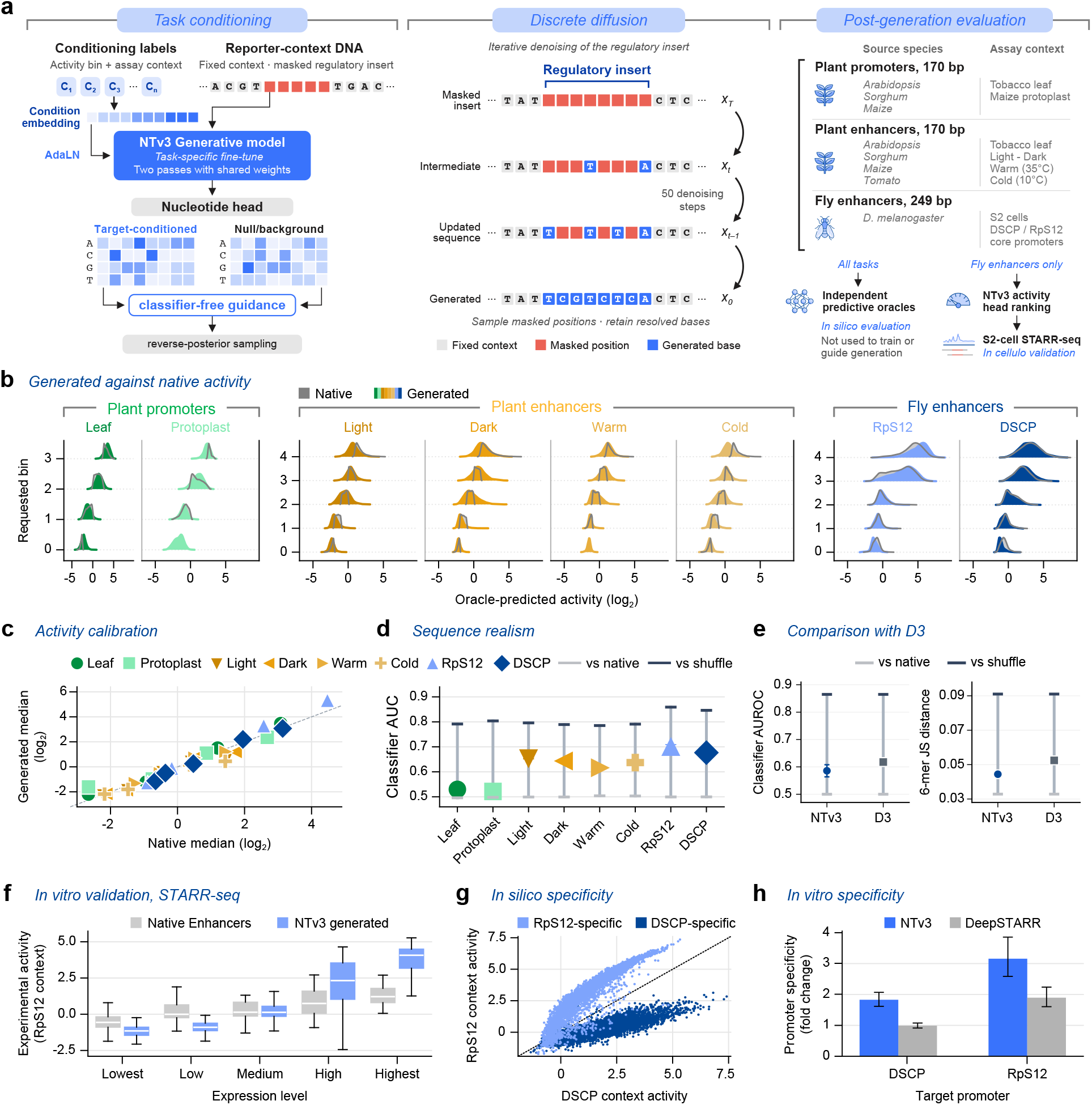
Conditional regulatory-sequence design with NTv3 and experimental validation in *Drosophila* S2 cells. **a** Conditional masked-diffusion sequence-generation framework. Separate NTv3 fine-tunes, each initialized from the same post-trained NTv3 checkpoint, generate inserts for plant promoter (170 bp), plant enhancer (170 bp) and *D. melanogaster* S2 enhancer (249 bp) reporter assays. The assay condition and requested activity bin condition iterative denoising of the insert while the plasmid backbone remains fixed. **b** Distributions of activity predicted by task-specific oracle models for native and generated sequences across eight assay contexts, stratified by requested activity bin. Native sequences are shown in gray and generated sequences in color. **c** Median oracle-predicted activity of generated sequences compared with native sequences from the corresponding activity bin. Each point represents one activity-bin–context combination; the dashed line denotes equal medians. **d** Classifier auROC for distinguishing generated from native sequences in each assay context. Reference ticks show classification between two disjoint subsets of native sequences and between native and dinucleotide-shuffled sequences. Values closer to the native-versus-native reference indicate greater similarity to native sequence. **e** Comparison with D3-Tran on the D3 fly-enhancer benchmark. Classifier auROC (left) and 6-mer Jensen–Shannon distance (right) are shown with native-versus-native and native-versus-dinucleotide-shuffled reference levels; lower values indicate greater similarity to native DNA. Both models were evaluated using the same held-out sequences and evaluators. The NTv3 arm is the draw matched to the native activity mix; raw metric values for both models are given in Supplementary Fig. A.18b. **f** STARR-seq activity measured in *Drosophila* S2 cells for 300 generated enhancers conditioned on five activity bins in the RpS12 promoter context, compared with native enhancers. Box plots summarize the sequence-level activity distributions. **g** Oracle-predicted activity of promoter-selective enhancer designs in the DSCP and RpS12 contexts. Each point represents one generated sequence; the diagonal denotes equal predicted activity in the two promoter contexts. **h** Experimentally measured promoter specificity of NTv3-generated and DeepSTARR-selected sequences in S2 cells. Specificity is defined as the activity fold change between the target and alternative promoter contexts. A total of 150 sequences from each design strategy were evaluated at a matched candidate-generation budget. Because a fold change is a ratio, bars show geometric means and error bars denote 95% confidence intervals computed in log space and exponentiated. Panels **b**–**e** and **g** report computational evaluations; panels **f** and **h** report STARR-seq measurements in S2 cells. We note that the experimentally tested fly sequences (**f**–**h**) were generated and selected using an earlier checkpoint from the same NTv3 model lineage; the revised checkpoint used elsewhere in this manuscript assigned comparable activity scores to these designs (see Methods 4.10).

We first assessed conditional fidelity using published task-specific predictive oracles that were not used to train or guide the generator. Across the three datasets, the oracle-predicted activities of generated sequences spanned the ranges observed for native sequences (Fig. 6b). More importantly, predicted activity shifted with the requested condition, separating designs from different activity bins rather than merely reproducing the overall native distribution (Fig. 6c, Supplementary Fig. A.16). Generated sequences also approximated the GC-content distributions of native sequences (Supplementary Fig. A.17) and BLASTN searches identified no training-set matches above the prespecified identity and alignment-length thresholds. These analyses establish conditional control in predictor space, which we experimentally validated for the fly enhancers below.

To assess sequence-distribution similarity beyond low-order composition, we trained binary classifiers to distinguish generated from native sequences (Methods 4.9.6). Classification between two disjoint subsets of native sequences provided a near-indistinguishable reference, whereas classification of native sequences against dinucleotide-shuffled controls provided a deliberately separable reference. Across all tasks, generated sequences were less distinguishable from native sequences than were the shuffled controls; for maize promoters, discrimination approached the native-versus-native reference (Fig. 6d, Supplementary Fig. A.18a). We also compared NTv3 with D3, a discrete diffusion model trained on the same fly enhancer dataset [58]. Using the identical held-out native set and the same evaluators, NTv3 designs showed lower generated-versus-native discrimination and a smaller 6-mer Jensen–Shannon distance than D3 designs (Fig. 6e, Supplementary Fig. A.18b).

#### 2.8.1. Experimental validation of generated fly enhancers

Fly enhancers provide a stringent test of conditional design because *Drosophila* enhancers are not equally compatible with all core promoters: developmental enhancers preferentially activate developmental core promoters such as DSCP, whereas housekeeping enhancers preferentially activate housekeeping core promoters such as that of the ribosomal-protein gene *RpS12* [59]. Because the two enhancer classes differ in their sequence motifs, designing an enhancer that activates one promoter but not the other requires a model to have learned promoter-specific regulatory rules, not only overall enhancer strength. To determine whether NTv3’s conditional control translated into activity in cells, we synthesized 600 fly enhancer designs spanning five requested activity bins, 300 in each of the two core-promoter contexts, and quantified their activity by STARR-seq in S2 cells [57]. Measured enhancer activity increased across the requested bins, producing the intended stratification of activity levels (Fig. 6f; Supplementary Fig. A.19). Thus, the conditional signal learned by NTv3 predicted experimentally measurable differences in enhancer activity.

We next tested whether NTv3 could design enhancers that preferentially activate one core promoter over another. We jointly conditioned generation on high activity with a target promoter and low activity with the alternative promoter. Before synthesis, the DSCP-targeted and RpS12-targeted designs separated in their predicted activities under the two promoter contexts (Fig. 6g). The two design sets also exhibited distinct motif-enrichment profiles consistent with established differences between developmental and housekeeping enhancer–core-promoter programs [59].

We compared direct conditional generation with an oracle-screening baseline under a matched computational budget. For the baseline, we generated a pool of random sequences and selected candidates using DeepSTARR predictions. We then assayed 150 NTv3-generated designs and 150 DeepSTARR-selected sequences with both core promoters. Promoter specificity was quantified as the ratio of activity with the target promoter to activity with the alternative promoter. Because this ratio is a fold change, we summarized it as a geometric mean. The NTv3 designs achieved target-to-alternative activity ratios of 1.83 (95% CI 1.62–2.07) for DSCP-targeted sequences and 3.16 (2.59–3.86) for RpS12-targeted sequences, exceeding the corresponding ratios obtained by oracle-based selection, whose DSCP-targeted arm did not differ from unity (0.99, 95% CI 0.91–1.08) (Fig. 6h). Under this budget-matched comparison, direct conditional generation therefore produced greater experimentally measured promoter selectivity than screening random sequences with a predictive oracle.

Together, these results show that the representations learned by NTv3 can be adapted through masked diffusion for conditional regulatory-sequence design. Computational analyses demonstrate control across plant and fly reporter assays, whereas experiments in S2 cells validate both graded enhancer activity and enhancer–core-promoter specificity. NTv3 thereby connects genome-scale representation learning and functional prediction with experimentally testable sequence design within a single model family.

## 3. Discussion

NTv3 shows that multispecies functional-track supervision can serve as a bridge between what a genomic foundation model learns from natural sequence and what it can subsequently predict, annotate and design, connecting comparative genomics and synthetic biology. Until now, genome-scale self-supervision, functional-track prediction, structural genome annotation and regulatory-sequence design have largely advanced through separate model families [5, 6, 19, 25, 29, 58]. In NTv3, self-supervision first supplies a phylogenetically broad sequence representation, post-training on measured transcriptional and chromatin activity grounds that representation in genome function, and task-specific adaptation converts the same trunk into a gene caller or conditional sequence generator.

The controlled ablations establish that both genome pre-training and phylogenetically diverse functional supervision contribute to track prediction. Although representations from pre-trained DNA language models have offered little advantage over conventional supervised models on some regulatory genomics tasks [60], here we demonstrate that pre-training can improve performance on both functional-track prediction and gene annotation. Extending post-training from human and mouse to 15 species improved expression prediction across species and in particular in human and mouse, extending earlier evidence for cross-species transfer beyond the conventional human–mouse setting [15]. Functional data from diverse species can therefore improve regulatory interpretation in well-characterized genomes as well as broaden the set of genomes that can be modeled. Such cross-species transfer is notable because regulatory elements themselves are poorly conserved: much of transcription-factor binding differs between human and mouse and across vertebrates, and enhancers turn over rapidly across mammals [61–63]. Yet in mice carrying human chromosome 21, transcription factors bind the human chromosome largely as they do in human liver [64], so what transfers is likely the sequence code that directs binding rather than the positions of individual sites, consistent with NTv3 locating a shared AP-1 motif at different positions in an orthologous human–mouse enhancer pair (Fig. 5k). Both interventions benefited transcriptional measurements more than accessibility, histone modifications or transcription-factor binding. This selectivity may reflect the predominance of RNA-seq in the post-training corpus or a closer correspondence between genome self-supervision and transcript structure. Balanced assay collections and factorial ablations will be required to distinguish these explanations.

This breadth did not require sacrificing performance in the best-characterized genome. On AlphaGenome’s five human tasks, NTv3 performed better on all three transcription-related modalities and remained close on chromatin accessibility; it also outperformed Borzoi across assay groups on Borzoi’s benchmark [5, 6]. Across standardized and external evaluations, NTv3 achieved the highest performance among the evaluated models in 19 species spanning mammals, birds, fish, plants, fungi and protists [36, 37, 39, 40]. The central advance is thus the combination of AlphaGenome-level performance on matched human assays with substantially broader transfer to model organisms, crops and sparsely characterized taxa, including species and assay tracks absent from functional post-training.

Functional supervision also improved structural genome annotation. Although NTv3 receives DNA alone at inference, its trunk has previously learned to reconstruct measured transcriptional and chromatin activity. Because RNA-seq coverage outlines exons, part of this transfer may amount to exon-level supervision. Post-training also includes close relatives of some test species (sheep for cow, wheat for *Brachypodium*, soybean for *Medicago*). Across held-out mammalian, plant and insect genomes, the resulting clade-specific NTv3 models outperformed Helixer and Tiberius at gene and CDS-exon level and outperformed the recent plant-specialist GeneCAD in all five evaluated plant genomes [27–29, 32, 42]. Genome pre-training and functional post-training contributed cumulatively, and at gene level only their combination exceeded Tiberius in both held-out mammals. The gains arose principally from higher precision and reduced fragmentation: for loci longer than 500 kb, NTv3 more than doubled exact recovery and reduced fragmentation three-to fivefold. These observations are consistent with its longer decoded span and greater flanking context, although a context-length ablation is needed to establish causality. The lower locus- and exon-level recovery observed in insects and the shared concentration of errors at first coding exons also identify clear targets for improvement.

The learned representation additionally supports regulatory hypotheses at multiple scales. At the *HBE1* locus, predicted tracks, attribution maps and *in silico* perturbations recovered an experimentally characterized enhancer hierarchy [14, 47], while attribution maps also recovered recognizable enhancer-associated motif families across taxa. Accurate track reconstruction did not by itself translate into better enhancer–gene ranking, and these analyses should be interpreted as hypothesis generation: attributions, attention and sequence perturbations do not establish mechanism without experimental intervention [65, 66].

Through discrete diffusion we demonstrate how NTv3 can be adapted from inference to regulatory-sequence design [34, 58]. Bidirectional denoising permits promoters or enhancers to be generated within fixed reporter contexts while conditioning on requested regulatory activity. Across plant and fly assays, the generated sequences followed the requested activity conditions and were more similar to native sequences than shuffled controls. Importantly, this workflow is validated not only with *in silico* oracles but also experimentally, grounding the generative capability in measurable regulatory outcomes. For fixed target activity and promoter context, NTv3 generates diverse families of sequence-distinct enhancers that satisfy the same functional constraints, enabling robust exploration of regulatory design space rather than reliance on a single solution. The combination of megabase-capable representations, supervised sequence-to-function heads, and experimentally validated conditional generation positions NTv3 as a general framework for design problems and suggests a path toward iterative, model-in-the-loop regulatory engineering that remains consistent with the predictive interfaces used for variant interpretation and functional annotation.

Together, these capabilities support complementary uses in comparative genomics and synthetic biology. Multispecies track prediction can prioritize regulatory activity in organisms with sparse functional data; sequence-only annotation can accelerate characterization of new assemblies and improve recovery of extended gene structures; and conditional generation can propose regulatory elements for specified assay and promoter contexts. This is especially relevant to model organisms and crops, where specialized foundation models have shown the value of taxonomically broader sequence learning but comprehensive functional measurements remain scarce [31, 37]. Because one model family covers animals and plants, these uses extend well beyond human and model-organism genomics. In agriculture, annotation and regulatory design in crops and farm animals could support breeding, including for changing climates; in biodiversity genomics, initiatives such as the Earth BioGenome Project aim to sequence all known eukaryotic species, most of which will have to be annotated and interpreted without functional data [67]; and in biotechnology, regulatory elements must often be designed for specific host organisms. We anticipate that NTv3 will serve as a scalable substrate for the next generation of genomic foundation models, enabling increasingly comprehensive, multispecies, long-range models as functional supervision, perturbation datasets, and generative objectives continue to mature.

### 3.1. Limitations and open questions

The NTv3 functional-track post-training corpus is imbalanced across taxa and assays, reflecting the broader concentration of functional-genomics data in a few well-characterized organisms. Human contributes substantially more tracks and cellular contexts than most species, and several genomes lack entire assay classes. This imbalance may constrain transfer to sparsely represented species, and it prevents us from fully separating the effects of phylogenetic breadth, training-data volume and assay composition. Similarly, the human-and-mouse versus 15-species ablation establishes the value of broader multispecies supervision as a whole but does not identify which species or clades drive the improvement. We therefore cannot determine whether human prediction benefits more from adding closely related primates and other mammals or from more distantly related species, nor what taxonomic mixture would be optimal. Resolving this question will require controlled ablations that match data volume and assay composition while adding or removing individual species or clades. Finally, we bounded the effect of cross-species homology on the held-out human comparison and found no detectable association (Supplementary Fig. A.1), but this analysis tests a homology gradient rather than a strict zero-homology holdout. Residual homology is intrinsic to multispecies genomic data and evaluations. We therefore regard performance on unseen species and unseen tracks as the strongest evidence for multispecies generalization.

The downstream interpretation benchmarks reveal a gap between functional-track prediction and regulatory prioritization. The gains achieved by NTv3 in track prediction did not translate into improved performance on GTEx eQTL classification or ENCODE-rE2G enhancer–gene ranking. On the GTEx benchmark, NTv3 was comparable to a single-fold Borzoi model but did not outperform the four-replicate Borzoi ensemble. On ENCODE-rE2G, NTv3 was comparable to Borzoi overall but performed below AlphaGenome, with the largest differences at intermediate and distal distances. The distal bins also contain relatively few experimentally supported positive pairs, limiting the certainty of the distance-stratified estimates. We therefore interpret these analyses as evidence that the NTv3 representation retains information relevant to variant and enhancer–gene regulation, rather than as demonstrating improved downstream prioritization. Closing this gap may require task-specific objectives and direct supervision for variant effects or enhancer–gene relationships.

The annotation results do not yet establish a universal gene predictor. Separate models are still required for mammals, plants and insects, and other major clades remain untested. Annotation throughput also depends on assembly contiguity: every contig is padded to a whole 1-Mb window, so NTv3 is fastest on chromosome-scale assemblies and loses its advantage on fragmented ones with many short scaffolds. The benchmark is also restricted to sequence-only *ab initio* methods and does not compare with pipelines such as BRAKER3 or GeneMark-ETP that receive RNA or protein evidence at inference [68]. Future work should determine whether broader clade training can yield a single annotation model without eroding taxon-specific accuracy, and whether functional representations complement rather than duplicate extrinsic transcript and protein evidence.

Experimental validation of generation was confined to fly enhancers in a plasmid reporter assay. The plant results demonstrate conditional control according to independent predictive oracles, not experimentally established activity, and some generated pools remain distinguishable from native elements. Oracle evaluation can reward evaluator-specific biases, while reporter activity may not transfer to an endogenous chromosomal context. Likewise, a 1-Mb receptive field and plausible attribution maps do not by themselves show that distal regulation is used correctly. Broader synthesis experiments, genomic integration and perturbational measurements will be needed to test design and interpretation across taxa, cell states and chromatin contexts.

Perturbational, single-cell, multiomic and population-scale data offer a path to more context-specific supervision [48, 69, 70], while functional measurements from additional species should test how far the observed transfer continues across phylogenetic distance. Integrating diffusion objectives earlier in training may also bring representation learning and generation under a more unified objective. By making functional-track supervision the bridge between phylogenetic-scale sequence learning and downstream biological tasks, NTv3 provides a route toward genomic models that learn from well-characterized organisms and transfer that knowledge to the long tail of species that remain functionally unexplored.

## Methods

## 4. Methods

### 4.1 NTv3 Models

NTv3 is a family of long-context genomic sequence models that share a U-Net-inspired backbone and are adapted to prediction and generation tasks through task-specific heads and objectives. The backbone accepts base-resolution DNA sequences of up to 2^20^ = 1,048,576 nucleotides and returns a representation at the same resolution. All NTv3 models derive from a model pre-trained by masked language modeling (MLM) on genomic sequences from OpenGenome2. We subsequently post-trained it on multispecies functional genomic tracks using a single output head shared across species, with species conditioning supplied only to that head. We further fine-tuned the NTv3 backbone for structured gene annotation as well as for conditional sequence generation by masked diffusion. The functional-track, annotation and generative models share a backbone and checkpoint lineage but are separate task-adapted models rather than a single jointly trained multitask checkpoint. We therefore write NTv3 for the family throughout, and name the training stage in words wherever a statement depends on which one is meant: the pre-trained backbone, the model after functional-track post-training, and the annotation and generative fine-tunes. Fig. 1 labels the four stages individually, because it draws the lineage.

#### 4.1.1. Model Architecture

NTv3 follows a U-Net [71] encoder-decoder architecture tailored for long sequences, similar to recent models such as Borzoi [5] and AlphaGenome [6]. The model consists of six main components: sequence embedder, sequence encoder, central transformer tower, sequence decoder, species conditioning and output heads. The pre-trained NTv3 backbone has approximately 703 million trainable parameters, rising to 711 million once the functional-track head is added. Of the post-trained parameters, 22% are in the encoder, 49% in the sequence transformer, 28% in the decoder and 1% in the functional-track head; the language-model head accounts for a further 0.002%. Below we describe the model architecture’s components along with relevant pseudocode.

##### Sequence Embedder

An embedding layer first transforms DNA sequences into sequences of tokens with continuous embeddings. Pre-training used an 11-token vocabulary: six special tokens followed by the five nucleotide symbols in the order A, T, C, G, N at indices 6 to 10. The post-trained tracks and annotation NTv3 models instead use a 16-token soft-masked vocabulary, which appends the lowercase symbols a, t, c, g, n at indices 11 to 15 so that repeat-masked reference sequence is represented explicitly; the uppercase indices are unchanged between the two vocabularies. The expansion changes exactly three tensors: the token embedding and the kernel and bias of the language-model output head. In each, the five lowercase rows were initialized as copies of their uppercase counterparts, so the expanded model is functionally identical to its predecessor at initialization. The language-model head received no gradient during functional-track post-training, which sets the masked-language-model loss weight to zero.

Token embeddings of dimension 16 are then processed by a stem consisting of a 1D convolution with kernel size 15 and stride 1, which projects them to the 1024 channels that open the encoder while preserving the sequence-length dimension, followed by a residual branch of a normalization, an approximate GELU [72] activation and a further kernel-5 weight-standardized convolution.

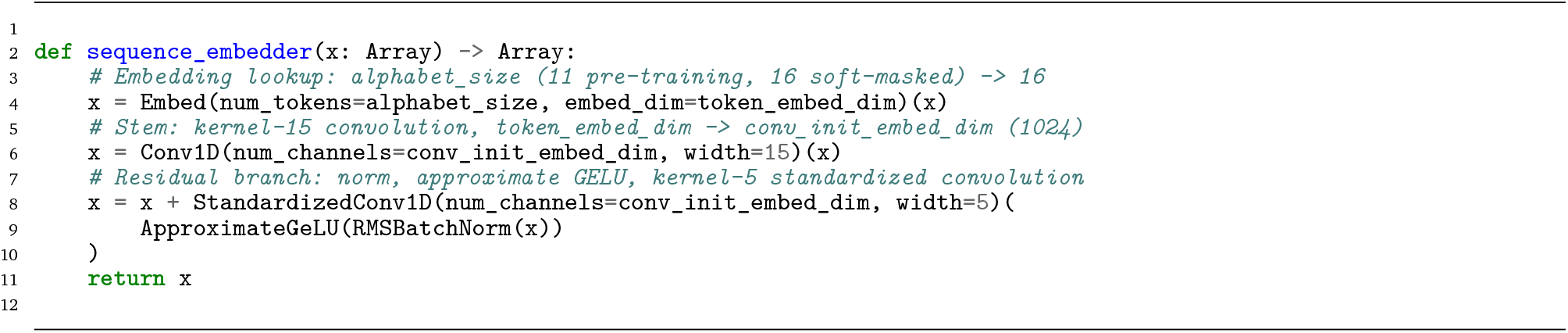

##### Sequence Encoder

The sequence encoder progressively down-samples the input DNA sequence from 1 bp resolution to 128 bp resolution embeddings. Down-sampling is applied in seven steps: a single max-pooling layer (window size 2, stride 2) immediately after the stem, followed by six residual convolutional blocks that each halve the sequence length, for a total down-sampling factor of 2^7^ = 128. Each convolutional block contains two convolutions of kernel size 5 in a pre-normalized arrangement, in which a root-mean-square batch normalization and a GELU activation are applied before each convolution rather than after it, composed as a residual pair [73]: the first convolution is added to the block input and the second is added to that sum. Where a block widens the representation, the skip path is zero-padded along the feature axis rather than projected, so no additional convolution is introduced on the residual branch. A max-pooling layer (window size 2, stride 2) then halves the sequence length.

Unlike architectures that hold the channel width fixed, the encoder widens monotonically: the number of feature channels is interpolated linearly from 1024 at the stem to 2048 at the transformer trunk across the six blocks, giving the schedule 1024, 1194, 1365, 1536, 1706, 1877, 2048. Intermediate channel counts are therefore not constrained to powers of two.

Intermediate representations from each down-sampling stage, corresponding to resolutions of 1 bp, 2 bp, 4 bp, …, 64 bp (before the pooling step), are stored for use as U-Net skip connections in the sequence decoder. The final encoder output has a resolution of 128 bp, sequence-axis length 1 Mb/128 bp = 8192, and 2048 feature channels.

All kernel-5 convolutions in the stem, encoder and decoder are weight-standardized with a learnable per-channel scale (the kernel-15 stem convolution and the 1 × 1 U-Net skip projections are not), and the convolutional activation is the sigmoid approximation *σ*(1.702*x*) · *x* to GELU rather than the tanh approximation used inside the transformer. The normalization used throughout the convolutional stages is a root-mean-square batch normalization, which rescales by a running estimate of the second moment without mean centering; because it carries running statistics, model outputs differ between training and evaluation modes.

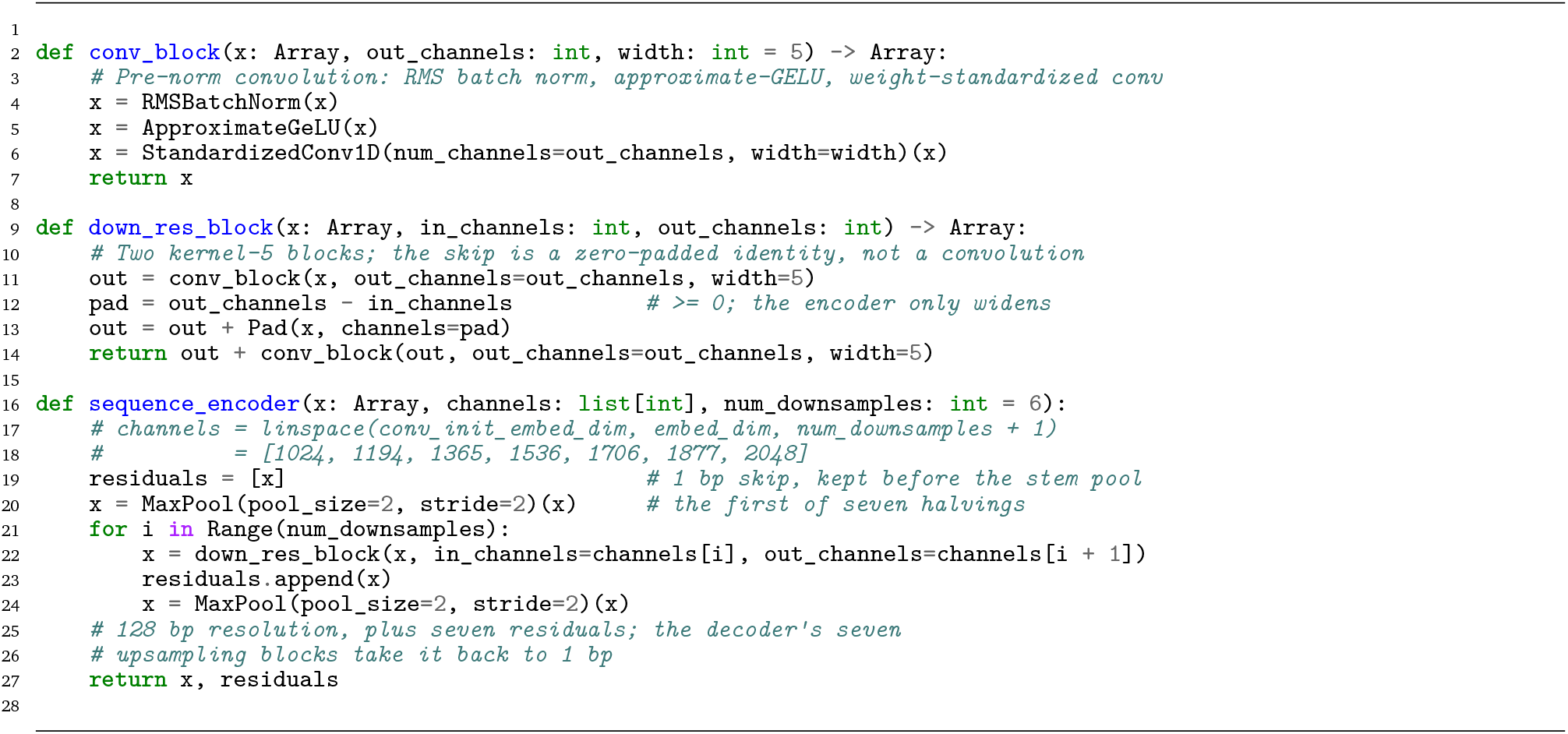

##### Transformer Tower

Following the encoder, a transformer tower [74] of 14 layers processes the 128 bp resolution sequence embeddings to model long-range interactions across the full 1 Mb input context. Attention is *global* over the down-sampled sequence: every trunk position attends to every other, which at 1 Mb input is an 8192 × 8192 score matrix. No windowed, blocked or sliding-window attention is used. Positions that fall entirely within padded input are masked out, and because one trunk position summarizes 128 input nucleotides, this validity mask is computed by grouping the input into blocks of 2^num_downsamples+1^ = 128 tokens and marking a trunk position valid if its block contains any non-padding token. Normalizations in the transformer are RMSNorms, applied before and after each attention and feedforward sub-layer and to the transformer output before it is consumed by the decoder, with a single exception: the per-head layer normalizations of the queries, keys and values. The output heads use layer normalization.

Each layer uses grouped-query attention [75] with 8 query heads and 2 key/value heads, giving a group size of 4, and a per-head dimension of 192 for queries, keys and values. A LayerNorm [76] is applied per head to each of the query, key and value projections; these keep the attention logits bounded, so no logit soft-capping is used. Each layer employs pre- and post-normalization [77] with root-mean-square norms around self-attention and a gated feedforward network [78], both wrapped in residual connections. The feedforward network is a GeGLU: the gate branch passes through a tanh-approximated GELU [72] and multiplies the value branch, with a hidden width of ⌊4*d*_model_/3⌋ = 2730 chosen so that the gated network matches the parameter count of a conventional 2× expansion.

Positional information is supplied by rotary positional embeddings (RoPE) [79] applied to the query and key projections, rotating each consecutive even/odd channel pair. Our frequencies follow a genomic rather than a language-model recipe: instead of the usual geometric spacing base^−2*i*/*d*^, inverse frequencies are set to 1/(*i* + *g*_*i*_), where *g*_*i*_ is geometrically spaced from 1 to 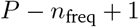 over the *n*_freq_ = *d*/2 frequency bands and *P* = 8192 is the largest trunk position, that is, a 1 Mb input at 128 bp resolution. This places the high-frequency bands on a near-integer spacing while the low-frequency tail is dominated by the geometric term, so that the longest period covers the full input context. No base frequency and no rescaling factor are involved.

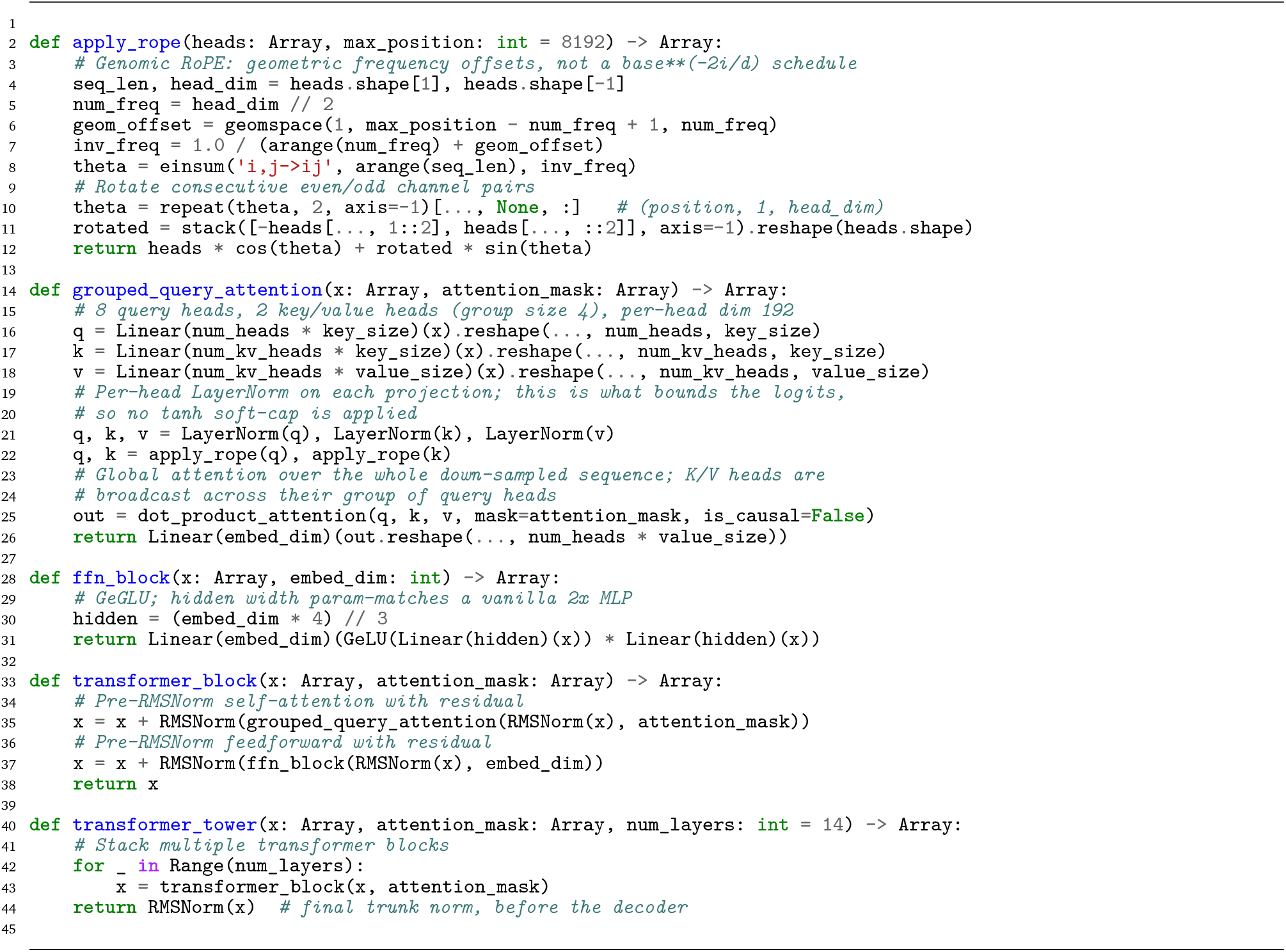

##### Sequence Decoder

The sequence decoder mirrors the encoder and operates in reverse, progressively upsampling the 128 bp resolution embeddings produced by the transformer tower back to 1 bp resolution while narrowing the representation along the encoder’s channel schedule in reverse. Each block applies a kernel-5 pre-normalized convolution with a residual connection, upsamples the sequence axis by a factor of two by *nearest-neighbor repetition* rather than a transposed convolution, scales the result by a learned scalar initialized to one, and then adds the matching encoder representation through a 1 × 1 projected U-Net skip [71]; a second kernel-5 convolution is added residually to close the block. These skip connections allow the decoder to recover the fine-grained sequence information lost during down-sampling.

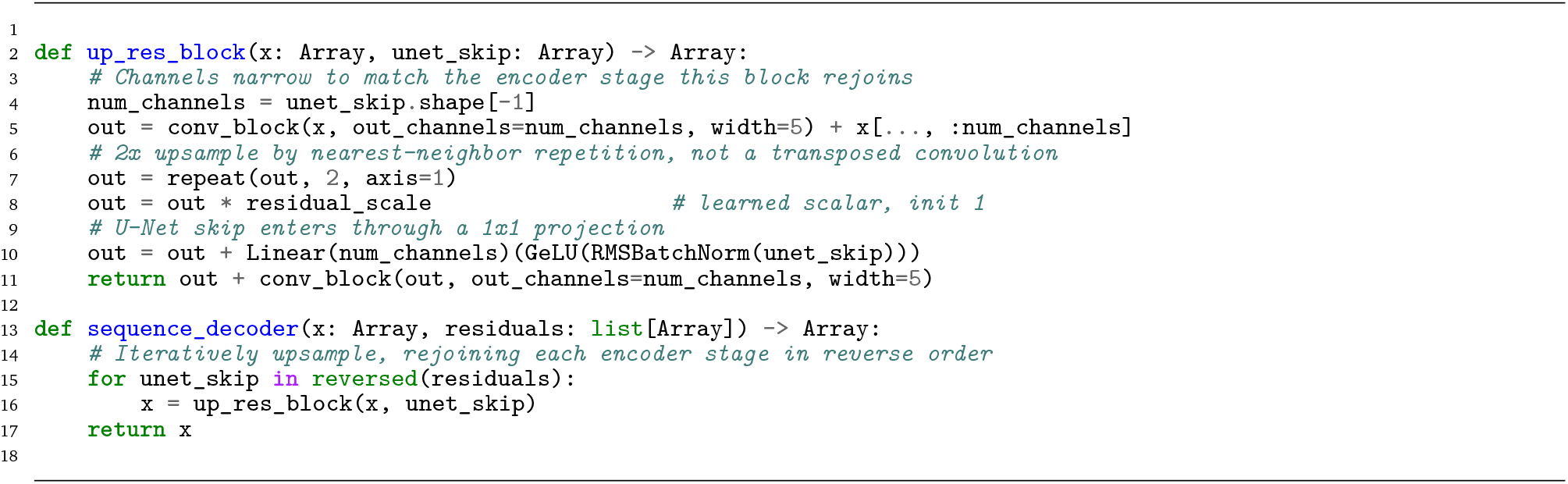

##### Output Heads

The NTv3 backbone produces a base-resolution representation that is mapped to different outputs by task-specific heads depending on the model variant.

###### Language-model head

A GELU nonlinearity followed by a linear projection, predicting a probability distribution over the DNA token vocabulary at each position. This head was optimized during masked-language pre-training and was subsequently reused for masked-diffusion sequence generation.

###### Functional-track head

NTv3 uses a linear head that predicts each of the functional tracks for the specified species. Species identity modulates the head twice, through adaptive layer normalization on its input, before a shared linear projection, and a per-species additive term on its logits, after which a softplus activation produces non-negative base-resolution track predictions. Species-specific padding and loss masks accommodate differences in the number of available tracks between genomes.

###### Gene-annotation head

NTv3 uses a classification head that predicts fifteen gene-structure classes on a single strand at each position, followed by the structured decoder described in Section 4.4. Unlike the functional-track head it is left unconditioned, and the opposite strand is recovered by a reverse-complement forward pass.

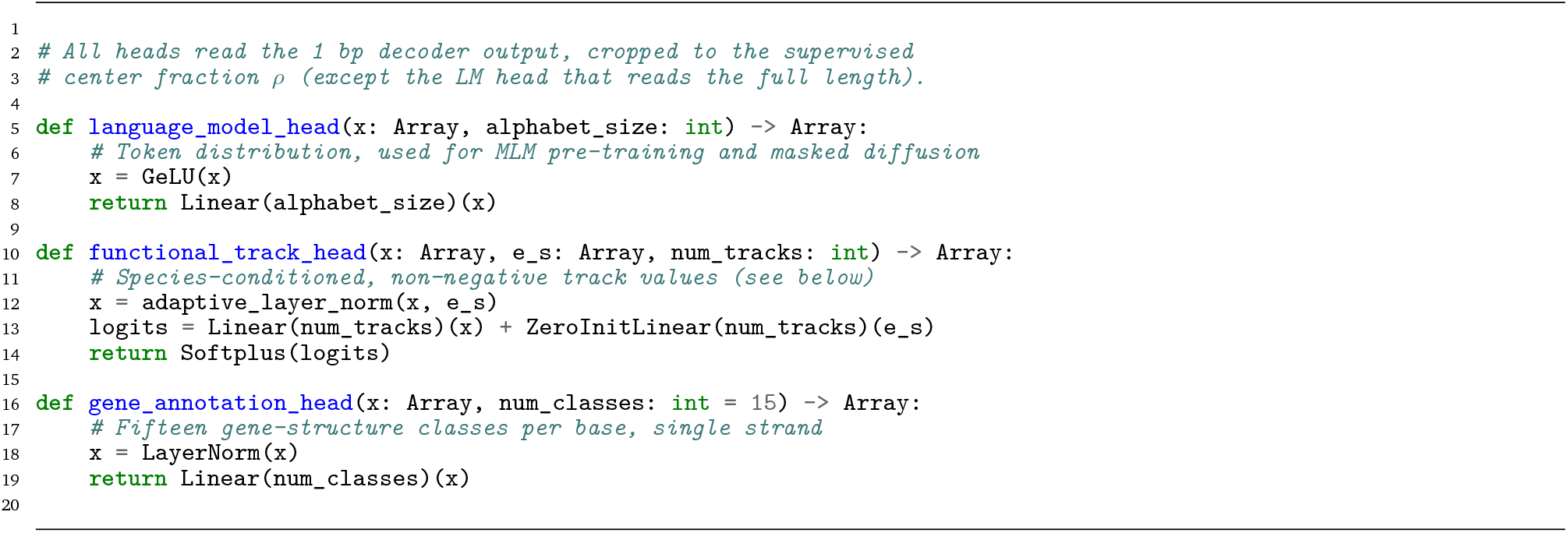

##### Species-conditioned Tracks Head

Species conditioning was used for multispecies functional-track prediction. Let *h*_*i*_ ∈ ℝ^*d*^ denote the base-resolution decoder output at position *i*, and let *e*_*s*_ denote a learned embedding for species *s*. Two learned projections map *e*_*s*_ to channel-wise scale and shift vectors, *γ*_*s*_, *β*_*s*_ ∈ ℝ^*d*^. The representation supplied to the functional-track head is

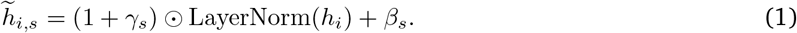

The scale and shift projections were initialized to zero, such that conditioning initially reduced to ordinary layer normalization. A single linear projection was then shared across species, followed by a second, species-specific additive term *b*_*s*_ applied to the logits and also initialized to zero, so that at initialization the head reduced exactly to an unconditioned linear head:

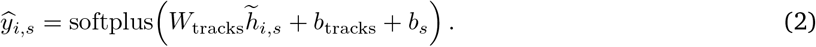

Species identity was supplied only to this output head; the convolutional encoder, transformer and decoder received DNA sequence alone. Thus, all species shared the same trunk and functional-track projection, while adaptive normalization allowed the head to adjust its predictions to the input genome and respective tracks.

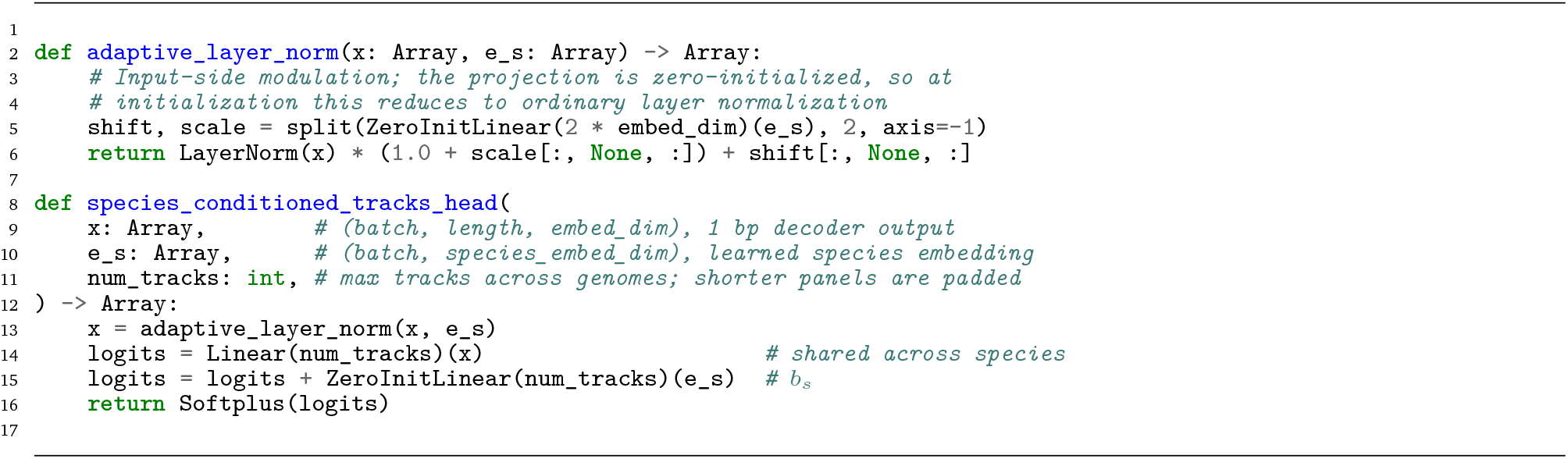

### 4.2 Pre-training on Genomes

NTv3 was pre-trained on the OpenGenome2 dataset through a self-supervised MLM objective. The pre-trained models have a single output head, the language-model head. We used the same approach as for the Nucleotide Transformer model [19], following the standard BERT masking strategy [35]: for each sequence, 15% of tokens were selected for potential modification; of this subset, 80% were replaced with a [MASK] token, 10% were substituted with a random token from the vocabulary, and the final 10% remained unchanged. For each sequence, the loss was computed as the sum of the cross-entropy losses between the predicted probabilities over tokens and the ground-truth tokens at the selected positions, divided by the number of non-padding positions in that sequence; the batch loss is the mean over sequences. Note that the denominator counts all non-padding input positions rather than only the selected ones, so the reported loss is smaller than a conventional per-masked-token cross-entropy by a factor of approximately the masking rate.

Pre-training was conducted in two main phases with different sequence lengths for performance optimization: a 8kb-training phase with sequences up to 8,192 token context focused primarily on functional elements, that accounted for ∼95% of all tokens seen during pre-training; and a sequence length extension phase during which we extend the sequence context progressively up to 1 Mb and add whole genomes to the data distribution, while continuing to interleave all previously seen sequence lengths. This ensured that the model retained strong performance on shorter contexts. Exact sequence length proportions per phase are detailed in Table 1. We refer to the models after pre-training but before post-training as NTv3 (pre).

**Table 1.** Sequence length distribution at each stage of training.

| Stage | Sequence length |  |  |  |  |  |  |  |  |  |  | Tokens (B) |
| --- | --- | --- | --- | --- | --- | --- | --- | --- | --- | --- | --- | --- |
| | $2^{10}$ | $2^{11}$ | $2^{12}$ | $2^{13}$ | $2^{14}$ | $2^{15}$ | $2^{16}$ | $2^{17}$ | $2^{18}$ | $2^{19}$ | $2^{20}$ | |
| 8 kb Pre-training | 0.079 | 0.130 | 0.132 | 0.660 | 0 | 0 | 0 | 0 | 0 | 0 | 0 | 5600 |
| Length Extension 1 | 0.052 | 0.077 | 0.079 | 0.394 | 0.398 | 0 | 0 | 0 | 0 | 0 | 0 | 50 |
| Length Extension 2 | 0.056 | 0.056 | 0.056 | 0.225 | 0.228 | 0.379 | 0 | 0 | 0 | 0 | 0 | 50 |
| Length Extension 3 | 0.053 | 0.053 | 0.053 | 0.129 | 0.130 | 0.218 | 0.363 | 0 | 0 | 0 | 0 | 50 |
| Length Extension 4 | 0.052 | 0.052 | 0.052 | 0.076 | 0.076 | 0.127 | 0.212 | 0.353 | 0 | 0 | 0 | 50 |
| Length Extension 5 | 0 | 0 | 0.050 | 0.050 | 0.050 | 0.084 | 0.140 | 0.233 | 0.394 | 0 | 0 | 50 |
| Length Extension 6 | 0 | 0 | 0.050 | 0.050 | 0.050 | 0.050 | 0.079 | 0.131 | 0.221 | 0.369 | 0 | 50 |
| Length Extension 7 | 0 | 0 | 0.050 | 0.050 | 0.050 | 0.050 | 0.050 | 0.074 | 0.124 | 0.207 | 0.345 | 50 |

**Table 2.** Token count by sequence length and OpenGenome2 sub-dataset. This dataset consisted of all sequences longer than 1 kb. Sequences longer than 8 kb are chunked to 8 kb. The total number of tokens is 1,808,911. All cells are in millions of tokens, so the table describes approximately 1.8 trillion tokens in total.

| Sequence Length | Eukaryotic Genic Windows | GTDB v220 | IMGPR | IMGVR | Metagenomes | mRNA | mRNA Splice | ncRNA | Organelles | Transcripts |
| --- | --- | --- | --- | --- | --- | --- | --- | --- | --- | --- |
| 1,024 - 2,048 bp | 2027 | 1516 | 0 | 124 | 132013 | 12747 | 9410 | 3057 | 0 | 603 |
| 2,048 - 4,096 bp | 4899 | 12000 | 27 | 520 | 184436 | 17940 | 38018 | 2397 | 0 | 2146 |
| 4,096 - 8,192 bp | 11943 | 26433 | 352 | 7911 | 169243 | 13256 | 29753 | 1682 | 0 | 3563 |
| ≥8,192 bp | 395278 | 314785 | 5370 | 29748 | 344121 | 7294 | 14277 | 4971 | 2784 | 2267 |

#### 4.2.1. OpenGenome2

We pre-trained NTv3 on a subset of the OpenGenome2 dataset introduced in Evo2 [25], which originally comprises 8.84 trillion non-redundant nucleotides spanning diverse taxa and genomic contexts. Specifically, the dataset includes 357 billion nucleotides from prokaryotic genomes, 6.98 trillion nucleotides from eukaryotic genomes, 854 billion nucleotides from metagenomic sequencing data, 2.82 billion nucleotides from organellar genomes, and 602 billion nucleotides from targeted subsets of eukaryotic sequences enriched for functional regions centered on protein-coding genes. The original dataset can be found here: https://huggingface.co/datasets/arcinstitute/opengenome2.

For pre-training NTv3, we applied two filtering steps to this dataset. First, we removed all sequences shorter than 1 kb, which primarily excluded promoter regions and accounted for a negligible number of nucleotides. Second, within the `ncbi_eukaryotic_genomes` subset, we retained only sequences longer than 32 kb; this filtering removed approximately 800 billion nucleotides. After these steps, the total number of unique nucleotides used for pre-training was approximately 8 trillion composed of 6.2 trillion from the `ncbi_eukaryotic_genomes` subset, and 1.8 trillion from all other subsets.

#### 4.2.2. Data Processing and Augmentation

To train the model, we utilized the original OpenGenome2 splits for train, test and validation and divided each fasta file into fixed-length genomic windows without overlap. In the pre-training of NTv3 we sought to avoid padding as padded tokens can introduce distributional artifacts, reduce the proportion of informative positions contributing to the loss, and create boundary effects that are incorporated into the receptive fields and can be particularly detrimental for U-Net architectures.

To eliminate the need for padding, we restricted training to sequence segments longer than the model’s input length at each stage. Since genomic sequences rarely have lengths that are exact multiples of the training window, a single fixed tiling would leave parts of each sequence uncovered. To ensure complete coverage, we generated two complementary tilings for every FASTA file. The first divides the sequence into non-overlapping windows starting from the first base and discards any leftover bases at the end. The second performs the same operation in reverse, starting from the final base and discarding any incomplete window at the beginning. By mixing segments from both tilings during training, the model sees every genomic region in at least one full-length window, entirely avoiding padding.

#### 4.2.3. Pre-training Data Composition and Sampling

During the initial pre-training phase, which used sequences up to 8,192 bp, we followed prior work [25] and trained on all subsets of the OpenGenome2 dataset, with the exception of whole eukaryotic genomes (`ncbi_eukaryotic_genomes`). These genomes, though comprising 6.98 trillion nucleotides, are dominated by long intergenic regions. Segmenting them into 8 kb windows would often yield sequences with minimal functional content. We therefore deferred their inclusion until the model was able to process longer contexts. This approach leaves a pre-training dataset of approximately 1.8 trillion nucleotides, enriched for gene-proximal functional regions.

To ensure meaningful input for the transformer, we further filtered sequences shorter than 1,024 bp, which correspond to fewer than eight tokens in the transformer. This filtering removes the Promoters subset of OpenGenome2, but its contribution to the total token count is negligible.

For 8kb-training, sequences were partitioned into length-specific data sources, where datasource_*i*_ contained all sequences *s* such that:

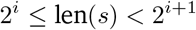

for *i* = 10, 11, 12 (1 kb, 2 kb, 4 kb). All sequences ≥ 8 kb were placed in a final bin. In total, this yielded four distinct data sources, collectively partitioning all eligible tokens in OpenGenome2 while excluding whole genomes and sequences shorter than 1,024 bp.

Training batches were sampled from these bins following the strategy in Section 4.2.2. Each sequence was segmented twice at its target length: once in a left-to-right pass and once in a right-to-left pass. The resulting paired segmentations were then combined and shuffled, yielding four data sources of fixed-length sequences. This procedure ensures full coverage of all pre-training tokens. Most tokens appear in both views, while only a small fraction near the sequence boundaries are included once.

In training we specified sampling weights for each sequence length (data source), and, in this way, maintain precise control over the fraction of tokens seen at each sequence length. For 8kb-training, these weights were set proportional to each bin’s share of the total token count. The final sampling weights for the 1 kb, 2 kb, 4 kb, and 8 kb bins are presented in Table 3.

**Table 3.** Token distribution in the pre-training phase by sequence length.

| Sequence length range | % of total tokens |
| --- | --- |
| [1024, 2048) | 7.9 |
| [2048, 4096) | 13.0 |
| [4096, 8192) | 13.2 |
| [8192, ∞) | 66.0 |

#### 4.2.4. Optimization and Data Loading

Both pre-training phases used the same training recipe, including all optimizer settings, learning-rate schedule, masking strategy, and batch configuration, only differing in the sequence length schedule. Key hyperparameters are detailed in Table 4. We trained all models using the AdamW optimizer [80] with *β*_1_ = 0.9 and *β*_2_ = 0.98. Similar to prior work [25, 81, 82], we found that reducing *β*_2_ from the canonical 0.999 was crucial for maintaining stability when training our largest models with the largest effective batch sizes.

**Table 4.**
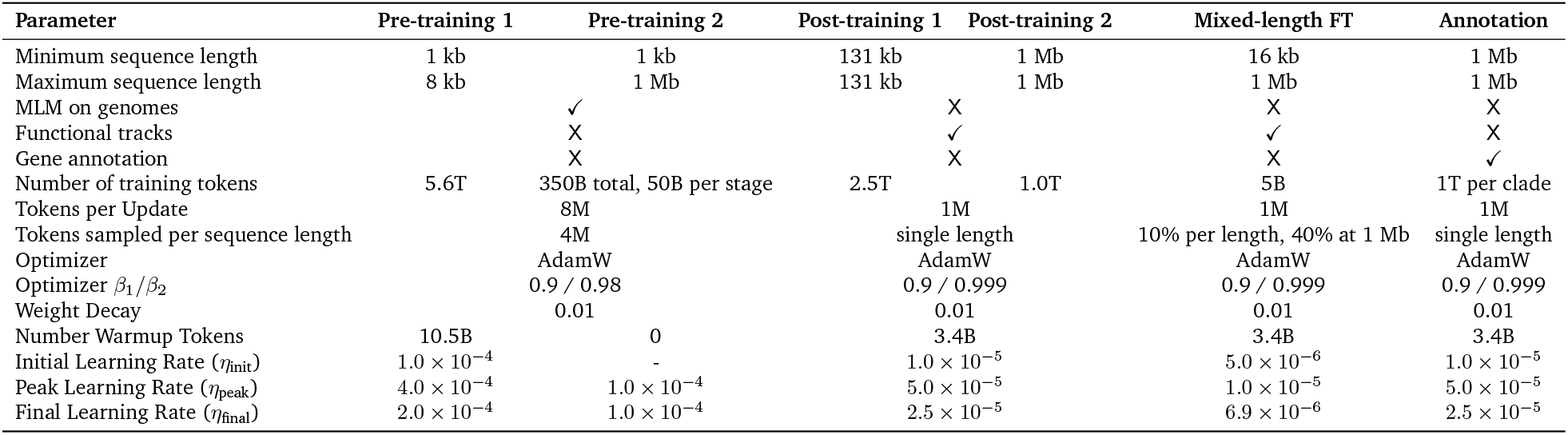
Training parameters for each training stage. Pre-training phases 1 and 2 correspond to 8kb-training and sequence length extension respectively. Post-training on functional genomic tracks is split into phase 1 at a fixed 131 kb context and phase 2 at a fixed 1 Mb context, both trained on bigWig tracks alone. The mixed-length column is a short fine-tuning of the post-training phase 2 checkpoint on inputs of 16 kb to 1 Mb (Section 4.3.9). The annotation column describes the clade-specific NTv3 fine-tunes, which are separate runs initialized from NTv3 and trained on gene annotations alone; the mammal, plant and insect runs share these settings, each with its own 1T-token budget. Each stage optimizes a single objective: masked language modeling during pre-training, functional-track prediction during post-training, and gene annotation during fine-tuning.

In both stages, gradients were accumulated to reach an effective batch size of 8 million tokens. We observed that increasing the batch size further improved training efficiency but led to degraded model performance. The learning rate followed a modified polynomial decay schedule with a linear warm-up. It started at *η*_init_ = 1.0 × 10^−4^, increased linearly to a peak of *η*_peak_ = 4.0 × 10^−4^ over the first 10.5 billion tokens, or approximately 1,312 steps with an effective batch size of 8 million tokens. After reaching a peak the learning rate decayed to a final value of *η*_final_ = 2.0 × 10^−4^. Formally, the schedule is defined at the step level as

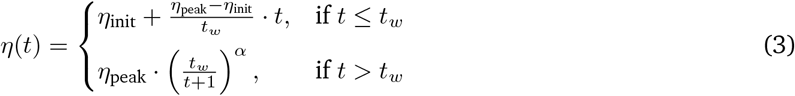

where *t* is the current step, *t*_*w*_ is the number of warm-up steps, and the decay exponent *α* = ln(1/*f*)/ ln(*T* /*t*_*w*_) is chosen such that the learning rate decays to *η*_final_ = *f* · *η*_peak_ after *T* total steps. In our experiments, we set *f* = 0.5.

To optimize data loading on our hardware, we structured the sampling process such that 4 million tokens of a single sequence length were yielded consecutively before moving to the next sequence length. Specifically, a sequence length was sampled according to the probabilities specified in Table 1, and mini-batches were yielded until a total of 4,194,304 tokens (2^22^) was collected, with the additional constraint that each mini-batch contained a number of tokens that evenly divided 4 million for all sequence lengths. Sequence lengths were sampled in this manner until the accumulated total reached 8 million tokens, at which point a gradient step was taken. This design allowed each optimization step to include multiple sequence lengths, thereby increasing gradient diversity. Using fewer than 4 million tokens per sequence length could have allowed for an even greater diversity of sequence lengths per accumulation step; however, in later stages with longer sequences, this would have constrained data parallelization, as it would have limited the number of tokens that could be processed per step to be below 4 million.

#### 4.2.5. 8kb Pre-training

The 8kb-training phase was run for 5.6 trillion (T) tokens on a filtered subset of OpenGenome2 that excludes whole eukaryotic genomes to focus on functional gene-rich regions and comprised approximately 1.8 trillion tokens in total. We train on a mixture of sequences of 1,024, 2,048, 4,096, and 8,192 bp to maintain performance across the different context lengths. For this initial pre-training phase, sequence lengths were sampled in proportion to the total number of tokens available at each length, with detailed distributions provided in Table 1. The duration of 8kb-training corresponded to approximately 3.1 epochs over the 8kb-training filtered dataset. A full description of the dataset composition and sampling approach is presented in Section 4.2.1. Tokenization is described in Section 4.1.1.

Training exhibited stable dynamics throughout, without the need for gradient clipping or restarts. Checkpoints were saved every 100 billion tokens, and the final 5.6T-token checkpoint was used to initialize the subsequent sequence-length extension phase. The loss continued to improve steadily through the end of this stage and training was stopped for time considerations.

#### 4.2.6. Sequence Length Extension

After completing 8kb-training, we implemented sequence length extension to adapt the model for processing of longer sequence contexts. Although our architecture supports long-context processing, we kept sequence length extension shorter than 8kb-training for compute efficiency, following prior work [25]. We conducted the sequence length extension in seven stages, each consisting of ∼50 billion (B) tokens. The first stage introduced sequences of 16 kb. At each subsequent stage, the context length was doubled, culminating in sequences of 1 Mb. Throughout sequence length extension, we continued to mix in all shorter sequence lengths, maintaining the ability to process shorter contexts, and preventing performance degradation on the corresponding distributions of short sequences. At the start of each stage, 40% of tokens were drawn from the newly introduced length, while the remaining probability was distributed among shorter sequences. Additionally, we enforced a minimum of 5% probability per sequence length to ensure the shortest lengths continued to be seen in training, and re-normalized the weights accordingly. For stages containing sequences ≥262kb we required sequence parallelism and as such excluded sequences with length ≤2kb for efficiency. Details of the sequence lengths and data distributions across 8kb-training and sequence length extension phases are provided in Table 1, and Supplementary Note B.2 describes how the per-stage sampling weights were constructed, including the basic schedule and the retirement of the shortest length bins. Training hyperparameters were identical to those used in 8kb-training and are detailed in Table 4.

### 4.3 NTv3: Post-training on Functional Genomic Tracks

The NTv3 tracks model was obtained by post-training the pre-trained backbone on base-resolution functional genomic tracks. The sequence trunk was initialized from the final pre-training checkpoint, and the species-conditioned functional-track prediction head was initialized randomly. Post-training used 16,262 tracks across fifteen animal and plant genomes (Section 4.3.1).

Post-training comprised two consecutive phases. In the first phase, the model received fixed-length inputs of 131,072 bp and processed 2.5 trillion nucleotide tokens. In the second phase, the input length was increased to 1,048,576 bp and training continued for a further 1 trillion tokens. Unlike the context-length curriculum used during pre-training, neither phase mixed different input lengths, since the goal was to focus supervision on the 1-Mb scale at which the model is evaluated. A mixed-length variant, obtained afterwards by a short fine-tuning, is described in Section 4.3.9. The first phase used data parallelism, whereas the 1-Mb phase used sequence parallelism across four devices. For both input lengths, functional-track supervision was restricted to the central 37.5% of the input (as done previously, [5]), corresponding to 49,152 and 393,216 predicted nucleotides, respectively. This cropping reduced boundary effects by providing flanking sequence context on both sides of every supervised position.

#### 4.3.1. Multispecies Genomic Tracks Dataset

Our post-training dataset is composed of 16,262 functional genomic tracks for fifteen different animal and plant species. The functional tracks include RNA-seq, CAGE, ATAC-seq, DNase-seq, and ChIP-seq data across fifteen species, totaling 16,252 tracks in the five assay classes reported here; the remaining ten are mouse GRO-seq tracks, which the model predicts but which we do not report as a separate assay class. The tracks are distributed across the following species: (human (*Homo sapiens*): 7,362; mouse (*Mus musculus*): 2,440; pig (*Sus scrofa*): 94; sheep (*Ovis aries*): 63; Horse (*Equus caballus*): 96; chicken (*Gallus gallus*): 55; zebrafish (*Danio rerio*): 23; fly (*Drosophila melanogaster*): 180; worm (*Caenorhabditis elegans*): 42; Arabidopsis (*Arabidopsis thaliana*): 1,899; rice (*Oryza sativa*): 1,392; maize (*Zea mays*): 921; wheat (*Triticum aestivum*): 776; soybean (*Glycine max*): 590; cotton (*Gossypium hirsutum*): 319). The data sources and pre-processing steps for each species and data type are detailed in Supplementary Note B.4.

The human and mouse functional tracks are based on the regulatory datasets used in Borzoi [5], including ENCODE v3, FANTOM5 CAGE, and GEO RNA-seq, ATAC-seq, DNase-seq, and ChIP-seq data. The *Drosophila* data are composed of the embryogenesis single-cell atlas of Calderon et al. [70]. Chicken, horse, pig, and sheep data were obtained from previously published studies from the Functional Annotation of Animal Genomes (FAANG) Consortium [83]. The zebrafish, worm and six plant species data were obtained from large public collections in NCBI repositories (e.g., SRA/GEO; https://www.ncbi.nlm.nih.gov/), using stringent quality filters detailed in Supplementary Note B.4 (Table 5 and Supplementary Table C.1).

**Table 5.** Functional tracks training data. The full dataset is composed of 16,262 tracks over fifteen genomes, ordered by species tree with animals above plants.

| Species | Chromatin accessibility | Histone marks and TFs | Gene expression |
| --- | --- | --- | --- |
| Human | 805 | 3,792 | 2,765 |
| Mouse | 287 | 1,027 | 1,126 |
| Pig | 39 | 36 | 19 |
| Sheep | 2 | 5 | 56 |
| Horse | 29 | 36 | 31 |
| Chicken | 22 | 15 | 18 |
| Zebrafish | 5 | - | 18 |
| Fruit fly | 90 | - | 90 |
| Worm | 4 | - | 38 |
| Rice | 13 | 23 | 1,356 |
| Wheat | 14 | - | 762 |
| Maize | 31 | 77 | 813 |
| Thale cress | 15 | 245 | 1,639 |
| Cotton | 8 | 25 | 286 |
| Soybean | 1 | 13 | 576 |

#### 4.3.2. Species Genome Data Splits

NTv3 was post-trained on the following fifteen species and reference assemblies: human (*hg38*), mouse (*mm10*), pig (*Sscrofa11*), sheep (*Oar_v3*.*1*), horse (*EquCab3*.*0*), chicken (*GRCg6a*), zebrafish (*GRCz11*), fly (*dm6*), worm (*WBcel235*), Arabidopsis (*TAIR10*), rice (*IRGSP-1*.*0*), maize (*Zm-B73-REFERENCE-NAM-5*.*0*), wheat (*IWGSC*), soybean (*Glycine_max_v2*.*1*) and cotton (*Gossypium_hirsutum_v2*.*1*). For each genome we created train, validation and test splits of DNA sequence controlled for sequence similarity as described below.

##### Human and Mouse

To remain comparable with existing models, the folds used by Borzoi [5] are used to define the training, validation and test sets. Specifically, fold3 is used for test, fold4 for validation and the rest are used for training. In contrast with previous approaches that directly use these sequences, we merge the sequences for each set into continuous regions. These regions are then tiled with the same stride used for every other genome, 0.1% of the sequence length (see *Genomic Sequence Selection* below); human and mouse differ from the other species in the source of the intervals being tiled, not in the step between them. This approach enables high sequence diversity, ensuring that the model seldom sees repeat samples.

##### Other Species

To control for data leakage within a species, we used a homology-based clustering approach. Contiguous spans on the genome were divided into 1 Mb gene-aware windows; that is, each window was expanded to include the last gene that starts within its given span, ensuring that no gene was split across windows. Chromosomes or scaffolds shorter than 1 Mb were treated as their own windows. Ensembl BioMart was used to extract homologous gene pairs. The average base pair size of the homologous genes across windows was then used to weight the edges of a symmetric window-to-window connectivity matrix. The Markov Clustering (MCL) algorithm [84] was used to create clusters among the connected windows. Inflation rates ranging from 1.5 to 5.0 were evaluated in increments of 0.5. The optimal rate was selected based on the highest modularity score [85]. The clusters were then stratified into train, validation, and test splits balancing gene-containing and gene-free windows and according to a predefined target validation and test size. For species where the algorithm could not meet the predefined target validation and test set sizes, edges representing a shared homology of less than 10% of the window size were pruned. These species included *A. thaliana, C. elegans* and *O. sativa*.

#### 4.3.3. Dataset Weighting

During post-training, genomes are sampled according to fixed weights rather than in proportion to their length, so that the frequency with which the model sees a species is independent of its genome size. This matters for a corpus that spans two orders of magnitude in assembly length. Human receives the largest weight because it carries by far the most tracks, mouse the second largest, and the remaining thirteen genomes share the rest equally (Table 6). The weights were obtained by taking the functional-track weights of an earlier joint annotation-and-tracks recipe and renormalizing them over the fifteen track-bearing genomes, which preserves the human-to-mouse ratio of that recipe.

**Table 6.** Genome sampling weights used during NTv3 post-training. All fifteen genomes contribute functional tracks; the weights are validated to sum to one and are not renormalized at run time.

| Group | Number of genomes | Individual weight | Total weight |
| --- | --- | --- | --- |
| Human (*hg38*) | 1 | 0.3125 | 0.3125 |
| Mouse (*mm10*) | 1 | 0.078125 | 0.078125 |
| Remaining track-bearing genomes | 13 | 0.046875 | 0.609375 |
| **Total** | **15** | **–** | **1.0** |

#### 4.3.4. Genomic Sequence Selection

Training windows are drawn according to the dataset weights by slicing the train set of the genome with a stride of 0.1% of the sequence length. For example, when using a sequence length of 131 kb, the train set would be sliced with a stride of 131 bp. This strategy ensures that each nucleotide is seen in many different contexts, which is expected to improve generalization. For validation and evaluation the stride is instead set equal to the supervised central fraction of the input, that is 0.375*L*, giving 393,216 bp at a 1 Mb input. The supervised regions of consecutive windows therefore tile the genome exactly, so each evaluated nucleotide is scored once and once only, which is what makes the concatenation used for the correlation metrics well defined.

#### 4.3.5. Functional Track Scaling

Due to the large range and skewed distribution of functional track data, the values are scaled before calculating the loss. We follow AlphaGenome’s scaling approach [6], applied in three steps in this order: each value is divided by the non-zero mean of its track; values from assays whose name contains RNA-seq then receive a power transformation *x*^0.75^; and finally a square-root soft clipping, 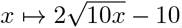, is applied to values above 10. Because the clip is applied last, it engages for RNA-seq tracks at a raw value of approximately 21.5 times the track mean rather than at 10 times. The targets are scaled before being used in the loss; when making predictions, the head output is unscaled to raw values. The per-track means and the RNA-seq mask were estimated on the training split only and held fixed for validation, testing and inference, so no normalization statistic is derived from evaluated data.

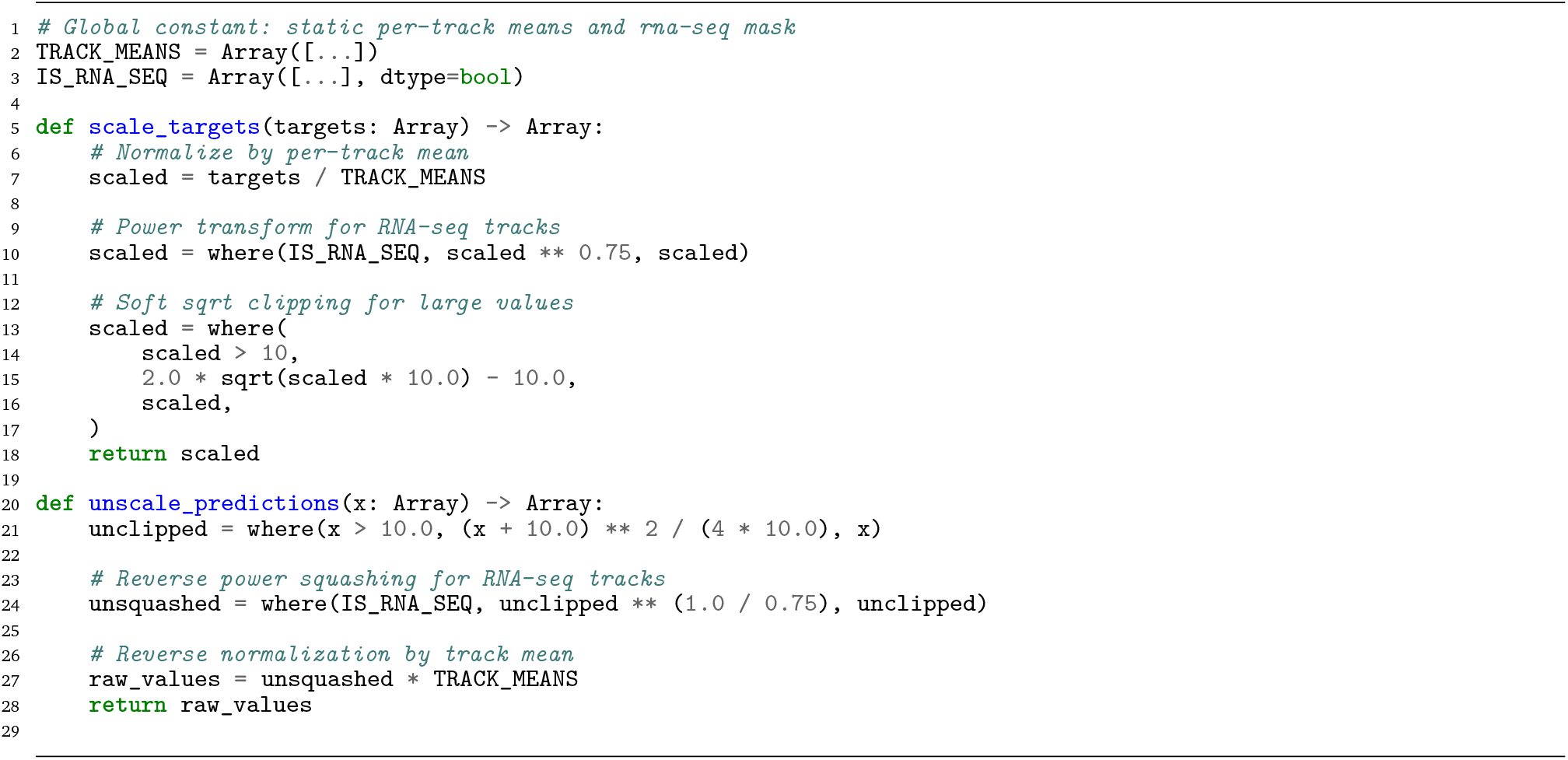

#### 4.3.6. Post-training Objective and Loss Function

Functional tracks were trained using a compound Poisson–multinomial objective, following previous sequence-to-profile models [4–6]. The Poisson component supervised the total signal within the output interval, whereas the multinomial component supervised its spatial distribution. The loss was calculated over the central 37.5% of each input sequence, providing flanking sequence context on both sides of every supervised position.

For batch element *b*, track *t* and the set of supervised positions *C*_*b,t*_ marked by the loss mask, we define the total predicted and observed signal as

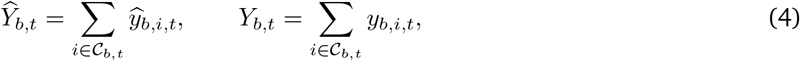

where 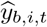 and *y*_*b,i,t*_ denote the predicted and observed signals respectively. Both components are stabilized by a small constant *ε* = 10^−7^, but not in the same way, and the equations below state the objective as implemented. In the Poisson scale term *ε* appears only inside the logarithm; the summed predictions and targets themselves carry none. In the multinomial shape term it is added to the predictions, to the targets, and once more in the normalizer, so the multinomial weights sum to slightly less than one rather than exactly one.

Omitting terms independent of the prediction, the Poisson scale loss for one batch–track pair was normalized by the number of supervised positions in that pair,

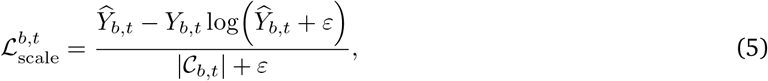

and averaged over the set *V* of batch–track pairs with at least one supervised position,

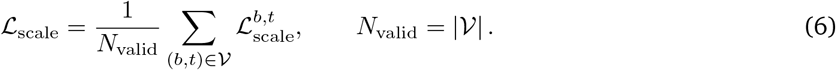

For the multinomial shape component, predictions were normalized across the supervised positions to obtain

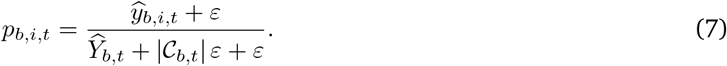

The shape loss was accumulated over every supervised position and normalized by their total number *M* =∑_*b,t*_ |*C*_*b,t*_|,

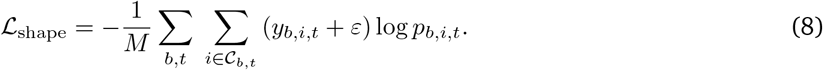

The two terms use different denominators: the scale loss averages one value per batch–track pair, whereas the shape loss averages over individual supervised positions. The Poisson scale loss was down-weighted fivefold relative to the multinomial shape loss, giving the complete functional-track objective

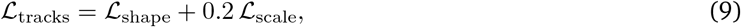

with padded track channels excluded by the loss mask.

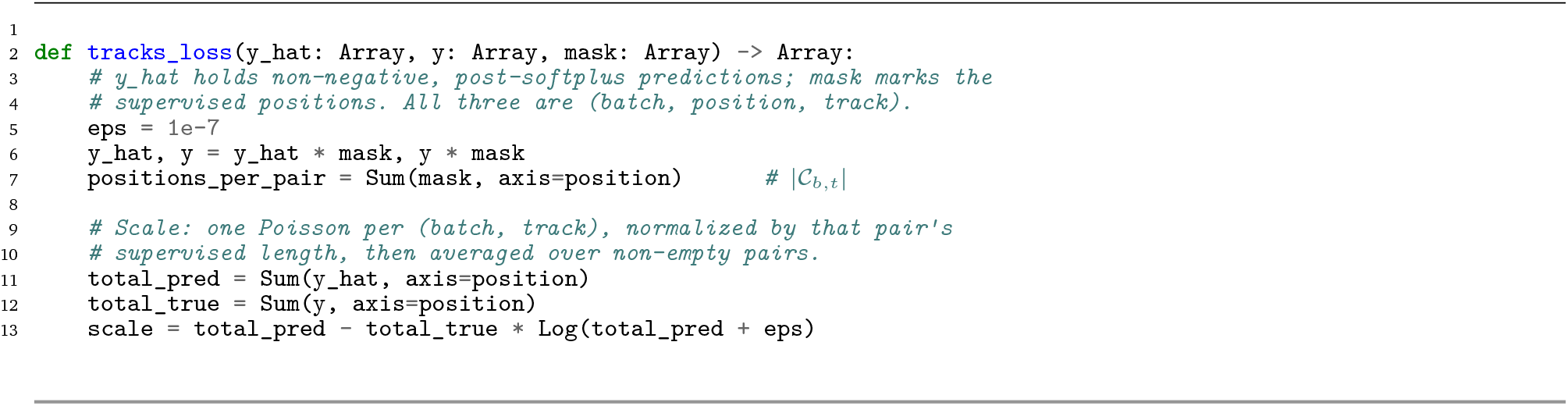
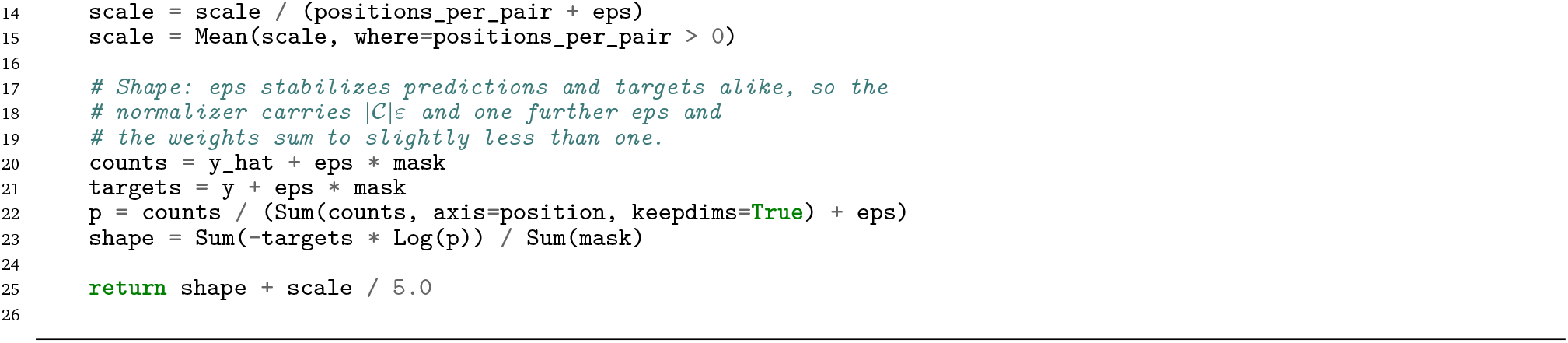

#### 4.3.7. Training Parameters

We describe the training parameters used during post-training, listed in Table 4. For post-training we use the AdamW optimizer with a weight decay of 0.01. The learning rate is increased linearly for the number of warm-up tokens and then decreased using the same polynomial decay as in pre-training over the number of training steps to the final value.

For the post-training, the learning rate schedule, including the warm-up, is restarted at the start of the second phase, and we used a lower learning rate compared to pre-training to retain the knowledge of pre-training in the model.

We found that using a smaller batch size of 1M tokens improved performance on track prediction, relative to larger batch sizes like 2M or 8M, similar to previous work [5].

#### 4.3.8. Evaluation Procedure

Post-training checkpoints were selected using the validation partition and evaluated on the held-out test partition. Predictions were first mapped from transformed signal space back to the original track scale. For every track *t*, predictions and targets from the supervised central regions of all test windows were concatenated before calculating Pearson correlation. Consecutive test windows were offset by the length of their supervised region, so that each position is scored once; no reverse-complement averaging and no mappability mask were applied.

We calculated correlations in raw signal space,

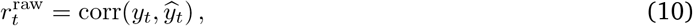

and after a log(1 + *x*) transformation,

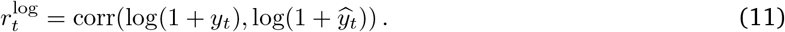

Species–assay scores were obtained as the unweighted mean across tracks belonging to the corresponding species and assay group. Post-training analyses report 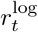 throughout, because at base-pair resolution raw coverage is dominated by a few extreme values (e.g. exon and TSS peaks), which log(1 + *x*) compresses so that the whole profile contributes. The one exception is human ATAC-seq in Fig. 2d,e, reported as 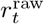, following AlphaGenome’s convention for accessibility assays [6]. Raw-space correlations are otherwise used only where required by an external benchmark protocol.

#### 4.3.9. Mixed-length Fine-tuning

Both post-training phases used fixed-length inputs (Section 4.3), so the single-length NTv3 model is intended for 1-Mb inputs and is less accurate on shorter ones (Supplementary Fig. A.4a). To provide a model that can be used at any input length from 16 kb to 1 Mb, we fine-tuned the final 1-Mb checkpoint on a mixture of input lengths: each 1,048,576-token update drew 10% of its tokens from each of 16, 32, 65, 131, 262 and 524 kb inputs and 40% from 1-Mb inputs. All other settings followed the second post-training phase (Table 4), except the learning rate, which was warmed up from 5 × 10^−6^ to 1 × 10^−5^ over 3.4 billion tokens and then decayed over a 30-billion-token schedule. Training used sequence parallelism across eight H100 GPUs.

##### Input-length benchmark

Each model was evaluated at seven input lengths, from 16 kb to 1 Mb, on identical scored regions. A region is the 393,216-bp supervised center of a 1-Mb input; at shorter lengths it was tiled by consecutive inputs whose supervised centers (37.5% of each input) abut, so that every length is scored on the same bases and only the context around them changes. Regions were drawn from the validation split to select the checkpoint, and from the test split to report performance and compare with the single-length model (Section 4.3.2). Accuracy is the Pearson correlation between predicted and observed log1p-transformed tracks at 1-bp resolution, pooled over regions and averaged over the tracks of each species, then over species.

##### Checkpoint selection and performance

We selected the checkpoint with the highest validation accuracy averaged over the six input lengths shorter than 1 Mb (16 to 524 kb): the checkpoint after 5 billion tokens, which is about 0.1% of the post-training token budget and cost about 10 GPU-hours. On validation regions, it was the most accurate at 16 and 32 kb (0.654 and 0.737) and within 0.003 of the best checkpoint at every longer input, including 1 Mb, where it remained above the single-length model (0.784 against 0.775). On test regions of the 15 post-training species, its accuracy rose with input length, from 0.656 at 16 kb to 0.737 at 32 kb, 0.764 at 65 kb and 0.777 at 131 kb, and reached 0.780 from 262 kb to 1 Mb, against 0.775 for the single-length model at 1 Mb, where it was higher in 14 of the 15 species (Supplementary Fig. A.4a).

##### Downstream benchmarks

To check that the fine-tuning preserves zero-shot downstream performance, we ran the mixed-length model through the eQTL and enhancer–gene benchmarks with the same protocols and 1-Mb inputs as the single-length model (Section 4.8.2). It matched the single-length model on eQTL classification (variant-weighted mean auROC 0.700 against 0.701 over 21,241 variants in 48 tissues) and on enhancer–gene linking (auPRC 0.575 against 0.582 over 10,356 pairs; Supplementary Fig. A.4b,c).

##### Interpretability agreement

We compared the single-length and mixed-length models on 24 randomly sampled genes in each of the 15 post-training species (Supplementary Fig. A.4d). Sampling was restricted to test-split genes with above-median measured RNA-seq signal near the TSS, with at least 10 kb between selected TSSs. Both models were evaluated on the same 1,048,576-bp TSS-centered inputs (at most 1% N bases; 2% in sheep), using the corresponding species token. For each species, track agreement is the median of the per-track Pearson correlations between log1p-transformed predictions over the central 393,216 bp. Attribution agreement is the cosine similarity between input-times-gradient maps over the central 49,152 bp. These maps were computed with respect to the log of the summed RNA-seq predictions over the first 2 kb downstream of the TSS and mean-centered across the four bases. Attention agreement is one minus the Jensen–Shannon divergence (base 2) between the models’ head-averaged last-layer attention profiles from the TSS bin to the remaining bins in the same 49,152-bp region. Across species, median track, attribution and attention agreement ranged from 0.987 to 0.997, 0.922 to 0.967 and 0.994 to 0.998, respectively. For the *HBE1* example (Supplementary Fig. A.4d, right), both models used the input window and K562 CAGE attribution target described in Methods 4.8.2.

### 4.4 NTv3: Fine-Tuning for *ab initio* Gene Prediction

#### 4.4.1. Model and Training

NTv3 uses the NTv3 U-Net backbone post-trained on multispecies functional tracks with a task-specific genome-annotation head in place of the functional-tracks head. We trained specific models for each clade: mammals, plants and insects. Each model was trained at a sequence length of 1,048,576 bp, for a budget of 1T tokens per clade.

##### Annotation head

The annotation head predicts fifteen classes at base-pair resolution, the label set of Tiberius [28]: *intergenic region*; *intron, CDS, splice acceptor*, and *splice donor*, each in three coding phases; *start codon* and *stop codon*. The phase index denotes the codon position of the labeled base. For introns, which carry no codon position of their own, it instead denotes the phase inherited from the upstream splice donor, so that coding frame is propagated across an intron rather than re-inferred after it. Jointly predicting gene states and coding phase distinguishes this task from nucleotide-level segmentation approaches such as SegmentNT, which label functional elements without resolving frame. The model uses a single-strand prediction head.

Reverse-complement augmentation is applied during training; at inference the minus strand is obtained from a second forward pass on the reverse-complemented sequence, mapped back to genomic coordinates and decoded by its own Viterbi pass with the emission constraints rebuilt on the reverse complement. The two strands are decoded independently and their gene sets concatenated without a merge step, so overlapping opposite-strand genes are both emitted. Annotating both strands therefore costs two forward passes and two decodes per window. Unlike the functional-track head, the annotation head receives no species conditioning: the adaptive normalization and the per-species additive term are disabled, so one set of weights is shared across every species in a clade and the prediction depends on the input sequence alone. Species identity therefore plays no role at inference, and an unseen genome is annotated by the same weights as a training genome.

##### Training objective

Training minimizes a masked, class-weighted focal loss over the fifteen classes at base-pair resolution, with focusing parameter *γ* = 2. Class weights are inverse class frequencies computed from the training annotations of each assembly, averaged across assemblies and normalized to mean one; each position is weighted by the larger of its target-class and predicted-class weight, so that false positives on rare classes are penalized at the rare-class weight. Writing *m*_*i*_ for the annotation mask, *w* for the mean-one inverse-frequency weights, *p*_*i,c*_ for the softmax probability of class *c* at position *i* and *ŷ*_*i*_ = arg max_*c*_ *p*_*i,c*_, the loss is

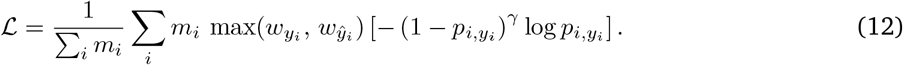

The arg-max carries no gradient, so the weight acts as a piecewise-constant multiplier and the objective is piecewise-smooth in the parameters, with a finite jump wherever the predicted class changes. Positions without an annotation target are masked from the loss. Annotation is the only training objective, with both the masked-language-model and functional-track losses disabled. Training examples expose the complete 1,048,576-bp input window to annotation supervision, without cropping the supervised region: writing *ρ* for the fraction of each window retained for supervision or prediction, annotation training uses *ρ* = 1.

##### Soft-masked inputs

Genome inputs are soft-masked and use the sixteen-token vocabulary described in Section 4.1.1, which represents lowercase nucleotides explicitly. Target construction and interval sampling use the same contig set, comprising all contigs of each training assembly.

##### Optimization and checkpoint selection

The annotation models were trained with AdamW using the modified square-decay schedule also used for functional-track post-training, with the learning rate increasing from 1 × 10^−5^ to a peak of 5 × 10^−5^. Warm-up, peak and final learning rates, tokens per update, the decay parameter and the token budget are given in the annotation column of Table 4. Training was performed in FP32 for numerical stability. Each update processes one 1,048,576-bp sequence, split along the sequence axis across all GPUs of a single node by sequence parallelism, with no data parallelism (Supplementary Note B.3). Activations are rematerialized in the backward pass so that the 1-Mb window fits in device memory. On four NVIDIA H100 80-GB GPUs, the mammal model reached its 1T-token budget in about eight days; the plant and insect models used the same configuration and comparable compute. Checkpoints were selected based on the highest mean gene-level F1 on the held-out validation species, decoded with the post-processing decoder described below. The reported checkpoints are step 925,089 for mammals, 877,404 for plants and 762,960 for insects. The full validation curves are in Supplementary Fig. A.5.

##### Post-processing decoder

Converting per-base posteriors into transcripts is where *ab initio* callers differ most. Helixer trains its network on base-wise labels alone and assembles gene models afterwards with a separate hidden Markov model, HelixerPost, which is decoded with the Viterbi algorithm under hand-set penalties [27]. Tiberius adopts this decoding step but places the hidden Markov model inside the network as a differentiable layer: the loss is computed on the layer’s posterior state probabilities and back-propagated through it into the network, while the layer’s 24 parameters stay fixed at hand-set values derived from expected region lengths [28], [29]. NTv3 trains the network without the hidden Markov model, as Helixer does, and applies it only at inference. The decoder is the Tiberius hidden Markov model, reimplemented in JAX. Its fifteen-state grammar (also the class set of our annotation head), its transition structure and length priors, its splice-site and start- and stop-codon emission constraints, and its uniform initial distribution are all taken from Tiberius [28] and applied unchanged to all three clades. The length priors are per-base probabilities of leaving the exon, intron and intergenic states of 1/200, 1/4500 and 1/10000. The intron and intergenic states carry self-loops, so these rates give geometric mean lengths of 4,500 and 10,000 bp. We depart from Tiberius in three places. First, the emissions are the head’s log-posteriors, without the smoothing Tiberius applies to its class probabilities (*α* = 0.01). Second, Tiberius splits the exit mass of its middle coding state between the stop-codon and splice-donor exits, *p*_*e*_/2 each. `CDS_p1` instead gives each exit the full 1/200, so the per-base exit hazard alternates 1/200, 2/200, 1/200 with codon position and the expected coding run is 150 bp rather than 200 bp. Third, we add a second, parallel set of splice states for the rare minor-spliceosome (U12) introns, which begin with AT and end with AC instead of the canonical GT and AG. A coding exon enters this path with a prior probability of 1.05 × 10^−3^, the share of such introns in mammals; otherwise it behaves exactly like the canonical intron path and adds no output class to the annotation head. The full transition table is in Supplementary Note B.6. Although the decoder is conceptually a post-processing step, it is compiled together with the forward pass and runs on the same accelerator, so its cost is included wherever inference time is reported below.

Two properties of the grammar, both inherited from Tiberius [28], matter for the results and are therefore stated here. The state diagram, the full transition matrix, the emission constraints, and the Viterbi recursion are given in Supplementary Note B.6. First, the coding states are phase-tagged and transitions between intron phases are forbidden, so coding frame can only change by passing through an exon and is carried across introns rather than re-inferred after them. Second, splice-donor and splice-acceptor states are constrained to the corresponding sequence motifs with Tiberius’s weights: GT at log(0.99) and the rare GC at log(0.01) for donors and AG for acceptors, plus AT and AC on the U12 path. Every other dinucleotide is vetoed, so that no predicted intron boundary is inconsistent with the underlying sequence.

Decoding is performed independently for each inference window, and the resulting state paths are concatenated without re-decoding across window boundaries. Tiberius, by contrast, re-decodes two adjacent tiles together wherever a parse ends inside a gene [28]. Inference uses the same 1,048,576-bp sequence length as training, of which the central 75% (*ρ* = 0.75, or 786,432 bp) is retained as the prediction region. Retaining the central region gave higher accuracy than predicting over the full window while keeping whole-genome decoding efficient. Consecutive windows therefore overlap by 262,144 bp, their prediction regions abutting at a 786-kb stride, so each base is predicted once and carries at least 131 kb of flanking context. The decoder drops predicted genes with less than 201 bp of CDS and those whose spliced CDS contains an in-frame stop codon. At scoring time, the evaluator applies the same 201-bp minimum to every caller (Section 4.4.3).

##### Training-initialization ablation

In addition, we performed a training-initialization ablation at 131 kb on the mammal panel. This experiment is separate from the 1-Mb headline annotation models, so its increments are not additive onto them. The randomly-initialized model used a higher learning rate, increasing from 1 × 10^−4^ to a peak of 4 × 10^−4^, followed by a decrease to 2 × 10^−4^, while the genome-pre-trained and NTv3-initialized models used a fine-tuning schedule from 1 × 10^−5^ to a peak of 5 ×10^−5^. The retained fraction was matched across the three arms at inference.

##### Annotation completeness

Each genome is annotated from its complete whole-genome FASTA, and every record in the file is annotated; no contig is removed beforehand. The prediction regions are laid out to cover each contig exactly, from its first base to its last. Because only the central fraction of a window is predicted, covering the two ends requires windows that extend beyond the contig: at the start, the first window begins ahead of base 1 and the missing sequence is supplied as N padding, and at the end, the last window is shifted back so that its prediction region terminates on the contig’s final base rather than being padded past it. The shifted final region is the only place where two prediction regions overlap, and the overlapping bases are taken from the earlier of the two. A contig shorter than a single prediction region is annotated as one padded window rather than skipped. NTv3 therefore annotates every base of the assembly, and scoring covers the whole assembly (Section 4.4.3).

#### 4.2.2. Training and Evaluation Data

##### Sources

Assemblies and their reference GFF3 annotations were obtained from the public archive that distributes each. For **mammals**, NCBI RefSeq supplied 30 of the 32 assemblies, with *Homo sapiens* (hg38) and *Mus musculus* (mm10) taken from Ensembl. For **plants**, JGI Phytozome supplied 36 of the 60 and NCBI RefSeq 22, with Araport11 for *Arabidopsis thaliana* and Ensembl for *Triticum aestivum* (IWGSC). For **insects**, the assemblies are those of the Tiberius Insecta panel throughout, but the annotations differ by split: Tiberius’s own BRAKER annotations for the 93 training and six validation assemblies, and the NCBI RefSeq annotation for each of the ten test assemblies. The assembly accession and the annotation provenance of every species are listed in Supplementary Table C.2.

##### Labels

For every assembly, the reference GFF3 was converted into the fifteen-class base-resolution representation of Section 4.4: each base carries one of *intergenic region*; *intron, CDS, splice acceptor* or *splice donor* in one of three coding phases; *start codon* or *stop codon*. Positions outside that label set are masked from the loss.

##### Genus-exclusion criterion

In every clade, test species are separated at the genus level from both the training and the validation set: no test species has a congener in either. Validation species are held out at the species level only, and two share a genus with a training species: *Mus pahari* with *Mus musculus*, and *Anopheles coluzzii* with three *Anopheles* species in the insect training panel. The criterion is what determines the test panels below. The one Tiberius test species that does not satisfy this criterion is *Drosophila melanogaster*, which has six congeners in the insect training panel; it is also the sole insect contributing functional tracks to post-training, and is excluded from the insect evaluation on that ground.

##### Splits

Each clade panel of species is derived from the corresponding Tiberius panel. Every species we score is one Tiberius holds out, so the benchmark is its own rather than one of our choosing, and both systems are run and scored here on the same assemblies, against the same reference annotations, under the same matching rules. The three clade panels of species are given in Supplementary Table C.2. The three panels comprise 17 held-out species in total: two mammals, five plants and ten insects. Three of the species Tiberius holds out cannot be scored here at all, because they contribute functional tracks to the post-training corpus the annotation trunk was trained on: *Homo sapiens, Arabidopsis thaliana* and *Drosophila melanogaster*. Every test species is therefore held out at every stage of the pipeline.

For **mammals**, 28 training assemblies totaling 71.0 Gb: 27 species of the Tiberius mammalian training panel, 15 of them at a more recent assembly, plus *Homo sapiens* (human), validated in *Panthera pardus* (leopard) and *Mus pahari*, tested in *Bos taurus* (cow) and *Delphinapterus leucas* (beluga whale).

For **plants**, 53 training assemblies totaling 77.3 Gb, 34 species of the Tiberius angiosperm panel together with 19 of our own, validated on *Populus trichocarpa* (poplar) and *Typha latifolia* (cattail), tested on *Brachypodium distachyon, Eschscholzia californica, Freycinetia multiflora, Medicago truncatula*, and *Mimulus guttatus*.

For **insects**, the 93 BRAKER-annotated assemblies of the Tiberius Insecta panel totaling 41.9 Gb, validated on six species of our choosing, drawn from the wider BRAKER pool to reflect the phylogeny of the test species: *Anopheles coluzzii, Autographa gamma, Harmonia axyridis, Notodonta ziczac, Plutella xylostella* and *Polistes fuscatus*. The model is tested on *Bombyx mori, Cataglyphis hispanica, Colias croceus, Danaus plexippus, Leptidea sinapis, Nymphalis io, Osmia bicornis bicornis, Tribolium castaneum, Vanessa cardui* and *Zerene cesonia*. This is the only clade whose training labels are the baseline’s own BRAKER predictions rather than a reference set we assembled, so it inherits the annotation construction as well as the species panel.

#### 4.4.3. Evaluation

##### Metrics

We evaluated predicted coding structures at the CDS-exon and gene levels over the whole genome, excluding untranslated regions, similar to Tiberius [28]. At the CDS-exon level, an exon is an exact match when contig, start, end, strand, and phase all agree with the reference. At the gene level, a prediction is an exact match only when the complete set of CDS exons defining the locus matches the reference. Precision, recall, and F1 are computed as

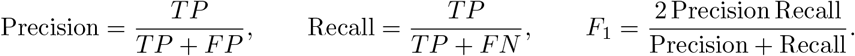

The CDS-exon quantity corresponds to the exon level of the evaluator described below and should not be confused with UTR-inclusive exon metrics. Results are reported per species and summarized as an unweighted mean across the species of each clade.

##### Scoring

Every caller is scored by one evaluator under one set of conventions. The reference is restricted to the genes retained by the same filter applied when building training targets, which removes genes with conflicting or overlapping CDS, so that predictions derived from those targets are scored on equal footing. Contig scope is matched per species rather than per caller: each species is scored on its whole assembly, applied identically to the reference and to every prediction, so a prediction on a scaffold the reference does not annotate counts against precision. Predicted genes are required to carry at least 201 bp of CDS, the threshold Tiberius applies to its own predictions [28]. Pseudogene filtering is enabled and follows BRAKER’s FilterPseudo semantics: pseudogene regions overlapping reference CDS are removed first, predicted CDS intervals overlapping the remaining pseudogene regions are then deleted, and a predicted gene disappears only if all of its CDS intervals are removed. The pseudogene source is each assembly’s own reference annotation, the same file its predictions are scored against, so the filter applies uniformly across clades and is a no-op for any assembly whose reference records no pseudogenes. Both provider conventions are recognized: a top-level pseudogene feature and a pseudogene biotype attribute on a generic gene row.

##### Isoform matching

Because reference annotations can carry several transcript structures per locus, we report both longest-isoform and any-isoform matching for every caller (see Supplementary Table C.3), with the former as our primary evaluation. Under the longest-isoform rule a prediction is compared with the single longest-CDS transcript the filtered reference retains. This is the stricter of the two conventions in use, and it charges a caller for recovering a real but non-canonical isoform. The scored loci, the predictions and the contigs are identical under both rules; only the set of structures that counts as a match changes. For the any-isoform rule we read the alternative transcripts from the multi-isoform annotation the reference was derived from, pinned to the release named in the reference’s own header, and accept a locus if the prediction reproduces any transcript that annotation assigns to that gene. This is the rule used by the Tiberius clade preprint [29], which computes it with gffcompare (`--strict-match -e 3`). Applying that protocol to our in-house Tiberius predictions reproduces their published gene-level F1 to within 0.13 points for all five plants and ten insects (Supplementary Fig. A.6b). An exact-match gffcompare run (`--strict-match -e 0 --no-merge`) reproduces our own scorer to within 0.04 points under the longest-isoform rule. Under the any-isoform rule our scorer additionally credits exact matches to alternative isoforms that share an intron chain but differ in their coding start, which gffcompare compares with a single isoform per chain; this raises our any-isoform values by up to 0.4 points on the three Phytozome-annotated plants (*E. californica, F. multiflora* and *M. guttatus*).

Before using a parent annotation we require it to reproduce at least 99% of the single-transcript reference’s gene structures exactly, which rejects a neighboring release of the same assembly; sequence names are aligned between the two by shared CDS intervals rather than by name, since the two sources do not always agree on naming. At CDS-exon level the rule is applied by scoring against the union of all accepted transcripts’ exons, following `gffcompare`. Because this enlarges the reference exon set by 4–11%, CDS-exon F1 does not rise with the gene-level score but falls, by 0.4 to 3.0 points across the callers and species we measure, so the two levels should be read together. The rule is reported for all 17 test assemblies; for the three Phytozome-annotated plants the multi-isoform annotation is the one distributed with the Tiberius training data release [86].

##### Tiberius

Tiberius was run in-house for all three clades, in each case from its released weights and with its HMM head in the default, never-trained configuration: the soft-masking mammalian configuration `mammalia_softmasking_v2` under Tiberius v2.0.7 on the soft-masked whole-genome FASTA, the unmasked `angiosperms` configuration for plant clade (weights v1.1.6) on uppercased input, as its own protocol specifies, and the unmasked insect configuration `insecta` (upstream `model_cfg/insecta.yaml`, weights v1.1.7, `insecta_unmasked_v2`). Scoring our in-house predictions and Tiberius’s own released predictions for the same mammal assemblies agrees to within 0.04 gene-level F1, confirming that the baseline was run as intended. We report Tiberius from these runs rather than quoting its published accuracies, which follow different conventions: the original study [28] evaluates cow on a different assembly, filters pseudogene loci and scores beluga whale on the scaffolds its prediction covers, and the clade study [29] matches any annotated isoform with `gffcompare` and reports its mammals from the `mammalia_nosofttmasking_v2` model (name as distributed).

##### Helixer

Helixer was run in-house from its published Docker image, one job per species, with whole-genome inference. Plants used its `land-plant` lineage and insects its invertebrate lineage (`invertebrate_v0.3_m_ 0100.h5`). Helixer publishes no mammal lineage but does release a mammal model, so for the two mammals we passed `mammal_v0.3_m_0100.h5` explicitly with `--model-filepath` while retaining the vertebrate lineage flag, which is what supplies the subsequence length of 213,840 bp.

##### GeneCAD

We additionally compared against GeneCAD, a sequence-only annotator that couples a plant DNA foundation model with a transformer head and a chromosome-scale conditional random field. Predictions were generated in-house with the released pipeline (v0.3.0) in its plant configuration, pairing the PlantCAD2 base encoder `emarro/pcad2-200M-cnet-baseline` with the released head `plantcad/genecad_plant`, both retrieved from Hugging Face at run time at revisions 5574ff5d (2026-03-11) and 9fb642bb (2026-07-03) respectively. Each assembly was annotated one chromosome at a time from the same soft-masked sequence used by our own models, at the pipeline’s default 8,192 bp window and 4,096 bp stride, batch size 56, on a single NVIDIA H100 80-GB GPU, with frame-aware decoding, the split-gene merge and the ReelProtein screen left at their released defaults. Predictions were scored with the same evaluator, reference annotation and filters as every other caller in this benchmark, over each assembly’s nuclear contigs; GeneCAD is released for plants only: its animal configuration substitutes a different foundation model and reports no gene-level accuracy. Therefore it appears in the plant comparison only, and we do not reuse the figures from its own publication, whose transcript level is UTR-inclusive and whose test species are not ours.

##### Test species in baseline training data

Some test species were used in developing the baselines. Helixer’s land-plant model was trained on *B. distachyon, M. truncatula* and *M. guttatus* (earlier assembly versions), and its invertebrate model on *B. mori*, with *O. bicornis* and *Z. cesonia* used for validation [27]. None of our test species is among GeneCAD’s training or validation species [32], and none was used to train Tiberius. This overlap can only favor Helixer, which NTv3 nonetheless outperformed on each of these six species (Supplementary Table C.3).

##### Protein-level analyses

We assessed whether predicted coding structures translate into plausible proteins, using protein-identity recovery, open-reading-frame (ORF) validity, intron-chain matching and BUSCO completeness. Proteins were extracted identically from every GFF3, predicted and reference alike, by splicing and translating each gene’s CDS (Supplementary Note B.6.2). Protein-identity recovery is the best global identity between each reference protein and any predicted protein overlapping its locus on the same strand, normalized by the length of the longer of the two. A predicted ORF is valid when it starts with methionine and has no internal stop codon; intron-chain matching requires every internal splice boundary to match the reference while ignoring the outermost transcript coordinates.

BUSCO v5.7.1 was run in protein mode against `mammalia_odb10` (9,226 markers) on the cow and beluga whale proteomes of both callers and on the unfiltered reference proteome, built from the complete reference annotation before scoring-time filtering, which serves as the completeness ceiling. All proteomes cover the same contig set. Marker counts and completeness values are given in Supplementary Note B.6.2.

We classified each reference locus into four mutually exclusive structural outcomes: correct, where one prediction exactly matches the reference structure; boundary, where exactly one prediction overlaps the locus on the same strand but has an incorrect structure; missed, where no prediction overlaps the locus on the same strand; and fragmented, where two or more predictions overlap the locus on the same strand and none matches it exactly. Overlap is tested between individual coding exons rather than between gene envelopes, so a prediction lying within an intron of the locus does not overlap it. These four classes partition the reference. Predictions with no corresponding reference locus were counted separately, as invented predictions, since they belong to prediction-side precision rather than to any reference class.

For exon-position analyses, coding exons were indexed from 5^′^ to 3^′^ along the transcript, so that the first exon denotes the biological first coding exon on both strands. For failed exon matches, we measured the magnitude of coordinate shifts and the fraction of shifts that are multiples of three; a fraction near one third is expected under a null in which shifts are independent of coding phase, whereas an excess indicates the decoder selecting a different in-frame boundary. For the positional analysis of genes whose exact-match failure involved exactly one incorrect exon, a separate null was generated by sampling an exon uniformly from the exons of that same failed reference gene.

The primary longest-isoform rule can classify a biologically valid alternative isoform as an error. As a control, we found that 27.7% of loci classified as incorrect under that rule in cow (25.7% in beluga whale) exactly match another annotated isoform. Among those loci the second-longest isoform was selected in 63% of cases, with a median predicted-to-longest CDS-length ratio of 0.957, and the matched isoform lies within one coding exon of the longest in about 90% of cases, indicating a terminal-boundary difference rather than a wrong gene call. After allowing the best-matching annotated isoform, the remaining errors in cow comprise 43% internal-boundary errors, 29% missed loci and 26% terminal-boundary-only errors (50%, 26% and 23% in beluga whale). These analyses use the primary longest-isoform rule. Recomputing them against the best-matching annotated isoform strengthens rather than weakens the first-exon result: the first-coding-exon enrichment among one-exon errors rises from 2.73 to 3.10 in cow and from 2.58 to 3.07 in beluga whale, closely tracking Tiberius under both rules (2.72 to 3.03 and 2.73 to 3.25). The same per-class analysis was repeated against the best-matching annotated isoform: all three multi-exon classes fall by 21 to 25% (first coding exons from 26.9% to 20.3%, last from 20.4% to 16.2%, internal from 9.5% to 7.5%), so first coding exons remain the least accurately predicted multi-exon class and their excess over internal exons is unchanged (2.8-fold against 2.7-fold). Single-exon genes fall by under 2%, from 26.2% to 25.7%, and become the least accurately predicted class overall once alternative isoforms are allowed. The concentration of errors at the first coding exon is therefore not an artifact of the longest-isoform reference.

For the analysis by gene length (Fig. 4f), the span of a reference locus is the envelope of the coding sequence of its retained isoform, from its first to its last coding base, so untranslated regions are excluded. Loci were binned by span into half-open intervals with edges at 5, 20, 50, 131 and 500 kb. A locus counts as recovered when it is classified correct, and as fragmented as defined above. Predicted genes with less than 201 bp of coding sequence were discarded before classification, for both callers. Pooled over cow and beluga whale, the six bins hold 12,066, 11,475, 7,772, 5,512, 2,754 and 387 reference loci.

The thirteen structural aspects of Fig. 4e comprise six gene-level and seven exon-level measures. At the gene level, they are recall, precision and F1, the fractions of reference loci not missed and not fragmented, and the fraction of predicted genes overlapping a reference locus on their strand. At the exon level, they are CDS-exon recall, precision and F1, the fractions of reference coding exons predicted exactly and not missed, the fraction of predicted coding exons overlapping a reference coding exon, and the fraction of first coding exons predicted exactly. Recall, precision and F1 at both levels are the exact-match measures reported in the Tiberius benchmarks [28, 29]; the remaining seven measures describe failure modes. Each is a percentage oriented so that higher is better. Both callers were scored by the same implementation, against the same filtered longest-isoform reference, over the whole assembly and with the same 201-bp minimum on predictions; the gene- and CDS-level F1 recomputed in this analysis reproduce the headline values for every species.

##### Throughput benchmark

Each caller was timed end to end on one NVIDIA H100 80-GB GPU, from a soft-masked FASTA on local disk to its annotation written to local disk, including model loading, compilation, inference, decoding and file writing, and excluding repeat masking and evaluation. NTv3 ran with bfloat16 compute at *ρ* = 0.75, which is 1.7 times faster than FP32 and changes gene- and CDS-level F1 by at most 0.03 on cow. Each runtime comes from a single run. Seconds per Mb are reported as unweighted means over assemblies of wall-clock time divided by assembly length, separately for the 13 assemblies whose mean contig exceeds the 1-Mb input window and for the four fragmented ones.

NTv3’s runtime is affine in the number of input windows *W*, *t* = *a* + *bW*, with *a* = 80 s, most of it a 58-s compilation, and *b* = 0.74 s per window (*r*^2^ > 0.999 over the 17 assemblies). Because every contig is padded to whole windows, a fragmented assembly costs more per base than a contiguous one of the same size (Supplementary Note B.6.3).

##### Statistical reporting

Annotation models were trained with one random seed per training arm, and no seed replicates were performed for the annotation experiments. The evaluation therefore comprises *n* = 2 mammalian, *n* = 5 plant, and *n* = 10 insect test species rather than independent replicate trainings, differing from the three-seed protocol used for the functional-track benchmark. Differences smaller than the effects of region selection, checkpoint selection, or evaluation conventions are single-seed estimates and should not be over-interpreted. Consistency across species is assessed with a two-sided sign test over the 17 test species, each species counting as one observation.

### 4.5 Human Functional-Track Benchmark Comparisons

#### 4.5.1. Benchmarking Against Borzoi Track Predictions

NTv3 was compared with the released Borzoi model [5] in two settings, as done previously by AlphaGenome [6]. First, we evaluated predictions against the same 1-bp target tracks while retaining each model’s native output grid. Second, we fine-tuned NTv3 on the released Borzoi training set and compared the models at Borzoi’s native 32-bp resolution. Both settings used Borzoi replicate 0 folds: fold 4 for model selection and fold 3 for final evaluation. Since AlphaGenome reported performance on Borzoi tracks for fold-4 (Borzoi replicate 1 test set) [6], we report NTv3 performance also on fold-4 to compare with AlphaGenome. We note that this is acceptable because NTv3 performance is similar between both fold-3 and -4.

For the 1 bp comparison, we inverted Borzoi’s assay-specific target transformations using the scaling factors, aggregation functions and clipping parameters distributed with its data release. Each 32-bp Borzoi prediction was expanded by nearest-neighbor repetition over the corresponding 32 nucleotide positions. This aligns the predictions to the 1-bp target grid but does not create information below Borzoi’s native 32-bp resolution. For input length *L*, consecutive windows were stepped by the central prediction length, 0.375*L*, giving 62.5% overlap and covering each evaluated nucleotide once after cropping. We evaluated the intersection of target tracks available to both models. GTEx-derived tracks were excluded from this comparison because AlphaGenome does not report per-track performance for them, so no matched baseline exists; per-track values were published only for the Borzoi target set. After inverse transformation, predictions and targets were transformed with log(1 + *x*). Pearson correlation was calculated separately for each track across the complete fold evaluation set and then averaged within assay classes.

For the matched 32-bp comparison, we converted the released Borzoi TFRecords to TensorStore without altering sequences, targets or fold assignments. We replaced the NTv3 prediction head with a randomly initialized head spanning the human (hg38) Borzoi target set of 7,611 tracks, and fine-tuned all model parameters. NTv3 was compared with both Borzoi and AlphaGenome on matched folds.

#### 4.5.2. Benchmarking Against AlphaGenome Track Predictions

NTv3 was compared with AlphaGenome [6] on AlphaGenome’s own released training data. We used Al-phaGenome’s tasks released at single-base resolution: ATAC-seq, DNase-seq, CAGE, PRO-cap and RNA-seq. We excluded the two ChIP-seq tasks because they are binned at 128 bp rather than 1 bp, and the splice tasks since they are pairwise quantities that our architecture does not predict. GTEx RNA-seq data were not available from AlphaGenome and were excluded.

We fine-tuned one model per assay, each predicting only the tracks of that assay, rather than a single model spanning all five. All five were warm-started from the same NTv3 checkpoint and trained under an identical protocol, so they differ only in which assay they saw. Each replaced the NTv3 prediction head with a randomly initialized head over its assay’s tracks, conditioned on a learned species embedding, and fine-tuned all model parameters at 1,048,576 bp inputs with reverse-complement augmentation. Training used the same Poisson-multinomial objective as the Borzoi fine-tune, with no masked-language auxiliary loss and no genome-annotation head.

Models were evaluated on AlphaGenome’s own held-out split, the 6,853 windows of 196,608 bp it reports on, centered in the longer input, with predictions averaged over the forward strand and its reverse complement. AlphaGenome reuses Borzoi’s eight folds but exchanges the two held-out ones, so this split corresponds to Borzoi fold 4, which NTv3 holds out as its validation split; no gradient of any model reported here saw it. For DNase-seq, whose signal records cut sites on the boundary at the start of each base, the reverse-complement read is shifted back by one base before averaging.

Pearson correlation was calculated separately for each track across the full evaluation set and averaged without weighting within each assay. Each assay is scored on the transform AlphaGenome reports it on: untransformed for the two accessibility assays, log(1 + *x*) for CAGE, PRO-cap and RNA-seq. NTv3 is compared against the per-assay correlations reported by AlphaGenome [6].

### 4.6 NTv3 Benchmark

We constructed NTv3 Benchmark to compare sequence representations on functional-track prediction across eleven species under a common supervised adaptation protocol. For species represented during NTv3 post-training, the benchmark included only experiments absent from the post-training target corpus. For species absent from functional-track post-training, all benchmark tracks were new supervised targets. Here, “unseen species” therefore means unseen during functional-track supervision, although the genome, or genomes of related taxa, is present in the OpenGenome2 self-supervised corpus.

The benchmark comprises 155 held-out functional tracks over those eleven species, spanning chromatin accessibility, histone and transcription-factor ChIP-seq, transcription initiation, RNA binding, transcription and translation. The species, assemblies, assays and track counts are listed in Table 7, and the data sources and processing for each species are given in Supplementary Note B.5 and Supplementary Table C.4.

**Table 7.** The NTv3 Benchmark: eleven species across four clades, totaling 155 held-out functional tracks. Six species are new relative to the first version of the benchmark, which extended it from mammals and plants into fungi and protists. For species present in NTv3 post-training, only assays absent from the post-training corpus were included.

| Species | Assembly | Assays | Tracks |
| --- | --- | --- | --- |
| *Mammals* |  |  |  |
| Human | hg38 | ATAC-seq, histone ChIP-seq, PRO-cap, eCLIP, polyA<sup>+</sup> and total RNA-seq | 34 |
| Cattle | ARS-UCD1.2 | ATAC-seq, histone ChIP-seq, TF ChIP-seq, RNA-seq | 30 |
| Rat | mRatBN7 | ATAC-seq, RNA-seq | 12 |
| *Plants* |  |  |  |
| *A. thaliana* | TAIR10 | Ribo-seq | 4 |
| Rice | IRGSP-1.0 | Ribo-seq | 14 |
| Maize | Zm-B73-REFERENCE-NAM-5.0 | Ribo-seq | 8 |
| Tomato | SL3 | ATAC-seq, RNA-seq | 20 |
| *Fungi* |  |  |  |
| *S. cerevisiae* | R64 | ATAC-seq, RNA-seq | 12 |
| *S. pombe* | ASM294v2 | ATAC-seq, RNA-seq | 7 |
| *Protists* |  |  |  |
| *P. falciparum* | GCA000002765v3 | ATAC-seq, RNA-seq | 7 |
| *D. discoideum* | dicty_2.7 | RNA-seq | 7 |
| **Total** |  |  | **155** |

#### 4.6.1. Models and Training Protocol

We evaluated nine baselines against NTv3. Two are convolutional models trained from random initialization on each task: a 6-million-parameter BPNet [4] using the scaled backbone configuration of ChromBPNet [87], and a 44-million-parameter dilated residual convolutional network following the SpliceAI architecture [10]. The smaller variants of both families were not included, because they cover only two of the eleven species. Five are released self-supervised models: Nucleotide Transformer v2 [19], HyenaDNA [20], Caduceus [24], PlantCAD2 and Evo 2 [25]. The last two are Borzoi [5] and AlphaGenome [6], sequence-to-function models trained on human and mouse functional data.

Coverage is deliberately uneven, because the released models differ in taxonomic scope and in the compute their fine-tuning demands. BPNet, the residual convolutional network, Borzoi, AlphaGenome and NTv3 were evaluated on all eleven species. Nucleotide Transformer v2 and Evo 2 were evaluated on five (human, *A. thaliana*, rice, maize and tomato), PlantCAD2 only on plants, and HyenaDNA and Caduceus only on human since they are pre-trained on human data. AlphaGenome fine-tuning was performed by the Peter Koo Lab at Cold Spring Harbor Laboratory.

For the published models, we used the following checkpoints:

- `InstaDeepAI/nucleotide-transformer-v2-500m-multi-species`;
- `LongSafari/hyenadna-large-1m-seqlen-hf`;
- `kuleshov-group/caduceus-ph_seqlen-131k_d_model-256_n_layer-16`;
- `kuleshov-group/PlantCAD2-Small-l24-d0768`;
- `evo2/1b-8k-bf16:1.0`; and
- Borzoi released replicate 0.
- AlphaGenome fold 1 for human (since its distilled all-folds version saw the human test regions used in this benchmark) and the distilled all-folds model for all other 10 species.

The selected Evo 2 checkpoint has an 8,192-token context and was pre-trained by autoregressive next-nucleotide prediction; it is distinct from the 1-Mb Evo 2 model.

Every model received the same 32,768-bp input sequence and produced 1-bp predictions for the central 12,288 positions. Several of the released models were therefore evaluated beyond their pre-training context length, which is a property of the comparison rather than a defect of the protocol: the window, the scored region, the training objective and the optimization budget are held fixed across models. Architecture, parameter count, native output resolution, tokenization and, for Evo 2, the optimizer and numerical configuration necessarily differ, so the comparison evaluates the pre-trained representations under a common adaptation protocol, although it does not make the representation the only variable. A layer-normalization, linear-projection and softplus head mapped nucleotide representations to non-negative task targets. One multi-output head was trained per species–dataset combination, and all tracks in that task were optimized jointly. For pre-trained models, the released prediction head was replaced and all model parameters were updated; convolutional baselines were trained from random initialization on the same examples. For NTv3, fine-tuning was initialized from the post-trained tracks NTv3 model.

##### Optimization and evaluation

Models were trained for ten epochs over each species-specific training partition, and the validation Pearson correlation was measured on up to 1,000 randomly drawn validation windows, twenty times per epoch. We selected the checkpoint maximizing mean validation Pearson correlation independently for each random seed; no early stopping was applied. Every model was run with three seeds on every species it covers, with the exception of Evo 2, whose fine-tuning cost restricted it to a single seed, so no error bar is reported for it. Each selected checkpoint was evaluated once on the held-out test partition, and reported values are means across seeds with error bars giving the standard error of that mean. Pearson correlation was calculated separately for each track over all evaluated positions and then averaged across tracks; correlations are computed in the transformed target space described in Section 4.3 rather than on raw signal.

All models used an effective batch size of 32, implemented by gradient accumulation where necessary. Except for Evo 2, optimization used AdamW with *β*_1_ = 0.9, *β*_2_ = 0.999 and weight decay 0.01. The learning rate increased linearly from 10^−5^ to 5 × 10^−5^ over the first 3% of training nucleotide tokens and subsequently followed the polynomial-decay schedule used for NTv3 track post-training. From-scratch BPNet and SpliceAI models instead used a peak learning rate of 10^−3^. Evo 2 was trained with the numerical configuration required by `bionemo-framework`: Adam, BF16 parameter storage, FP32 gradient accumulation and gradient clipping at a norm of 1.0. The JAX implementations of NTv3 and the convolutional baselines used data parallelism; other released models used PyTorch with DeepSpeed ZeRO-2 and `bfloat16` precision.

All functional targets underwent the transform in Section 4.3. Every model was optimized with the same Poisson–multinomial objective, computed only over the central 37.5% of each input.

### 4.7 External Multispecies Tracks Benchmarks

#### 4.7.1. SeiPlant

SeiPlant [36] adapts the Sei architecture [13] to plant genomes, predicting seven histone modifications from 1,024 bp of sequence. Its authors release family-level checkpoints trained on Brassicaceae and on Poaceae, together with a model trained on both. We used this benchmark to ask whether NTv3 transfers to plant species that neither model has seen, and we evaluated both models zero-shot: no parameter of either was updated, and NTv3 was given no new prediction head.

The evaluation covers five species absent from NTv3 post-training and from SeiPlant’s training set: *Brassica napus, Arabis alpina, Arabidopsis lyrata, Hordeum vulgare* and *Brachypodium distachyon*. Target signal and peak calls were obtained from ChIP-Hub (https://biobigdata.nju.edu.cn/ChIPHub_download) for the 23 experiments these species contribute. We restricted the comparison to *H3K4me3*, the one mark for which both models carry a directly comparable output. Because the two released SeiPlant checkpoints differ in which family they were trained on, each species is scored against both: a *within-family* arm, whose training family contains the test species, and a *cross-family* arm, whose training family does not.

##### Evaluation procedure

SeiPlant writes one value per 1,024 bp window at the central 128 bp of that window, and we scored exactly that interval for both models rather than the full window, since correlating a 1,024 bp average against a 128 bp one would compare two different quantities. NTv3 was run at a 1 Mb input with the standard 37.5% central crop and its base-resolution predictions averaged over the same 128 bp bin, after inverting the training-time target transform so that the average is taken in physical signal units. Predictions were made on the forward strand only, matching SeiPlant, which uses no reverse-complement augmentation. None of the five species is represented in NTv3’ species vocabulary or output heads, so each was scored by borrowing the conditioning token and the functional-track head of a training species, averaging over that head’s H3K4me3 columns. Here, the *A. thaliana* token and head were used for all five species. Windows containing any non-ACGT base were excluded, using the same mask for both models. Performance is Pearson correlation between predicted and observed signal over the scored bins, reduced as the median across chromosomes within an experiment and then the median across experiments, so that species contributing many experiments are not dominated by whichever experiment has the most chromosomes.

#### 4.7.2. Gene-level Expression in Plants

Plant data on gene expression was based on that compiled for the Plant Genomic Benchmark presented in the Agronomic Nucleotide Transformer (AgroNT) study [37]. Unlike the functional track prediction tasks, this task aims to predict gene expression profiles across tissues at the gene level, i.e., producing predictions per gene rather than per base pair. Briefly, the dataset was composed of bulk RNA-seq data from four plant species, namely *A. thaliana, O. sativa, S. lycopersicum*, and *Z. mays*, across a range of organs and developmental stages. Unlike the original study, which used DESeq2-normalized counts, our downstream task relied on transcripts per million (TPM). To compute TPM values, we used the same pre-computed gene counts and applied gene length correction based on collapsed exon lengths for each gene. TPM values were then log-transformed using a pseudo-count of 1 to avoid taking the logarithm of zero.

For model training, we extracted promoter-proximal sequences centered on annotated transcription start sites (TSS). To evaluate the benefit of providing a larger genomic context, we constructed three versions of each promoter sequence by extending the flanking region symmetrically around the TSS by 2,944, 16,000 and 65,536 bp. Since the flank is applied on both sides, the corresponding total input lengths are 5,888 bp, 32,000 bp and 131,072 bp; below we refer to each version by its total length, rounded, as 6 kb, 32 kb and 131 kb. Only genes successfully extracted at every flank size were retained, so that all context sizes are evaluated on an identical gene set. To prevent data leakage, genes were assigned using the same homology-based splits described above, in which a target combined validation and test size in base pairs is set per assembly and the remaining sequence forms the training set.

##### Evaluation procedure

NTv3 models were compared with the AgroNT 1-billion-parameter model [37] on these tasks. Training used an effective batch size of 16, the Adam optimizer with a fixed learning rate of 5 × 10^−5^ and no weight decay. Training proceeded for up to 200,000 update steps with early stopping based on validation performance. Validation was performed every 2,000 steps on the complete validation set. Early stopping was triggered if validation performance did not improve for 10 consecutive validation evaluations. All experiments used a fixed random seed of 0 for reproducibility. Final model evaluation was conducted on the held-out test set using the checkpoint with the best validation performance.

#### 4.7.3. Gene-level Protein Abundance in Plants

Plant protein abundance data were obtained from Ribo-seq assays using the dataset and pipeline described above. As in the gene-level expression task, protein abundance was formulated as a gene-level prediction problem. Specifically, protein abundance values were derived using StringTie [88] with the options `-e -A`. These options enabled gene-level quantification in TPM from the high-quality alignments and produced per-gene abundance estimates based on the reference annotation, providing a proxy for protein synthesis. Models were fine-tuned using a regression objective, with input sequences constructed using the same TSS-centered flanking regions used for the gene expression task. Protein abundance prediction was evaluated across the same set of total input lengths: 6 kb, 32 kb and 131 kb.

##### Evaluation procedure

Same as above for the gene-level expression task.

#### 4.7.4. DeepFARM

DeepFARM [39] models chromatin activity in farmed animal genomes, framing prediction as multi-label binary classification of 200 bp bins against published peak calls. Its authors release one DanQ [38] checkpoint per species, trained on cattle, pig and chicken. We evaluated NTv3 against those released checkpoints on chicken and pig, the species present in our NTv3 training data.

Peak calls were retrieved from the Gene Expression Omnibus under SuperSeries GSE158430, covering ATAC-seq, CTCF, DNase-seq, H3K27ac, H3K27me3, H3K4me1 and H3K4me3; reference genomes were taken from Ensembl release 104. Following DeepFARM, a bin is labeled positive for a track when a single peak covers at least half of it, and bins tile each chromosome from the first base with any trailing partial bin discarded. DanQ receives 1,000 bp centered on the scored bin, so its first and last two bins per chromosome are not predicted.

##### Evaluation procedure

Neither model was trained or fine-tuned for this benchmark. DanQ was run from the released per-species weights, and NTv3 from its released checkpoint, conditioned on the species token for the assembly being scored and read through its existing functional-track head. Base-resolution predictions were written into a chromosome-length buffer and binned once on the same global 200 bp grid used for the labels, a bin being emitted only when all 200 of its bases were predicted; the bin value is the mean of those predictions. Input windows were required to fall entirely within the chromosome rather than padded, so that no prediction rests on fabricated sequence.

The two models cover different tracks and different bins, and the comparison is restricted to the intersection of both. NTv3 predicts only those tracks that pair with one of its head columns under a mapping fixed before any inference was run, which gives 30 of 97 chicken tracks and 16 of 80 pig tracks; DanQ is then scored on that same subset rather than on its full output. Bins that NTv3 cannot predict, at the leading and trailing ends of each chromosome, are removed from DanQ as well. Performance is auROC per track, computed with one metric implementation for both models, and compared as a paired per-track difference. Confidence intervals come from a block bootstrap over contiguous 25 kb blocks, which respects genomic autocorrelation; a track is counted as improved only when its interval excludes zero.

The NTv3 trunk saw FAANG coverage of the same assays and tissues, though from different studies: the peak calls derive from the UC Davis FAANG pilot, whereas our tracks come from separate European FAANG submissions. Both pig studies are late embryonic, so pig coverage is compared against adult peak calls. What makes the comparison informative is therefore not the conditioning but the region restriction: scoring is confined to bins whose entire input window lies inside sequence held out of NTv3’ training splits, so no prediction can draw on a region the model was trained on. A secondary analysis restricts only the scored bin and not its context. Before any NTv3 inference was run, we reproduced DeepFARM’s own published per-track results from their released weights as a precondition, so that a fault in our harness could be distinguished from a real difference between the models.

#### 4.7.5. DanioDecima

DanioDecima [40] predicts zebrafish gene expression from sequence across 304 single-cell pseudobulks spanning ten developmental stages, using a Borzoi trunk [5] with a global average pool and a linear head. Unlike the benchmarks above, this one is a fine-tuning comparison rather than a zero-shot one: the question is whether a multispecies functional trunk is a better starting point for a zebrafish expression model than a human or mouse one.

Pseudobulk expression was assembled from the publicly released per-stage ZebraHub atlas files, summing raw counts within each fine ontology class in each stage and converting to log(1 + CPM). Sequences were taken from GRCz11 with Ensembl release 116 annotation. We adopted DanioDecima’s task definition unchanged: each gene is represented by a 524,288 bp window *anchored* on the gene start rather than centered on it, with 163,840 bp upstream, reverse complemented for genes on the minus strand, and a binary mask marking the gene body. Predictions are the mean of the model’s output over the 196,608 bp beginning at the gene start, which is the extent their global pool corresponds to. Training used their task-wise Poisson–multinomial objective, in which the multinomial term runs along the pseudobulk axis rather than along sequence positions. We used their gene splits, defined by held-out chromosomes, giving 18,368 training, 6,110 validation and 7,289 test genes out of 31,767.

##### Evaluation procedure

NTv3 was fine-tuned from the post-trained checkpoint with the species token for zebrafish, for up to 40 epochs at a learning rate of 3 × 10^−5^ and an effective batch size of 20, selecting the checkpoint by validation performance. Performance is reported two ways, both as the mean over columns of a column-wise Pearson correlation: *per gene*, correlating across the 304 pseudobulks within each gene and averaging over genes, and *per pseudobulk*, correlating across genes within each pseudobulk and averaging over the 304.

Because the targets are pseudobulks over as few as a few dozen cells, they carry sampling noise that bounds any predictor. We therefore also estimated a label reliability ceiling by splitting the cells of each pseudobulk into random halves, recomputing each half’s log(1 + CPM) independently, correlating the two replicates under both reductions, and applying a Spearman–Brown correction to project the split-half value to full sampling depth.

### 4.8 Model Interpretation Analysis

To understand the regulatory mechanisms captured by NTv3, we used three complementary model readouts: (i) predicted functional tracks, which report assay-specific regulatory activity at base-pair resolution; (ii) gradient-based attribution maps, which identify the sequence positions most influential for a selected model prediction; and (iii) transformer attention maps, which summarize how the model integrates information across long genomic distances at a resolution determined by the downsampling architecture. Together with controlled sequence perturbations, these readouts support multiscale interpretation from local sequence features to distal regulatory interactions. All interpretability results in Fig. 5 were obtained using the single-length NTv3 model on 1,048,576-bp inputs; attention maps have a resolution of 128 bp per token, corresponding to seven downsampling stages (1,048,576 bp input ÷ 8,192 tokens).

#### 4.8.1. Interpretability Methods

##### Predicted Functional Tracks

Functional track predictions are cell-type and assay specific. For each input sequence, the functional-track module produced per-base predictions for all species-matched regulatory assays. Consistent with the post-training configuration, supervision for functional tracks is applied only to the central 37.5% of the input sequence. Accordingly, all analyses used predictions extracted from this supervised region to ensure that every interpreted position had full receptive-field coverage.

Functional-track predictions were obtained directly from the model output tensor of shape `(sequence length, number of tracks)` without additional smoothing or normalization. For visualization (e.g., Fig. 5), predictions for assays such as K562 DNase-seq, K562 GATA1 ChIP-seq, K562 CAGE-seq, and GTEx RNA-seq were retrieved by indexing into this tensor using the track metadata.

##### Gradient-Based Attribution Maps

Attribution maps are cell-type specific, as they are computed with respect to a chosen functional track. Gradient-based attribution provides a first-order, base-pair-resolved estimate of how NTv3’s predictions depend on the underlying DNA sequence. We employ gradient-based saliency maps [89], which quantify the local sensitivity of a predicted regulatory signal to infinitesimal changes at each input base. In regulatory genomics this is well motivated: TFBSs, splice junctions, and other *cis*-regulatory elements often depend on a small number of informative nucleotides, such that single-base substitutions can meaningfully alter regulatory output. As a result, large-magnitude gradients arise at positions with high predictive influence, enabling identification of the motif-level sequence features used by the model.

The interpreted checkpoint is the post-trained NTv3 model, which uses the 16-token soft-masked vocabulary described above. To compute these attributions, we differentiated model outputs with respect to the one-hot encoded DNA input [89, 90]. Because NTv3 represents DNA using a discrete token vocabulary, gradients cannot be taken with respect to token IDs. Instead, we used a differentiable embedding override: for a sequence of length *L*, the one-hot matrix

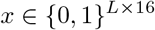

was multiplied by the learned embedding matrix to produce continuous token embeddings. These embeddings were supplied directly to the model, ensuring that *∂y*/*∂x* is well-defined and enabling gradients to flow to every position.

For each selected functional track, the model outputs logits of shape `(batch, sequence length, number of tracks)`. To form a scalar suitable for differentiation, we aggregated the logits over sequence positions and (if applicable) over a subset of tracks, followed by a log transform for numerical stability:

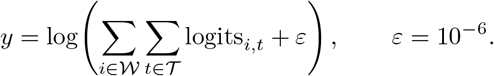

Here, *W* denotes the set of sequence positions included in the supervised output window (e.g., the central 49,152 bp for functional tracks), and *T* denotes the subset of functional-track indices used for the attribution calculation.

We then computed

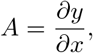

yielding an attribution tensor *A* ∈ ℝ^*L*×16^ encoding the local sensitivity of the aggregated prediction to perturbations at each input token.

For base-pair-resolution interpretation, we retained only the uppercase A/T/C/G channels (vocabulary indices 6–9), discarding the non-nucleotide channels. Note that under the soft-masked vocabulary the lowercase tokens at indices 11–14 represent the same four bases in repeat-masked intervals, so attribution over such intervals is not carried by the retained channels. These channel-resolved hypothetical gradients correspond to counterfactual substitutions at each base and form the basis for motif discovery, local sequence-grammar analysis, and allele-specific interpretation.

##### Transformer Attention Maps

Attention maps are cell-type agnostic, reflecting the model’s internal context aggregation independent of any specific output track. Transformer self-attention provides a complementary interpretability modality by revealing how NTv3 integrates information across distal regions of the input sequence. For genomic inputs, attention scores measure how sequence information at one downsampled token (128 bp per token; see above) influences the contextual representation of other tokens across the locus. Because these scores are computed after substantial sequence compression, they offer a coarse-grained but informative view of long-range dependencies not captured by base-resolution gradient methods.

Multi-head self-attention computes pairwise similarity between token representations via learned query– key interactions, producing attention weights that modulate how information at one position influences the contextual encoding of others. Although attention weights are not direct measures of biological causality, they provide a useful diagnostic of the long-range dependencies the model has learned. We therefore use attention maps as a post-hoc interpretability modality to examine which distal regions NTv3 emphasizes when forming its predictions.

To analyze these dependencies, we extracted multi-head self-attention weights from the final transformer block and averaged them across heads to obtain a single *L*^′^ × *L*^′^ attention matrix (with *L*^′^ ≈ 1024 for 131 kb inputs). This follows prior work demonstrating that transformer attention can highlight structured token–token interactions in language models [91] and genomic models such as Borzoi [5].

To standardize interpretation and mitigate asymmetries in directional attention, matrices were symmetrized as (*A* + *A*^⊤^)/2. Lower-triangular entries were masked, and matrices were visualized using a 45° rotated triangular layout, which highlights long-range interaction bands as pyramidal structures. For variant-centered or enhancer-focused analyses, attention was cropped to windows surrounding the locus of interest and aligned to functional-track outputs and attribution maps.

##### Interpretability Output Alignment

Functional-track predictions and attribution maps were aligned directly to genomic base-pair coordinates. Token-resolution outputs such as attention maps were aligned by converting token indices to base-pair positions using the fixed downsampling factor (128 bp per token) described above. This alignment enabled direct comparison of assay-specific signal, base-resolution sequence attribution and coarse long-range dependencies across the same locus.

#### 4.8.2. Interpretation Tasks

##### Differential Comparison Across Modalities

To compare paired reference/alternative or perturbed sequences, we computed differential profiles for the model readouts used in the interpretation analyses: functional tracks, attribution maps and attention maps. Because these outputs have distinct numerical properties, each was transformed in a modality-appropriate manner while preserving the direction and magnitude of the corresponding change.

###### Predicted functional tracks

For RNA-seq, DNase-seq, CAGE-seq, and ChIP-seq-like prediction heads, which produce non-negative per-base signals, log_2_FC was computed elementwise as

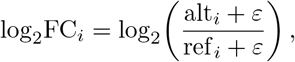

with a numerical stabilizer *ε* = 10^−6^. Positive values correspond to increased predicted activity under the alternative allele, and negative values correspond to reduced activity.

###### Gradient-based attribution maps

To compare attribution maps between reference and alternative alleles, we computed log_2_FC after shifting both matrices by their joint minimum to ensure positivity. The resulting *L* × 4 log_2_FC matrix (positions × nucleotide channels) was collapsed to a one-dimensional bidirectional profile by separately summing positive and negative values across channels at each position:

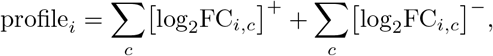

where [·]^+^ = max(·, 0) and [·]^−^ = min(·, 0). This aggregation preserves both gains and losses in attribution at each position.

###### Transformer attention maps

Let *A*^ref^ and *A*^alt^ denote symmetrized attention matrices for the reference and alternative sequences, respectively. We computed log_2_FC attention maps using

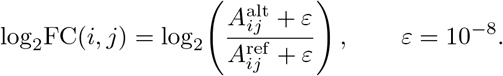

The resulting matrices highlight long-range dependencies whose strength changes between alleles, with positive values indicating gains in attention and negative values indicating attenuation.

##### Enhancer–Promoter (EP) Task

To evaluate whether NTv3 captures experimentally validated enhancer–promoter relationships, we analyzed the human *HBE1* locus, a well-characterized erythroid regulatory region containing five DNase hypersensitive elements (HS1–HS5). These enhancers have been extensively assayed in erythroid cells, and their relative strengths have been quantified using massively parallel reporter assays and CRISPR-based perturbations [14, 47]. This locus therefore provides a biologically grounded test case for assessing long-range sequence-to-function modeling.

All analyses were performed on the 1,048,576 bp input window `chr11:4,744,107–5,792,683` (hg38), which spans the *HBE1* locus. Within this window, enhancer positions were defined using the hypersensitive-site coordinates reported by Agarwal [14]:

- HS1: `5,275,890–5,276,090`
- HS2: `5,280,660–5,280,860`
- HS3: `5,284,690–5,284,890`
- HS4: `5,288,180–5,288,380`
- HS5: `5,291,330–5,291,530`

To contextualize enhancer activity, we used matched erythroid functional assays from the post-training dataset: K562 DNase-seq (ENCSR000EKN), K562 GATA1 ChIP-seq (ENCSR000EFT), K562 CAGE-seq (CNhs11250).

These provide chromatin accessibility, TF binding, and promoter-proximal transcriptional output in the same cellular context in which HS1–HS5 enhancer activities were experimentally validated.

For the HBE1 enhancer–promoter analysis, attribution maps were computed on the reference sequence with respect to the K562 CAGE track. To evaluate enhancer function, we performed *in silico* perturbation experiments in which enhancer sequences were systematically ablated and restored. Dinucleotide-preserving shuffling was used to ablate regulatory content while maintaining local sequence composition, using the HS1–HS5 enhancer coordinates defined above.

A fully ablated background sequence was first constructed by shuffling all five enhancer elements simultaneously. Enhancer sufficiency was then assessed by restoring individual enhancers one at a time while keeping the remaining enhancers shuffled. For each enhancer, five independent shuffled replicates were generated for both the fully ablated background and each single-enhancer restoration condition.

All perturbed sequences were passed through NTv3 to extract predicted K562 CAGE signal. Track outputs were restricted to the supervised central 393,216 bp region of the output sequence.

To quantify enhancer sufficiency, we measured the change in predicted promoter-proximal CAGE signal when a single enhancer was preserved relative to the fully ablated background. Specifically, we extracted a 384 bp promoter window centered at the HBE1 transcription start site (`chr11:5,269,945`; hg38; ±192 bp). Within this window, we computed the log_2_ fold-change at each base position and then averaged these values across positions to obtain a single enhancer-effect score. More positive values correspond to greater enhancer sufficiency, reflecting the ability of that element alone to rescue promoter activity above the fully ablated baseline. Predicted sufficiency scores (mean ± s.d. across five shuffle replicates) were: HS2 (0.988 ± 0.108 log_2_FC), HS4 (0.738 ± 0.013), HS1 (0.651 ± 0.050), HS3 (0.323 ± 0.079), and HS5 (− 0.005 ± 0.012). This perturbation-based sufficiency metric parallels experimental measurements reported in massively parallel enhancer assays [14].

##### Quantitative eQTL Variant-Effect Benchmark

We evaluated quantitative variant-effect prediction across 21,241 eQTLs from 48 GTEx tissues using a per-tissue classification framework. Performance was computed separately within each tissue using auROC and auPRC and summarized across tissues using tissue-weighted means. Uncertainty was estimated with 400 bootstrap resamples stratified by tissue. Pairwise model comparisons were performed on matched bootstrap resamples so that each replicate preserved the same tissue composition across model arms.

Each variant was scored zero-shot with the 1-Mb NTv3 checkpoint, on a 1,048,576-bp human input centered on the variant, using predictions over the whole input rather than only its supervised center. For every GENCODE gene contained in the input and every tissue-matched RNA-seq track (mostly three GTEx samples per tissue), we averaged the predicted signal over the gene’s exons and computed 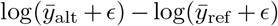 with *ϵ* = 10^−3^, averaged over the forward and reverse-complement strands. The variant score is the largest absolute value over tracks and genes, following AlphaGenome [6]. As in AlphaGenome’s variant benchmarks, variants were restricted to its zero-shot test chromosomes (3, 6, 9, 12, 16, 18, 19 and 21).

We compared NTv3 with Borzoi fold 0 and the four-replicate Borzoi ensemble. For each comparison, we report the paired difference in the tissue-weighted metric, its bootstrap interval and the corresponding paired bootstrap *P* value. AlphaGenome is included only through its published aggregate eQTL auROC; per-tissue values and eQTL auPRC were not available from the published benchmark.

##### Variant Effect Prediction (VEP) Task

To examine local regulatory responses to individual SNVs, we analyzed 3,792 variants from GTEx Whole Blood after mapping to the hg38 reference genome. Variants were excluded if the 1,048,576 bp centered window extended beyond chromosome boundaries.

GENCODE v44 [92] was used only as external locus metadata for visualization and to label transcript and splice context; these annotations were not produced by NTv3 and did not enter variant scoring. For each variant, we extracted a 1,048,576 bp window centered on the SNV and constructed matched reference and alternative input sequences differing only at the central allele.

Variant responses were evaluated using the GTEx Whole Blood RNA-seq prediction heads from the functional-track module (GTEX-1I4MK-0002-SM-EZ6M9.1, GTEX-1LB8K-0005-SM-DIPED.1, GTEX-1OKEX-0006-SM-DKPQ2.1). For each allele-specific sequence, NTv3 produced predicted RNA-seq signal across the supervised central region, and gradients were computed with respect to the corresponding functional-track prediction. No post-hoc smoothing or normalization was applied.

For motif-level inspection, we extracted attribution windows around each SNV, retained the A/T/C/G channels and visualized nucleotide-resolved hypothetical gradients using `Logomaker` [93, 94]. Reference and alternative attribution profiles were compared directly to identify representative local responses including motif ablation, motif creation and distributed multi-motif changes. External Ensembl VEP annotations [50] were used only to report known consequence labels for selected examples such as rs9610445 in *APOL4*.

##### Multispecies Motif Discovery

We constructed independent active-enhancer cohorts for five species using species-specific regulatory catalogs rather than a shared orthology set. Human (hg38) and mouse (mm10) elements were drawn from the ENCODE v4 cCRE registries (ENCFF420VPZ/ENCSR800VNX and ENCFF167FJQ/ENCSR412JPD, respectively), retaining distal or proximal enhancer-like signatures (dELS or pELS) and excluding promoter-like signatures. Fly enhancers were obtained from the FlyBase nine-state chromatin annotations for S2 and BG3 cells (FBrf0212862), merged after lift-over to dm6 and retaining state 3 (enhancer). Worm enhancers were obtained from the RegAtlas ce11 regulatory-element catalog (GSE141213), retaining records annotated as putative enhancers; ce11 and WBcel235 denote the same assembly, and these intervals were used under the WBcel235 naming adopted by the pipeline. *Arabidopsis thaliana* enhancers were obtained from the AraENCODE combined enhancer catalog on TAIR10. Because these resources use different regulatory ontologies, enhancer status was defined within each source before harmonization.

From each catalog, we sampled 1,000 active enhancers with a fixed random seed (20260902). Reference-genome sequence was fetched for each interval, and every native enhancer was paired with an exact dinucleotide-preserving shuffle. The sequence-layer cohort contained 1,000 valid enhancer sequences per species with no sequence rejections. Attributions were computed with the single-length NTv3 model on a 1,048,576-bp window of each species’ reference genome centered on the enhancer, under that species’ token; enhancers whose window extended beyond the chromosome were skipped, leaving 989 human, 998 mouse, 963 fly, 948 worm and 948 *Arabidopsis* enhancers. The attribution target was the mean prediction of all of the species’ DNase-seq and ATAC-seq tracks over the enhancer’s output positions, and gradients were mean-centered across A, C, G and T. TF-MoDISco (modisco-lite 2.4.0) was run on a 1,000-bp crop centered on each enhancer, with a 500-bp analysis window and at most 2,000 seqlets per metacluster. Activating patterns were matched with TOMTOM-lite to one pooled JASPAR 2026 collection of vertebrate, insect, nematode and plant motifs (*p* ≤ 10^−3^). A pattern whose contribution core, from the first to the last position with at least 25% of the peak contribution, exceeded 24 bp was counted as broad. Every other pattern was assigned to the DNA-binding-domain group shared by most of its top three matches (ties to the best match); a CTCF-like assignment was kept only if the best match had *p* ≤ 10^−8^. Fig. 5j shows, for each species, the percentage of activating enhancer seqlets in each group, for groups that reach 5% in at least one species, and in broad patterns.

##### Ortholog Motif Turnover

We selected one published human–mouse enhancer ortholog from Oh and Beer [46] (hg38 `chr22:42,317,026–42,317,326`; mm10 `chr15:82,940,846–82,941,146`). Each 300-bp enhancer sequence was centered within a 1,048,576-bp window of its own genome and evaluated with its own species token, and input-times-gradient attribution was computed with respect to that species’ DNase-seq and ATAC-seq tracks. The attribution maps were cropped to a 40-bp interval spanning the AP-1 motif matches (JASPAR MA0099.2) shown in Fig. 5k.

##### Enhancer–Gene Linking Benchmark

To evaluate whether NTv3 ranks experimentally validated enhancer– gene relationships at genome scale, we scored 10,356 K562 CRISPRi pairs (471 regulated) from the ENCODE enhancer–gene perturbation encyclopedia [48], without using those linking labels during training. For each gene, a 1 Mb input window was centered on the transcription start site, a scalar objective was formed from predicted RNA-seq signal at annotated exons, and input-times-gradient attribution was computed with respect to that objective. Each candidate enhancer was then assigned a Gaussian-weighted mean of absolute attribution over the element (2,400 bp window, *σ* = 300 bp); pairs falling outside the model input window were assigned a score of zero.

Performance was summarized within the ENCODE-rE2G enhancer–TSS distance bins using auPRC, auROC and precision at 70% recall, and compared against a reciprocal-distance baseline (1/TSS) and Borzoi [5], using fold 0 of four trained folds. AlphaGenome values were taken directly from the published zero-shot K562 RNA-seq input-gradient analysis [6] (their Fig. 4j) and were not re-scored. Performance is reported on all 10,356 pairs, with unscored pairs set to zero, and on 3,826 matched pairs scored by all three models. NTv3 was evaluated using the 1 Mb checkpoint with a keep-target-center fraction of 0.375.

#### 4.8.3. LLM-Guided Interpretation Workflow

To integrate model interpretation with locus-specific genomic context, we implemented a Claude Code agent skill [95] that coordinates NTv3 and NTv3 analyses through command-line tools (Supplementary Fig. A.20). The agent translates natural-language requests into structured plans specifying the reference assembly, locus, cell type, prediction tracks and analyses, resolving missing inputs through user clarification before approval and execution. Supported analyses comprise functional-track prediction, gene annotation, gradient-based attribution and single-nucleotide variant comparison. The agent also retrieves matched experimental signal from ENCODE [96], where available, and requested genomic annotations from UCSC [97].

Model outputs and retrieved context are synthesized into a locus-level report with an interactive genome browser, linking interpretations to the underlying analysis files. A rule-based audit, enabled by default, checks that referenced evidence files exist within the run directory and that literature claims have a corresponding bibliography entry. Claims failing these checks or matching predefined causal-overstatement rules are suppressed, with decisions retained in a claim-level audit log. Each report is accompanied by analysis provenance and a script for re-executing the approved plan; an optional notebook provides access to the saved results.

### 4.9 Fine-tuning NTv3 for Conditioned Sequence Generation

We adapted NTv3 to conditional sequence generation with masked diffusion language modeling (MDLM) [34]. Each design model was initialized from the post-trained tracks NTv3 model and reused its U-Net backbone. We added task-specific condition embeddings, which entered the network through adaptive layer normalization, and an auxiliary activity-classification head used for ranking candidates. Thus, no separate generative backbone or external guidance classifier was required, although the condition embeddings and activity head introduced task-specific parameters. This framework was applied to plant promoters measured in maize protoplasts and tobacco leaves [55], tobacco enhancers measured under four conditions [56] and *Drosophila* S2-cell enhancers; only the reporter context, generated interval and conditioning variables differed among tasks.

#### 4.9.1. Model Training

To enable conditional diffusion-based sequence generation, we fine-tuned NTv3 with the MDLM objective [34]. At a high level, MDLM training is akin to standard MLM training where we add noise to a sequence in the form of masked tokens and train models to recover the masked positions. In contrast to BERT-style [35] MLM pre-training, where the proportion of masked tokens is fixed, in generative MDLM training, we use a variable amount of noise. We formally introduce MDLM training below.

Let **x** and **z** represent one-hot vectors over some vocabulary *V*, i.e., **x, z** ∈ {0, 1} ^|*V*|^ ∩ Δ^|*V*|^, where Δ^|*V*|^ is the simplex over | *V* | categories. We denote a full sequence of *L* tokens as **x**^1:*L*^, with **x**^ℓ^ as the ℓ^th^ token in the sequence. In MDLM, we use a special [MASK] token and denote its one-hot representation by **m**.

MDLM is derived from the diffusion paradigm for generative modeling [98, 99], initially introduced for continuous signals, and later adapted for discrete data [100]. These models are characterized by a pre-defined corruption process *q* that transforms signal into noise and a learned reverse process *p*_*θ*_ that is trained to recover the original signal. Specifically, the forward process adds noise to samples drawn from some data distribution **x** ∼ *q*_data_ to produce latent variables **z**_*t*_ ∼ *q*(· |**x**), where the subscript *t* ∈ [0, 1] denotes time in the diffusion process. The latents **z**_*t*_ become progressively more corrupted as *t* goes from 0 to 1, with the fully corrupted final latent variable **z**_1_ drawn from a pre-defined prior distribution. In MDLM, this prior is equal to the Dirac measure that places all of the probability mass on the special mask token one-hot **m**. That is, at the end of the forward process, all of the signal from the data sample has been removed, as all tokens have been replaced with the [MASK] symbol. Let Cat(· ; *p*) denote a categorical distribution over *V* with probabilities given by *p* ∈ Δ^|*V*|^. A widely used forward process for discrete signals is one that interpolates between data and the limiting distribution [100], and we adopt this process for MDLM as well [34, 101–103]:

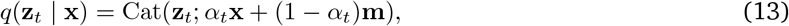

where *α*_*t*_ := *α*(*t*) ∈ [0, 1] denotes the noise schedule, which is monotonically decreasing in *t*, with *α*(1) = 0 and *α*(*t*) → 1 as *t* → 0. Here, *α*_*t*_ represents the fraction of the original signal that remains at time *t*, while 1 − *α*_*t*_ corresponds to the fraction replaced by the [MASK] token (i.e., the noise level). For a discrete-time process, we set some number of diffusion steps as *T* and let *t* = *i*/*T* for *i* ∈ {0, …, *T*} . Letting *s* < *t* denote some time earlier in the diffusion process, the marginals defined in (13) yield the following form for the true posterior distributions [34]:

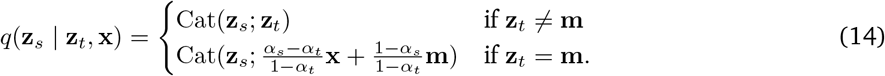

Importantly, the noising process *q* is assumed to factorize independently across the sequence length dimension, i.e., 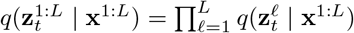 and 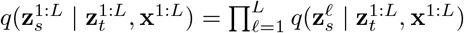.

The goal of diffusion modeling is to train a denoising process *p*_*θ*_ that can generate samples that resemble those from the original data distribution. Ideally, we could perfectly recover the exact posterior in (14); however, when generating samples, we do not have access to the true data sample **x** which is being generated. Therefore, a widely-used parameterization for *p*_*θ*_ is to define a denoising network **x**_*θ*_ ∈ Δ^|*V*|^ that takes in noisy (partially masked) inputs and returns a probability over the clean token for each position. We then plug this prediction into (14). This gives the following form for the reverse process [34]:

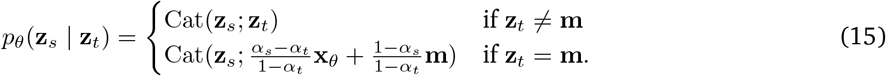

As above, we assume that this process factorizes along the sequence length: 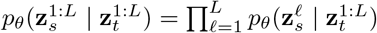. To mimic the behavior of the forward process, two additional restrictions can be placed on the denoising model: 1) we do not place any probability mass on the mask position: ⟨**x**_*θ*_, **m**⟩ = 0 and 2) for any tokens that are not noised, we simply ‘copy them over’: ⟨**x**_*θ*_, **z**_*t*_⟩ = 1, if **z**_*t*_ ≠ **m**, where ⟨·, ·⟩ denotes an inner product.

The denoising diffusion model is trained to minimize a variational upper bound on the negative log-likelihood of the training data. Taking *T*→ ∞ yields a continuous-time formulation whose variational bound simplifies to [34, 102, 103]:

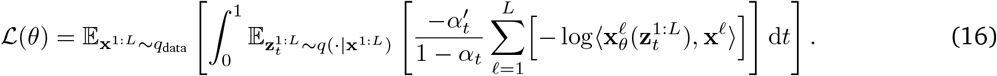

Here, 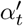 denotes the derivative of the noise schedule with respect to *t*. Because *α*_*t*_ is non-increasing, 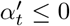, and therefore the weight 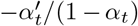 is non-negative for *t* > 0. Moreover, 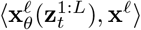 is the probability assigned to the true token, so its negative logarithm is a non-negative per-token negative log-likelihood. The integrand is therefore non-negative. Consequently, the MDLM objective is a weighted average of masked-language-modeling losses across different masking rates.

#### 4.9.2. Sequence-design Datasets and Conditions

We considered three high-throughput reporter-assay settings: plant promoters measured in maize protoplasts and tobacco leaves, tobacco enhancers measured under four conditions, and *Drosophila* enhancers measured with two core-promoters. In every setting, the full reporter construct was supplied as sequence context and only the regulatory element was corrupted and generated. Continuous activity measurements were discretized into task-specific bins and embedded as conditioning labels.

##### Plant Promoter

Promoter activity data are from a previously published plant promoter STARR-seq study [55]. The dataset comprises three species: *A. thaliana, S. bicolor, Z. mays*, each providing 170 bp promoter sequences with STARR-seq enrichment measured in two expression systems: tobacco leaf and maize protoplast. Leaf and protoplast records are outer-joined on (species, gene, sequence) so a given promoter can carry either or both measurements. The 170 bp promoter candidate is embedded into a fixed reporter backbone: pPSup (Addgene 149416), comprising a 154 bp CaMV 35S enhancer element, the 170 bp promoter slot, and a species-specific 5’ UTR from histone H3 (shared UTR for Arabidopsis and maize; a separate, longer UTR for sorghum, per Supplementary Table 8 of the 2021 paper), inserted at the geometric midpoint of the vector (splitting at the EGFP start position). The resulting insert length is 5,858 or 5,892 bp depending on species (per-species UTR length), padded to a model sequence length of 6,016 bp. Each tissue’s enrichment score is discretized into 5 bins using fixed cutpoints [-2.5, 0, 2.5, 5].

##### Plant Enhancers

Enhancer activity data were sourced from a previously published plant enhancer STARR-seq study [56]. The dataset comprises STARR-seq enrichment for 170 bp candidate accessible chromatin regions (ACRs) across four species (*Arabidopsis thaliana, Sorghum bicolor, Zea mays*, and *Solanum lycopersicum*) under four assay conditions: tobacco-leaf agroinfiltration at ambient light, dark, 35 °C (warm), and 10 °C (cold). We train on the four tobacco-leaf conditions (light/dark/warm/cold) jointly, as a single conditioning axis; the maize-protoplast channel is not used in the final model. Each raw record has a sequence ID, the 170 bp sequence, per-condition enrichment scores, and a train/val/test split label assigned by the original authors. The 170 bp candidate is embedded into a fixed reporter backbone, a modified pPSntF vector (Addgene 256230): CaMV 35S minimal promoter → maize histone H3 5’ UTR (*Zm00001d041672*) → start codon → 18 bp barcode → EGFP coding sequence, assembled via BbsI Golden Gate cloning. The vector is the same across all four species. The 170 bp insert is spliced in at the position of the Golden Gate stuffer; an 18 bp barcode region is resolved to a single fixed sequence (ATCCGTAGTGACGCAACG) before use. Backbone + insert yields a 5,670 bp construct, padded to a model sequence length of 5,760 bp. Each condition’s enrichment score is discretized into 5 bins using fixed cutpoints [−2.0, −1.0, 0.0, 1.0], chosen to work reasonably across all four conditions jointly.

##### Fly Enhancers

We used the UMI-STARR-seq data from *Drosophila* S2 cells compiled for DeepSTARR [3]. The dataset contains 484,052 genomic sequences of 249 bp with quantitative activity measurements in the DSCP and RpS12 core-promoter contexts, split 402,296 for training, 40,570 for validation and 41,186 for testing. Each candidate was inserted into the corresponding published STARR-seq plasmid [57], and the construct was cropped to 4,096 bp. Treating the two promoter contexts as separate examples yielded 968,104 sequence–activity pairs. Within each promoter context, activities were discretized into five equal-width intervals spanning the observed training-set range. We retained the training, validation and test assignments of the source study.

#### 4.9.3. Fine-tuning

For each design task, all model parameters were initialized from NTv3 and updated jointly. The condition embedding modulated adaptive layer-normalization layers. On 10% of training examples, the condition was replaced by a null label, allowing the same network to provide conditional and unconditional predictions for classifier-free guidance [53, 54]. In the fly models, an auxiliary multiclass head predicted the activity bin from the input construct; this head was used only to rank generated candidates and was not required by the diffusion sampler.

Fine-tuning used AdamW with weight decay 0.01 and an effective batch size of 32 distributed across two GPUs. The learning rate increased linearly from 5 × 10^−5^ to 10^−4^ over the first three billion nucleotide tokens and then decayed to 5 × 10^−5^ with the modified squared-decay schedule. We maintained an exponential moving average of model parameters with decay 0.9999 and used the averaged parameters for generation and evaluation.

#### 4.9.4. Conditional Sampling and Classifier-free Guidance

When generating sequences with NTv3, we used the guidance algorithm of classifier-free diffusion guidance (CFG) [53] adapted for discrete models [54].

Formally, let *y* denote some property or condition of interest. Controlled generation in diffusion models aims to steer each denoising step by sampling from the following unnormalized tempered, conditional distribution:

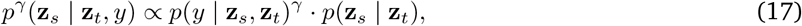

where *γ* represents a hyperparameter that controls how much we ‘focus’ on the conditioning. Applying Bayes’ rule to the first term on the right-hand side of (17) eliminates the need for the classifier *p*(*y* | **z**_*s*_, **z**_*t*_):

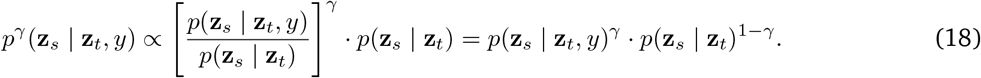

Note that in the first expression on the right of the proportionality symbol in (18), we absorbed the factor of *p*(*y* | **z**_*t*_)^*γ*^ into the proportionality constant. We can model both terms on the right-hand side of (18) using denoising diffusion models. In practice, these distributions can be generated using the same model. This is achieved by randomly dropping out or masking out the conditioning during training.

Finally, we can also introduce the notion of negative conditioning, by replacing the unconditional model in (18) with one conditioned on a background or base value *y*_bg_. In this way, we simultaneously steer the denoising distribution toward the condition of interest *y*_target_ and away from the base condition. Specifically, we sample from the following distribution:

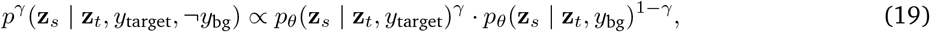

This framework allows CFG to flexibly steer generation under multiple constraints, including jointly specifying an activity level and a plasmid-specific context. In the promoter-selectivity designs, *y*_target_ and *y*_bg_ request the *same* activity bin in two different reporter contexts, so the selectivity is produced by the contexts and the guidance exponents rather than by asking for a lower activity in the background. For all experiments, we used a guidance scale *γ* = 3.0, the exponent on the implicit classifier in (17), and 50 denoising steps. This corresponds to *w* = *γ* − 1 = 2.0 in the convention of [53].

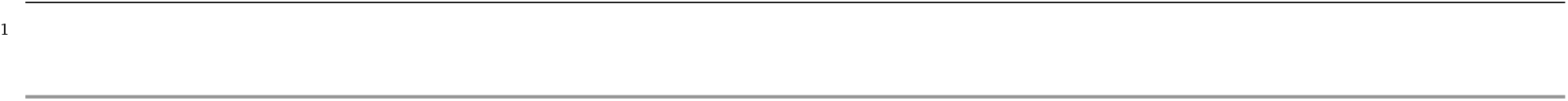
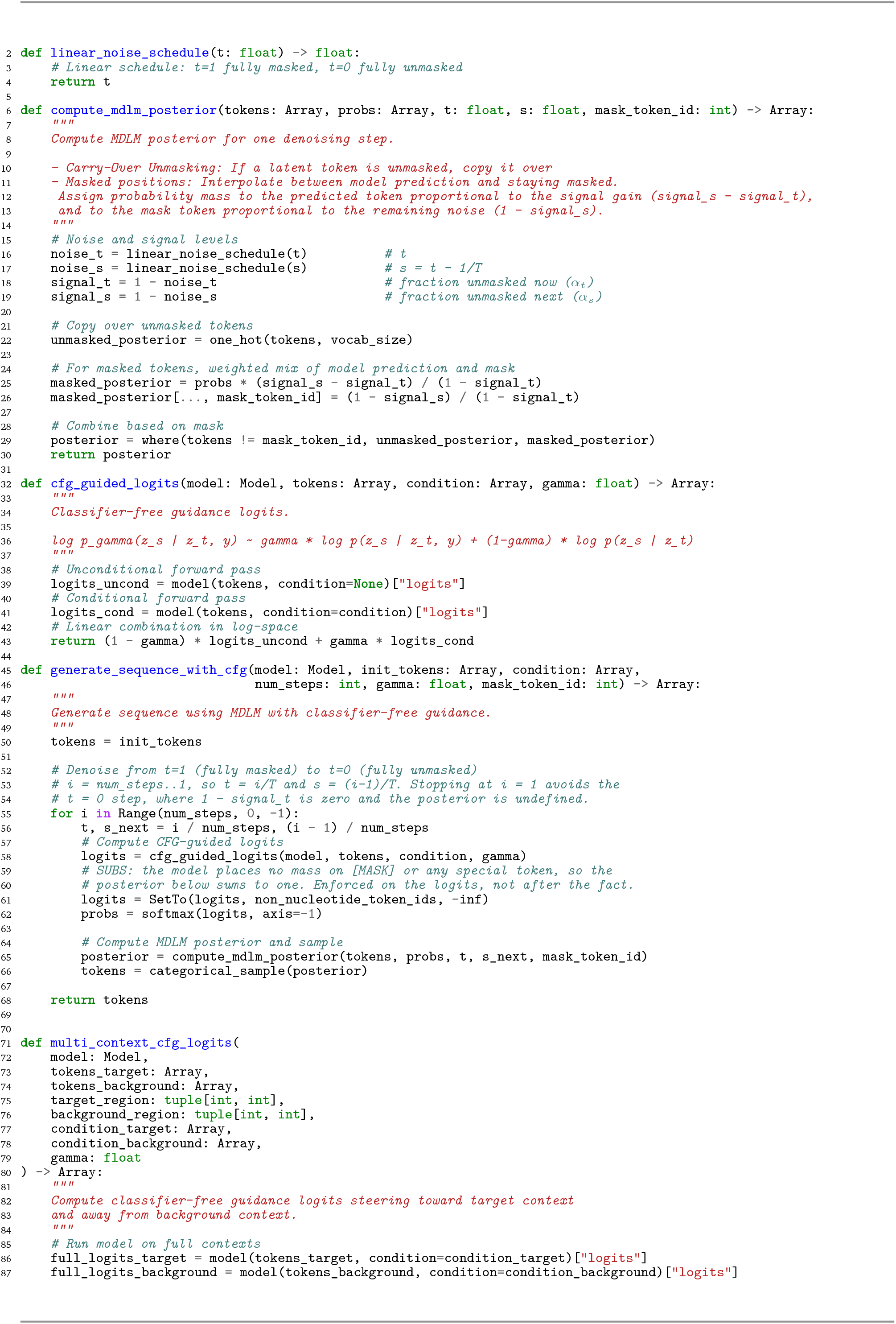
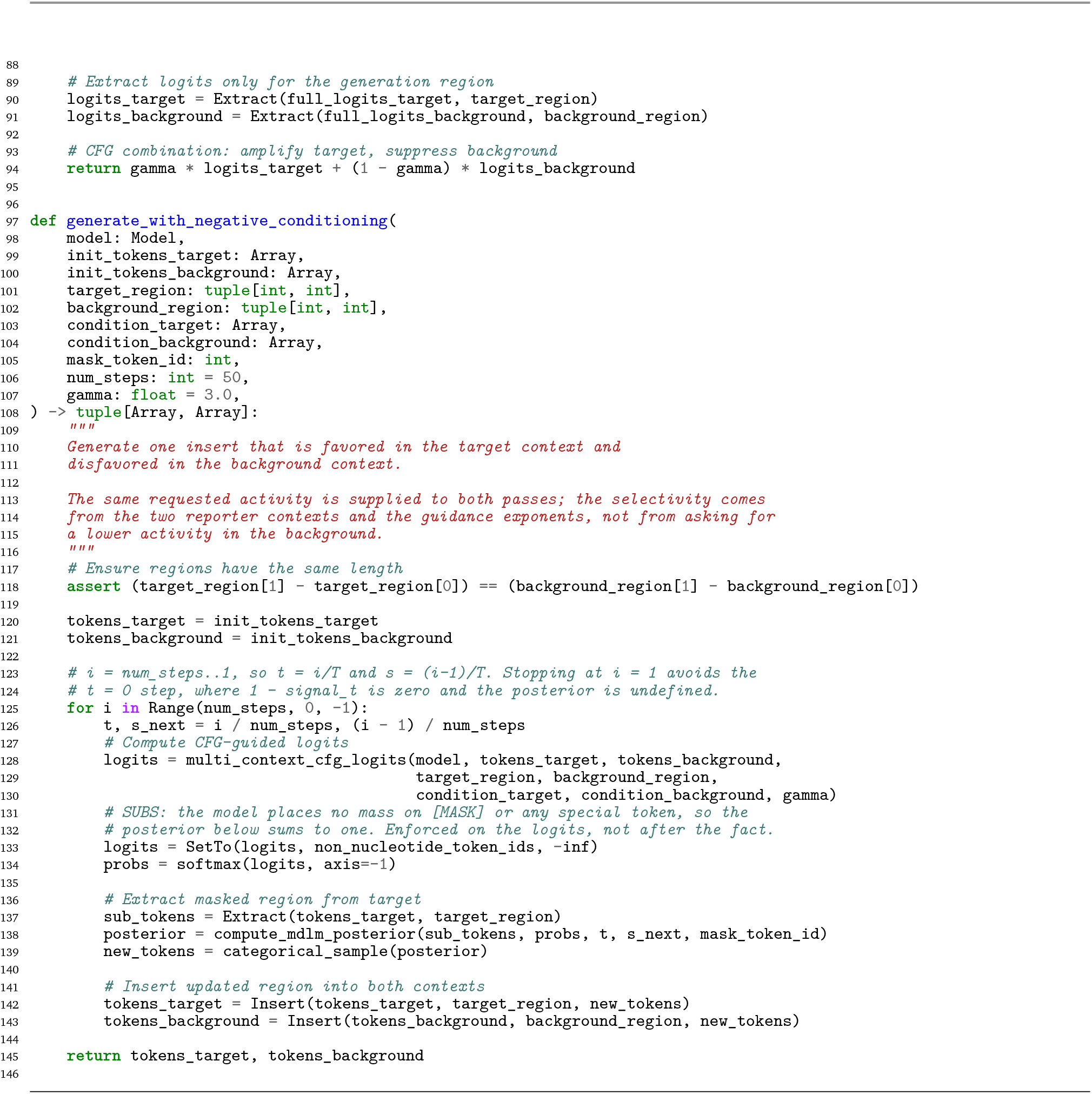

#### 4.9.5. In-silico Evaluation with Oracles

Generated sequences were evaluated with an independently trained activity predictor for each assay. Oracle outputs were compared with native held-out sequences satisfying the same measured functional condition. For graded designs, these were held-out sequences in the corresponding activity interval; for fly promoter-selective designs, they were enhancers active in only one of the two promoter assays. DeepSTARR [3] was used as the fly enhancer oracle. We additionally compared GC content and *k*-mer composition between generated, training and held-out native sequences. In addition, we ran BLAST searches against the respective genomes on all generated sequences to confirm that NTv3 proposes novel biological sequences rather than reproducing elements from the training data.

#### 4.9.6. Evaluation of Sequence Realism

Beyond predicted and measured activity, we asked how far generated sequences depart from native ones in composition. Two instruments were used: a 5-mer *L*_2_-regularized logistic regression and a small convolutional classifier, both trained to separate generated sequences from native ones and scored by area under the receiver operating characteristic curve.

The comparison is made against *activity-matched* native sequences rather than against the native pool as a whole. This matters: a generated set requested at high activity differs from a mostly middling native pool because of its activity, so an unmatched classifier can separate the two by learning the request rather than anything about sequence realism. Native sequences are therefore drawn to match the activity composition of the generated set being tested.

The convolutional classifier is the instrument reported. It applies a width-11 convolution with 64 filters, a rectified linear activation and max pooling by a factor of four, then a width-5 convolution with 64 filters and a rectified linear activation, a global maximum over positions, dropout, and two dense layers to a single logit. It is trained with Adam at a learning rate of 10^−3^ for a fixed budget of 25 epochs, with no early stopping, so that the reported value is not selected on the fold it is measured on. Reported values are out-of-fold only, averaged over four stratified folds and five random seeds.

An absolute area under the curve is not interpretable on its own here, because it depends on how many sequences the classifier is given: the same comparison on the same sequences moves by more than 0.1 across the pool sizes used in this work. Each measurement is therefore accompanied by two controls computed at the same pool size. The lower control splits the native pool into two disjoint halves and separates one from the other, which defines chance and the resolution of the test. The upper control separates the native pool from a dinucleotide-preserving shuffle of itself, which is the separability of sequence retaining native composition but not native order. Generated sequences are then reported as a position between the two, so that zero is indistinguishable from native and one is as distinguishable as a composition-matched shuffle. Dinucleotide-preserving shuffles were generated by the Altschul–Erikson algorithm [104].

We additionally report *k*-mer composition divergences. The Jensen–Shannon divergence between pooled 4-mer frequency profiles, in bits, is reported for each generated set against its own genome’s native profile, with the dinucleotide shuffle of that genome as a reference level. We also report the 6-mer Jensen–Shannon *distance*, the square root of the divergence, which is the statistic published generative baselines report. It is given for all eight design contexts, not only for the comparison against those baselines.

#### 4.9.7. Benchmarking with D3 on Fly Enhancers

D3 [58] applies score-entropy discrete diffusion to regulatory sequence design and releases its checkpoints, its activity oracle, its preprocessed data and its evaluation protocol on the same *Drosophila* enhancer dataset used here, which makes it the closest available point of comparison for NTv3’s generative model. We compared against its released transformer checkpoint.

The two methods condition differently, and that difference has to be controlled rather than ignored. D3 conditions each sample on a held-out record’s own continuous activity, so its designs reproduce the native activity distribution by construction. NTv3 is prompted with a discrete activity bin and was sampled uniformly across bins, which places a much larger share of its designs at the extremes than the native distribution contains. A classifier given those two pools can separate them by learning the requested activity rather than anything about the sequences. We therefore report NTv3 twice: as generated, and redrawn without replacement from the same design pool so that its activity composition matches the native one. The matched arm is the one used for comparison; the uniform arm is reported so the size of the effect stays visible.

Both methods are scored through D3’s own evaluator, using the activity oracle distributed with D3 rather than the original DeepSTARR checkpoint. Their oracle is a DeepSTARR model trained with EvoAug augmentation [3, 105], and scoring with the stock checkpoint instead would shift the oracle-dependent metrics enough to resemble a failed reproduction. Every arm is scored with that same oracle, so no asymmetry is introduced.

##### Evaluation procedure

Two metrics are reported, both of which depend only on sequence and not on the oracle. The first is the area under the ROC curve of a classifier trained to separate a generated pool from a native reference pool, so that 0.5 means the two are indistinguishable. The second is the 6-mer Jensen–Shannon distance between the pooled *k*-mer compositions of the two sets. Both are reported as a position within a bracket whose lower end is two disjoint halves of the native test set and whose upper end is a dinucleotide-preserving shuffle of it, so that zero is native-like and one is as distinguishable as a composition-matched shuffle. Reference and generated sets are drawn from opposite halves of a permutation of D3’s test split, so each comparison uses disjoint sequences, and each measurement is the mean over five such draws. Two further metrics from their protocol are not reported: a paired fidelity error, which requires one design per held-out record conditioned on that record’s own continuous activity and therefore has no counterpart for bin-prompted designs, and a marginal distribution statistic, which scores the conditioning scheme rather than the sequences.

### 4.10 Experimental Validation of Designed Fly Enhancers

#### 4.10.1. Selection of Candidate Generated Sequences

The activity prediction head of NTv3 was used to filter all generated sequences prior to *in cellulo* evaluation. For experiments aimed at designing enhancers with specified activity levels, we selected the 60 sequences with the highest predicted probability for each target activity bin from 1,024 generated candidates per bin, yielding a total of 600 sequences across five bins and two promoter contexts. For experiments targeting promoter-specific enhancers, we selected 150 sequences that displayed the largest predicted activity difference between the two promoters from 10,240 generated sequences, 75 for each promoter context. A further 100 sequences were generated under a condition requesting high activity in both promoter contexts simultaneously. We note that this selection of generated sequences was done by the first version of NTv3 post-trained model, but we have confirmed that the generated sequences retain similar scores with our newly revised NTv3 model presented in this manuscript version. As a baseline for sequence design, we sampled a pool of random sequences uniformly over A, T, C and G and used the DeepSTARR [3] oracle to identify the 150 with the largest predicted activity difference between the two promoter contexts, 75 per context, representing oracle-guided selection without a generative model.

#### 4.10.2. Generating STARR-seq Libraries

300-bp oligonucleotides containing 249-bp candidate sequences were custom synthesized by Twist Bioscience and amplified as inserts for oligonucleotide STARR-seq plasmid libraries in two consecutive PCR steps. The first PCR was performed in the linear range with the primer pair (forward: ACACTCTTTCCCTACACGACGCTCTTC-CGATCT, reverse: GTGACTGGAGTTCAGACGTGTGCTCTTCCGATCT). In the second PCR amplification step, overhangs were added for Gibson cloning using the following primer pair (forward: TAGAGCATGCACCGGA-CACTCTTTCCCTACACGACGCTCTTCCGATCT, reverse: GGCCGAATTCGTCGAGTGACTGGAGTTCAGACGTGT-GCTCTTCCGATCT). Amplified fragments were cloned using Gibson assembly (New England BioLabs, catalog no. E2611S) into the previously published STARR-seq plasmid containing the DSCP or RpS12 promoter (Addgene 71499[57] and Addgene 71504[59]). Oligonucleotide libraries were electroporated into MegaX DH10B electrocompetent bacteria (Thermo Fisher C640003) and grown in 2L LB-Amp (Luria-Bertani medium plus ampicillin, 100 µg/mL) each and purified with a Qiagen Plasmid Plus Mega Kit (catalog no. 12981). All libraries were then tagged with a unique molecular identifier (UMI), amplified and sequenced to derive an input count value for the representation of sequences in the plasmid library, as previously described[106].

#### 4.10.3. Cell Culture

*Drosophila* S2 cells (Thermo Fisher R69007) were grown in Schneider’s Insect medium (Gibco 21720001) with 10% fetal bovine serum (FBS) (Sigma-Aldrich, heat inactivated) and 1% P/S (Gibco 15140122) at 27°C and 0.4% CO_2_. For electroporations, the regular growth of cells was monitored, and cells were seeded at a density of 3 × 10^6^/mL the day before electroporation. For each biological replicate, 50 × 10^6^ cells were resuspended in 100 µL of a 1:1 dilution of HyClone MaxCyte electroporation buffer and serum-free Schneider’s and electroporated with 5 µg of the input libraries (see previous section) using the MaxCyte-STX system (“Optimization 1” protocol). Cells were then incubated in DNase I (Worthington Biochemical, 2000 U/mL) for 30 min and harvested for RNA extraction 24 h after addition of culture media for recovery and oligonucleotide library expression. RNA was extracted and processed following the previously described UMI-STARR-seq protocol [106].

#### 4.10.4. Illumina Sequencing

Next-generation sequencing was performed at the VBCF NGS facility on a NovaSeq SP platform (150 bp paired-end), following the manufacturer’s protocol, using standard Illumina i5 indexes as well as UMIs at the i7 index.

#### 4.10.5. STARR-seq Data Analysis

A custom reference containing all assayed candidate sequences was constructed using `buildindex` from Rsubread version 2.10.0 [107]. Paired-end reads from the plasmid-input and reporter-RNA libraries were aligned to this reference using `align` with `type = “dna”`, `unique = TRUE` and `maxMismatches = 2`. Only read pairs assigned uniquely to a candidate sequence were retained. UMIs were extracted from the i7 read and collapsed as described previously [3], such that reads assigned to the same candidate and carrying the same UMI contributed one molecule to the corresponding count. This produced a candidate-by-sample matrix of UMI-collapsed plasmid-input and reporter-RNA counts for each core-promoter context.

##### Activity quantification and quality control

Enhancer activity was quantified independently for the DSCP and RpS12 reporter contexts using DESeq2 [108]. Candidates were retained when they were represented by at least 10 UMI-collapsed molecules in every corresponding plasmid-input replicate.

We evaluated library quality from the number of uniquely aligned, UMI-collapsed molecules, the fraction of synthesized candidates detected in the plasmid input and the agreement between biological replicates. Replicate agreement was assessed by Pearson correlation of log-transformed UMI counts separately for plasmid-input and reporter-RNA libraries.

##### Validation of activity-conditioned designs

For graded enhancer design, 60 generated sequences were evaluated for each of five requested activity bins in each promoter context. The measured activity distributions were summarized separately by requested bin and promoter context. Native genomic enhancers were assigned to activity bins using their DeepSTARR predictions. The monotonic relationship between the requested activity bin and experimentally measured enhancer activity was quantified using Spearman’s rank correlation, calculated across individual sequences separately for generated and native enhancers and for each promoter context. Requested bins were encoded in increasing order from the lowest to the highest activity condition. To assess oracle calibration, we also computed, for each bin and promoter context, the difference between median oracle-predicted and median experimentally measured activity.

##### Promoter-specificity analysis

Promoter specificity was evaluated only for sequences with valid activity measurements in both reporter contexts. For a sequence designed to target promoter *p* rather than the alternative promoter 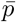 promoter specificity was defined as

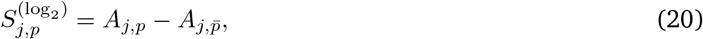

and converted to target-to-alternative fold change as

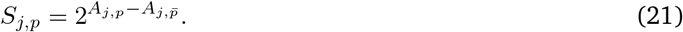

Thus, values greater than one indicate preferential activity with the requested target promoter. Specificity was calculated identically for NTv3-generated sequences and for sequences selected from the random candidate pool using DeepSTARR. The DSCP-targeted and RpS12-targeted groups each contained 75 selected sequences per design strategy before count-based quality control. For each group, we averaged log_2_ specificity and calculated its 95% confidence interval in log_2_ space. The mean and interval were then exponentiated, so bars in Fig. 6h show geometric mean target-to-alternative fold changes and error bars show 95% confidence intervals.

## Data availability

All data used for NTv3 pre-training, post-training, and downstream fine-tuning were obtained from publicly available resources. The OpenGenome2 pre-training corpus is available via Hugging Face at https://huggingface.co/datasets/arcinstitute/opengenome2. Functional genomics tracks were obtained from the ENCODE consortium (https://www.encodeproject.org/) and from NCBI repositories (e.g., SRA/GEO; https://www.ncbi.nlm.nih.gov/) and are detailed in Supplementary Table C.1. Functional genomic tracks for domesticated animals were restricted to datasets contributed through the Functional Annotation of Animal Genomes (FAANG) Consortium (https://data.faang.org/). For *C. elegans*, some functional genomic tracks were obtained from a Waterston laboratory modENCODE project (http://data.modencode.org/). NTv3 Benchmark data is described in Supplementary Table C.4 and available at https://huggingface.co/datasets/InstaDeepAI/NTv3_benchmark_dataset_revised. Gene annotations were obtained from GENCODE (https://www.gencodegenes.org/) and Ensembl (https://www.ensembl.org/). For NTv3, genome assemblies and reference gene annotations for mammals, plants and insects were obtained from NCBI RefSeq (https://www.ncbi.nlm.nih.gov/refseq/), Ensembl, JGI Phytozome (https://phytozome-next.jgi.doe.gov/) and TAIR (Araport11; https://www.arabidopsis.org/); the accession and annotation source of every species are listed in Supplementary Table C.2. The clade-specific train/validation/test genome splits and the BRAKER annotations of the insect training and validation genomes follow the Tiberius studies [28, 29], and the multi-isoform annotations of the three Phytozome-annotated test plants were taken from the Tiberius training data release [86]. NTv3 gene annotation predictions for all test genomes are available in the corresponding model repositories on Hugging Face (see Code availability). Additional dataset-specific accession identifiers and processing details are provided in the Methods and Supplementary Notes and Tables.

## Code availability

We make the code, model checkpoints, and interactive resources publicly available for research use as described below.

### Code repositories

The JAX implementation of NTv3, including model definitions and inference code, is available on GitHub at https://github.com/instadeepai/nucleotide-transformer. The same repository provides the Claude Code agent skill and accompanying tools for LLM-guided interpretation. PyTorch implementations supporting model loading, inference, fine-tuning, and downstream applications are distributed via Hugging Face and available from the *NTv3 Space* at https://huggingface.co/spaces/InstaDeepAI/ntv3.

### Model checkpoints

All NTv3 model checkpoints are released on Hugging Face via the NTv3 collection:

- NTv3 pre-trained: https://huggingface.co/InstaDeepAI/NTv3_pretrained
- NTv3 tracks, mixed-length (inputs from 16 kb to 1 Mb; recommended): https://huggingface.co/InstaDeepAI/NTv3_tracks
- NTv3 tracks, single-length 1 Mb (used for the results in this paper; use 1-Mb inputs): https://huggingface.co/InstaDeepAI/NTv3_tracks_1mb
- NTv3 annotation on mammals: https://huggingface.co/InstaDeepAI/NTv3_annot_mammals
- NTv3 annotation on plants: https://huggingface.co/InstaDeepAI/NTv3_annot_plants
- NTv3 annotation on insects: https://huggingface.co/InstaDeepAI/NTv3_annot_insects

### NTv3 Benchmark

A benchmark leaderboard reporting model performance across tasks and species is available at https://huggingface.co/spaces/InstaDeepAI/ntv3_benchmark.

### Interactive notebooks

We provide executable end-to-end tutorial notebooks for several workflows. They are provided in the *NTv3 Space*, under https://huggingface.co/spaces/InstaDeepAI/ntv3/tree/main/notebooks_tutorials:

- *Quickstart inference*: load NTv3 and run a forward pass to obtain per-base logits and sequence embeddings.
- *Tracks prediction*: predict functional tracks for a genomic region and plot selected assays.
- *Fine-tuning NTv3 on bigWig tracks*: fine-tuning on user-supplied bigWig tracks for a new assay or species.
- *Fine-tuning NTv3 into a gene-annotation model*: train on the genomes of a new set of species, select a checkpoint on held-out validation species and score its gene predictions on held-out test species.
  - *From per-base predictions to gene models*: decode the per-base state predictions into GFF3 gene models with the constrained Viterbi grammar of Supplementary Fig. A.14.
  - *Evaluating a gene annotation against a reference*: score a GFF3 file with the benchmark protocol of this study, reproducing the reported results for the released models or applying it to a user-supplied annotation.
- *Model interpretation*: compute gradient-based attribution maps and inspect transformer attention.
- *Fine-tuning NTv3 into a diffusion model*: masked-diffusion post-training of the backbone for conditional sequence design.
- *Generating plant promoter sequences*: sample plant core promoters conditioned on a requested activity level and assay system.

## Acknowledgments

This research was supported by Cloud TPUs from Google’s TPU Research Cloud (TRC). We also extend our gratitude to the researchers who deposited experimental data in public databases, those who maintain these databases, and those who make analytical and predictive methods accessible to the scientific community. F.K.L. is supported by an ESPRIT fellowship of the Austrian Science Fund (FWF; 10.55776/ESP9305624). The work of Y.S. and V.K. was supported by the National Science Foundation under award CAREER 2145577, and by the National Institutes of Health under award MIRA R35GM151243. Research at the Institute of Molecular Pathology (IMP) is supported by Boehringer Ingelheim GmbH. Next-generation sequencing (NGS) was performed by the Next Generation Sequencing Facility at Vienna BioCenter Core Facilities (VBCF), a member of the Vienna BioCenter (VBC), Austria.

## Author contributions

B.P.d.A. and T.P. conceived the research idea. S.B., B.E., Z.T., Y.A., Y.B., S.S., A.P., C.R., E.S., D.E., J.M.-R., A.H., F.A.-A., T.K., Y.S. and B.P.d.A. performed the computational analyses. F.K.L. performed the experimental work with advice from A.S. E.S. served as the scientific illustrator and led the figure synthesis. C.R. led revised figure assembly and data-to-figure integration. M.L., A.L., K.B., P.K., V.K. and A.S. provided advice on study design and analyses. B.P.d.A. and T.P. wrote the paper with contributions from all co-authors.

## Supplementary Material

### A. Supplementary Figures

**Supplementary Figure A.1.**
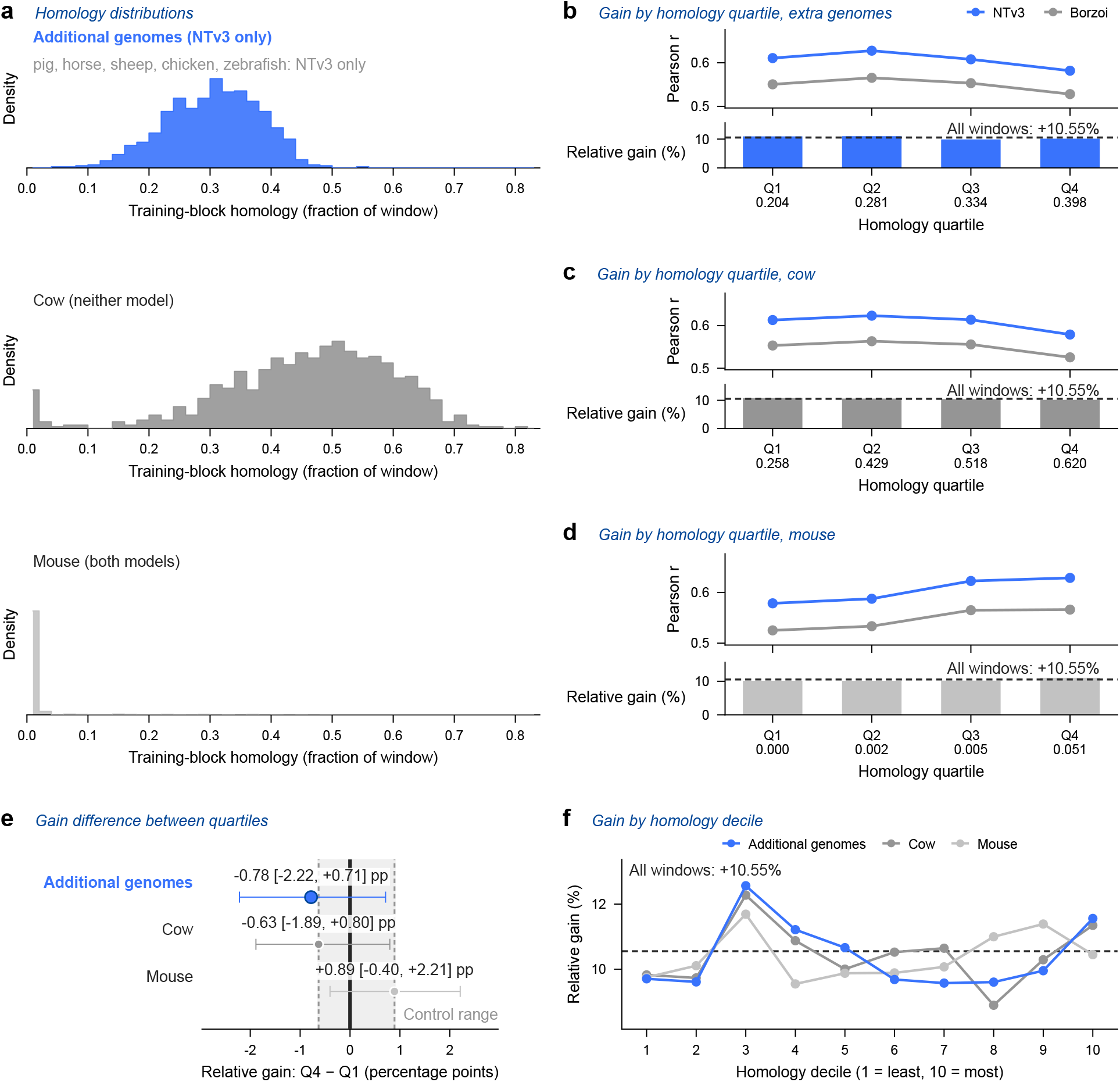
No detectable association between cross-species sequence homology and the performance margin. Association between homology to the additional training genomes and the performance difference between NTv3 and Borzoi on held-out human sequences. Analyses use a matched 312,213,504-bp footprint (mean Pearson *r*, 0.6092 and 0.5511, respectively; relative difference, +10.55% across 7,362 tracks). **a** Distributions of training-block homology to the five additional genomes (pig, horse, sheep, chicken and zebrafish), cow and mouse. Cow is absent from both training sets, whereas mouse is present in both. **b–d** Relative performance difference by homology quartile for the five additional genomes (**b**), cow (**c**) and mouse (**d**). The upper strip of each panel shows the mean per-track Pearson *r* of each model and the lower strip their relative difference, with the dashed line marking the +10.55% difference across all windows; the value beneath each quartile is its mean training-block homology. **e** Difference between the highest- and lowest-homology quartiles; intervals are 95% bootstrap confidence intervals. The shaded control range spans the two control point estimates, cow (−0.63) and mouse (+0.89); neither control can carry leakage, and the additional-genome estimate (− 0.78) is no higher than either. **f** Corresponding analysis using homology deciles. All human windows have non-zero homology to at least one additional genome, so this analysis tests a homology gradient rather than a zero-homology holdout. For the additional-genome analysis, the upper 95% confidence limit corresponds to 7% of the overall performance margin.

**Supplementary Figure A.2.**
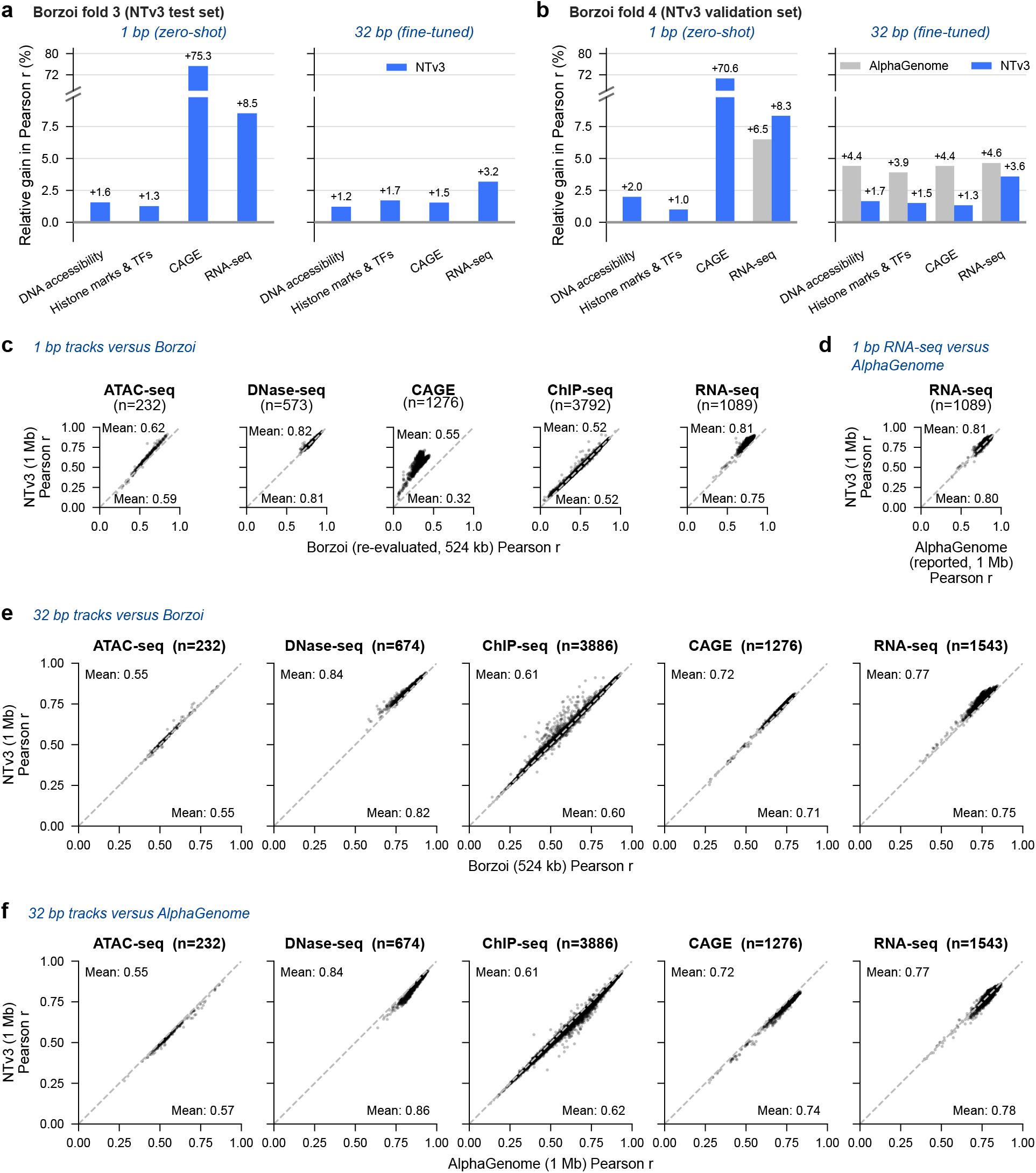
Fold- and track-level comparisons with Borzoi and AlphaGenome. **a**,**b** Relative difference in mean Pearson *r* between NTv3 and Borzoi for each assay group, evaluated zero-shot at 1-bp resolution and after fine-tuning at 32-bp resolution on Borzoi folds 3 (**a**) and 4 (**b**). In **b**, gray bars give the corresponding relative difference between AlphaGenome and the same Borzoi baseline. Fold 3 lies in the NTv3 test split; fold 4 lies in its validation split and is the fold for which published AlphaGenome results are available. The vertical axis is interrupted to accommodate the 1-bp CAGE result. **c–f** Per-track comparisons on fold 4 against Borzoi at 1 bp (**c**), AlphaGenome RNA-seq at 1 bp (**d**), Borzoi at 32 bp (**e**) and AlphaGenome at 32 bp (**f**). AlphaGenome reported 1-bp results only for RNA-seq. The 32-bp panels use all 7,611 published hg38 Borzoi tracks, whereas the 1-bp panels use the 7,362 tracks of the NTv3 post-training corpus. The 1-bp comparison uses our re-evaluation of Borzoi replicate 0, since published Borzoi values are 32-bp only. For the 1-bp comparison, Borzoi predictions were expanded from 32-bp bins across their constituent bases; the CAGE comparison therefore reflects output resolution as well as model performance. The 32-bp NTv3 values use the checkpoint obtained after 310 billion fine-tuning tokens.

**Supplementary Figure A.3.**
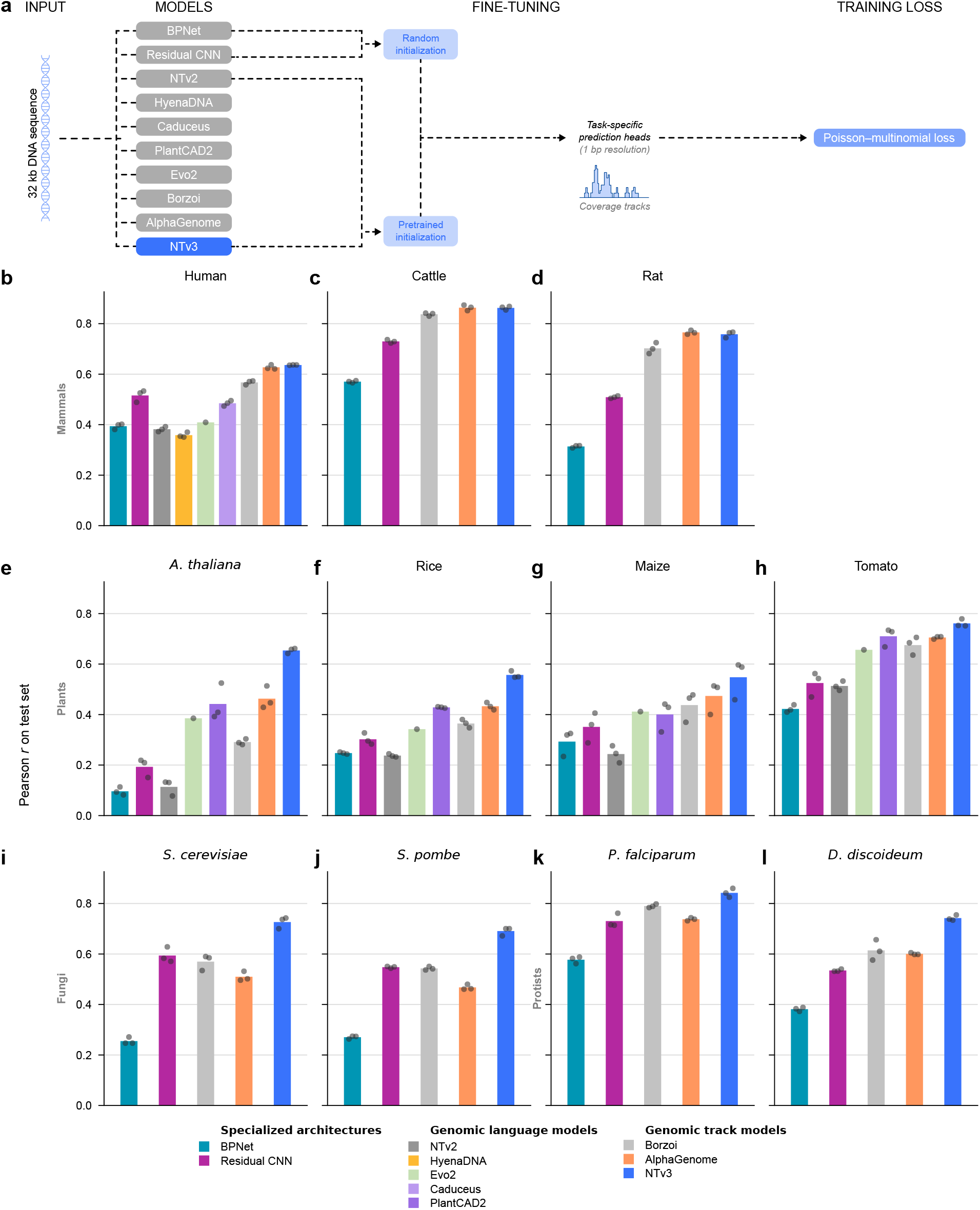
Per-species performance in the NTv3 Benchmark. **a** The standardized evaluation protocol. Each model receives the same 32 kb DNA sequence and predicts task-specific coverage at 1-bp resolution using the same prediction head and per-nucleotide Poisson–multinomial loss. BPNet and the residual CNN were trained from random initialization; the remaining models were initialized from pre-trained checkpoints and fine-tuned (Methods 4.6). **b–l** Mean Pearson *r* on held-out test tracks for 11 species, grouped as mammals (**b–d**), plants (**e–h**), fungi (**i**,**j**) and protists (**k**,**l**). Bars show means across independent fine-tuning seeds and points show individual seeds (*n* = 3 per model and species, except Evo2, for which *n* = 1). A missing bar indicates that the corresponding model was not evaluated for that species.

**Supplementary Figure A.4.**
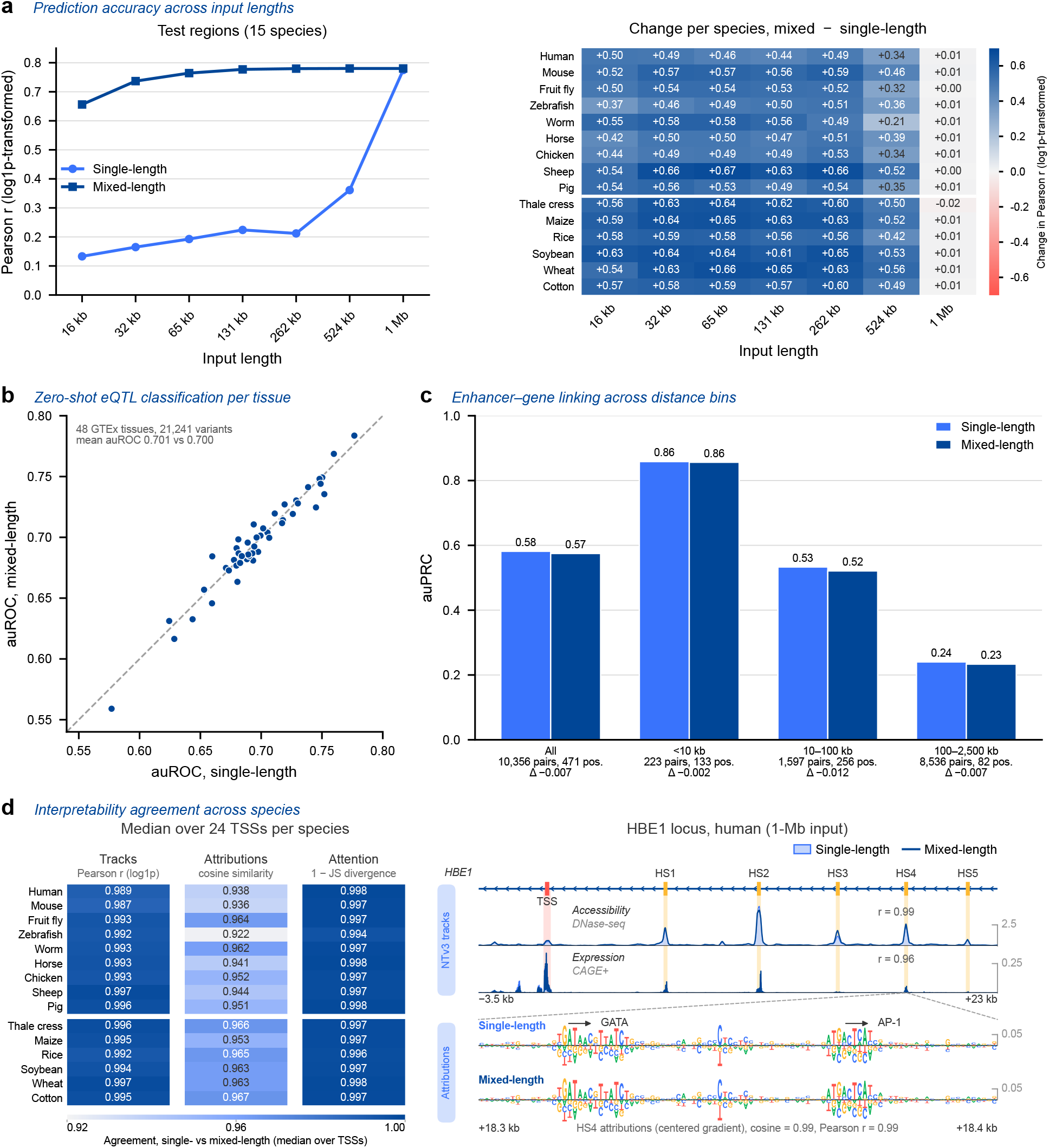
Comparison of the mixed-length and single-length NTv3 post-trained models. **a** Pearson correlation between predicted and observed log1p-transformed tracks (1-bp resolution) on test regions of the 15 post-training species, against input length. Every length is scored on the same central 393 kb of each region. Left, mean across species; right, difference between the mixed-length and single-length models for each species and input length (animals above the white line, plants below). **b** Zero-shot eQTL auROC per GTEx tissue (48 tissues, 21,241 variants, 1-Mb inputs); the dashed line marks equality, and the inset gives the mean auROC weighted by variants per tissue (single-length vs mixed-length). **c** Enhancer–gene auPRC on 10,356 K562 CRISPRi pairs by enhancer-to-TSS distance; below each bin, the number of pairs and positives and the auPRC difference (mixed-length minus single-length). **d** Agreement between the single-length and mixed-length models across the 15 post-training species (24 expressed genes per species; 1-Mb inputs; Methods 4.3.9). Left, medians over genes of track agreement (Pearson *r* of log1p-transformed predictions over the central 393 kb, median across tracks), attribution agreement (cosine similarity of input-times-gradient maps over the central 49 kb) and attention agreement (one minus the Jensen–Shannon divergence of last-layer attention from the TSS bin), on one color scale (0.92–1.00). Right, the *HBE1* locus (Fig. 5b): DNase and CAGE+ predictions of the single-length (filled) and mixed-length (line) models (*r*, Pearson correlation of log1p predictions over the window), and each model’s HS4 attributions (GATA and AP-1 motifs marked; cosine similarity and Pearson *r* over the 100 bp shown).

**Supplementary Figure A.5.**
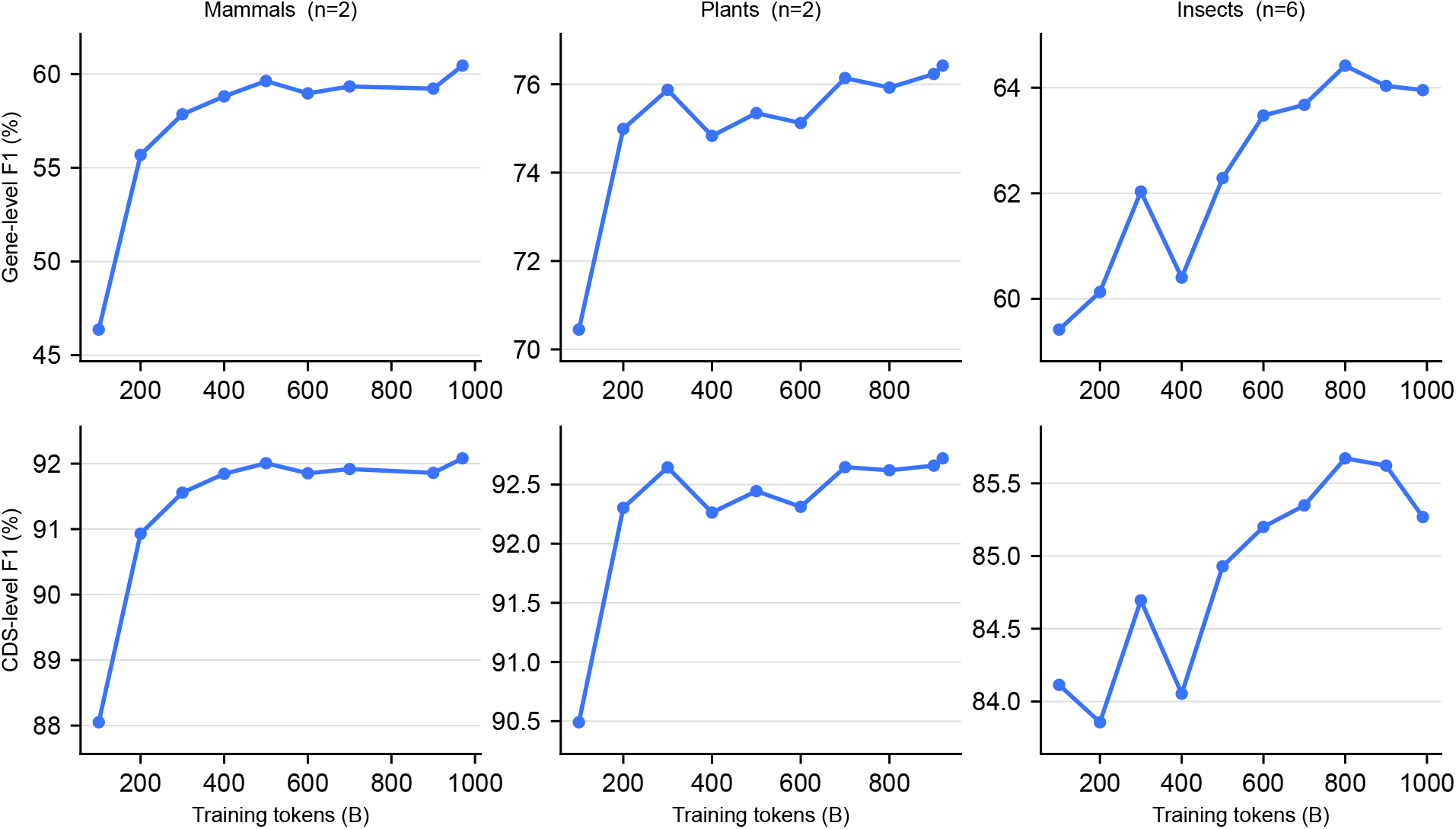
NTv3 clade-specific annotation model validation curves. Mean F1 on the held-out validation species of each clade (*n* in each title), at gene level (top) and CDS-exon level (bottom), over annotation fine-tuning; one checkpoint per 100 billion tokens plus each run’s final checkpoint. Every run has the same 1T-token budget. The reported model is the highest-ranking checkpoint on these species, reached at step 925,089 for mammals (970 B tokens seen), 877,404 for plants (920 B) and 762,960 for insects (800 B). Test species contribute to no part of that choice.

**Supplementary Figure A.6.**
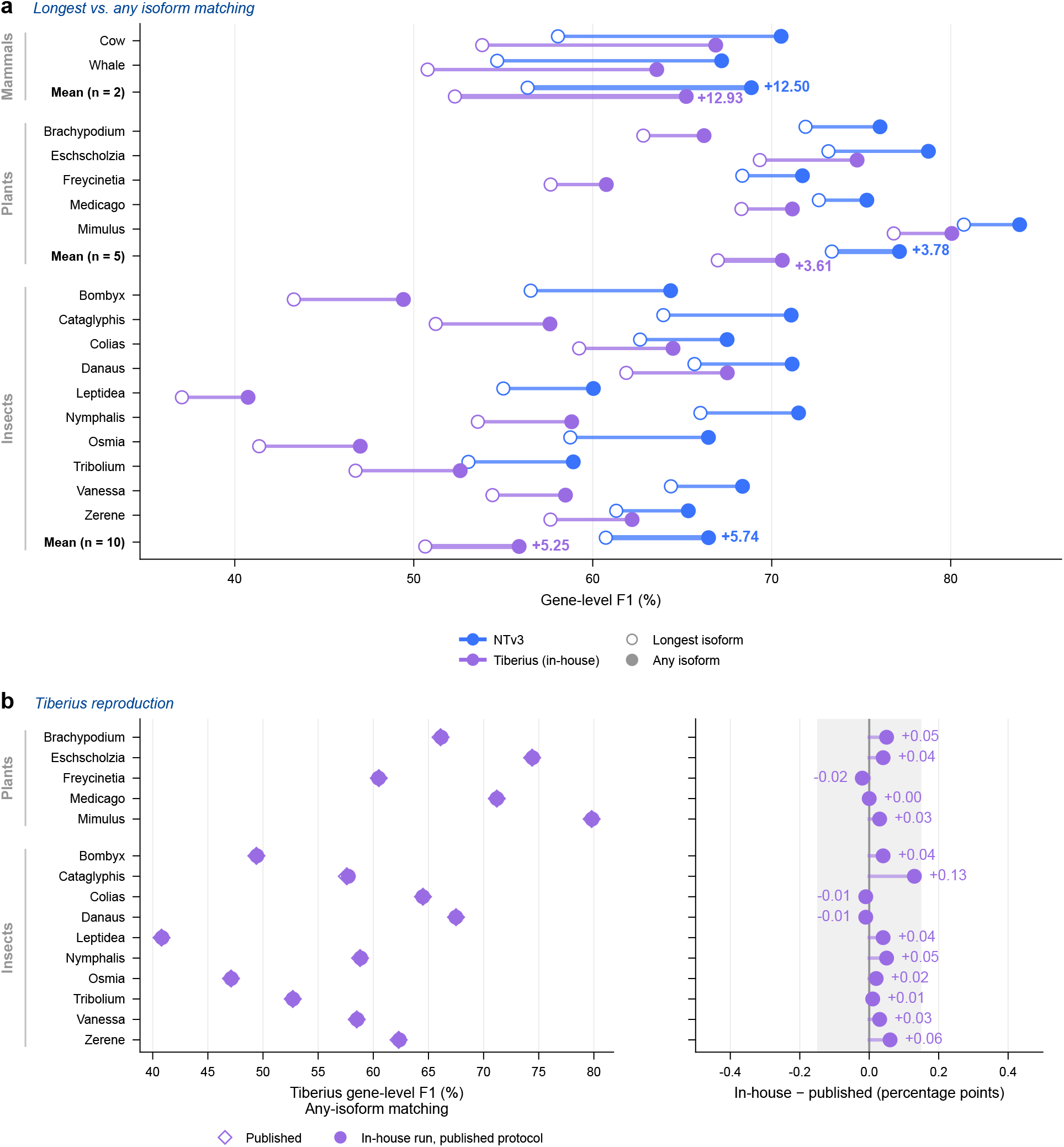
Isoform matching, and reproduction of published Tiberius accuracy. Held-out species of the three clades; *D. melanogaster* is excluded (see Methods 4.4.2). **a** Gene-level F1 of NTv3 and of our in-house Tiberius runs under our evaluation, when a prediction must reproduce the longest annotated isoform (open symbols; our primary rule) or any annotated isoform of the gene (filled symbols). Bold rows are clade means, annotated with the mean gain. The looser rule lifts both callers alike, by 12.5–12.9 points in mammals, 3.6–3.8 in plants and 5.3–5.7 in insects, and preserves their margin in every clade: many predictions counted as errors under the longest-isoform rule exactly reproduce another annotated transcript of the gene. The gain grows with the share of loci carrying alternative isoforms (13.2–45.4% across species, about 45% in mammals). **b** The Tiberius clade preprint reports any-isoform accuracy computed with gffcompare [29]. Scored with that protocol (`--strict-match -e 3`, full multi-isoform annotation, coding sequence only), our in-house Tiberius runs reproduce their published gene-level F1 for the five plants and ten insects to within 0.13 points (left, values; right, in-house minus published, shading *±*0.15 points). Mammals are omitted: their published values come from the `mammalia_nosofttmasking_v2` model (name as distributed), our runs from the soft-masking one.

**Supplementary Figure A.7.**
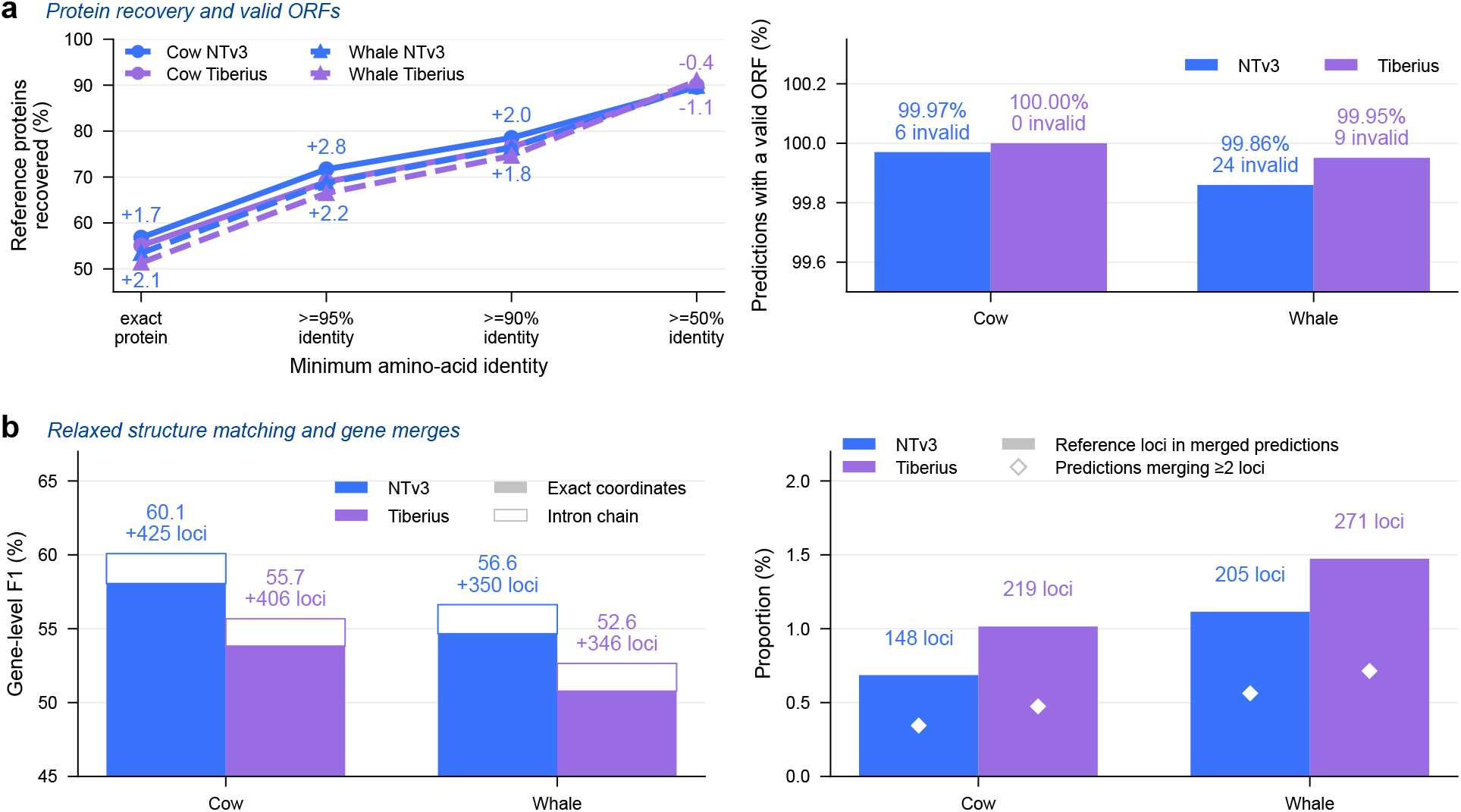
Protein recovery and relaxed gene-structure matching in held-out mammals. **a** Left, proportion of reference proteins recovered at the indicated minimum amino-acid sequence identity. Right, proportion of predicted proteins with a valid open reading frame, defined as beginning with methionine and containing no internal stop codon; numbers denote invalid predictions. **b** Left, gene-level F1 under exact-coordinate matching (filled bars) and intron-chain matching (outlined bars). Intron-chain matching requires all internal intron boundaries to match but ignores the outermost transcript coordinates; annotations give the number of additional loci recovered. Right, proportion of reference loci incorporated into merged predictions (bars) and proportion of predicted gene models responsible for these merges (diamonds).

**Supplementary Figure A.8.**
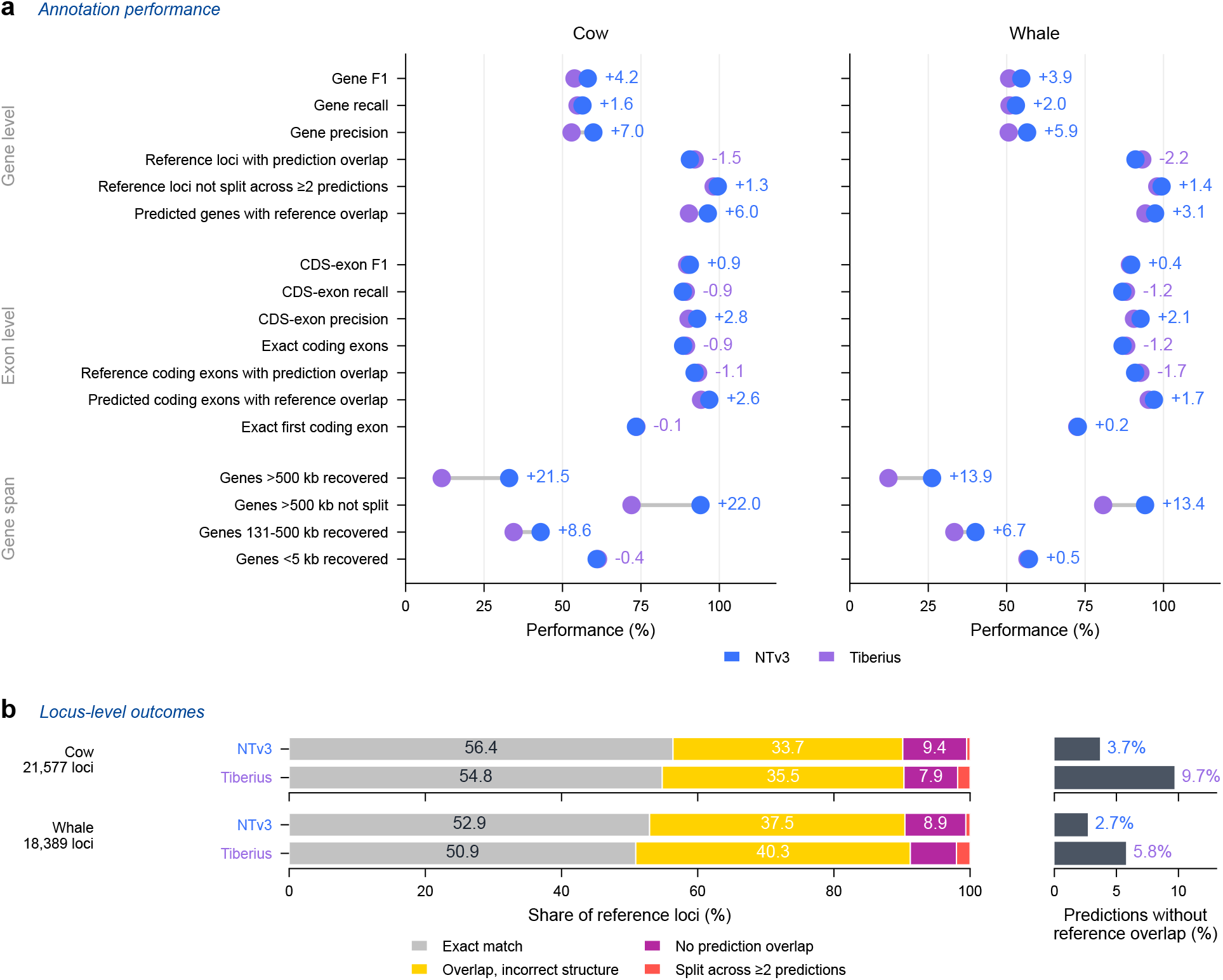
Absolute annotation performance and locus-level outcomes in held-out mammals. **a** Seventeen metrics for NTv3 and Tiberius in cow and beluga whale: the thirteen gene- and exon-level measures used throughout (Methods 4.4.3), plus four gene-span-level measures reported only here. Metrics are oriented so that higher values indicate better performance; labels give the signed difference, NTv3 minus Tiberius, in percentage points. **b** Partition of reference loci into four mutually exclusive outcomes: recovered exactly, overlapped by a prediction with an incorrect structure, not overlapped by any prediction, or split across two or more predictions. The right-hand panels show predicted gene models with no overlapping reference locus as a proportion of all predictions. The evaluation includes 21,577 cow loci and 18,389 beluga whale loci.

**Supplementary Figure A.9.**
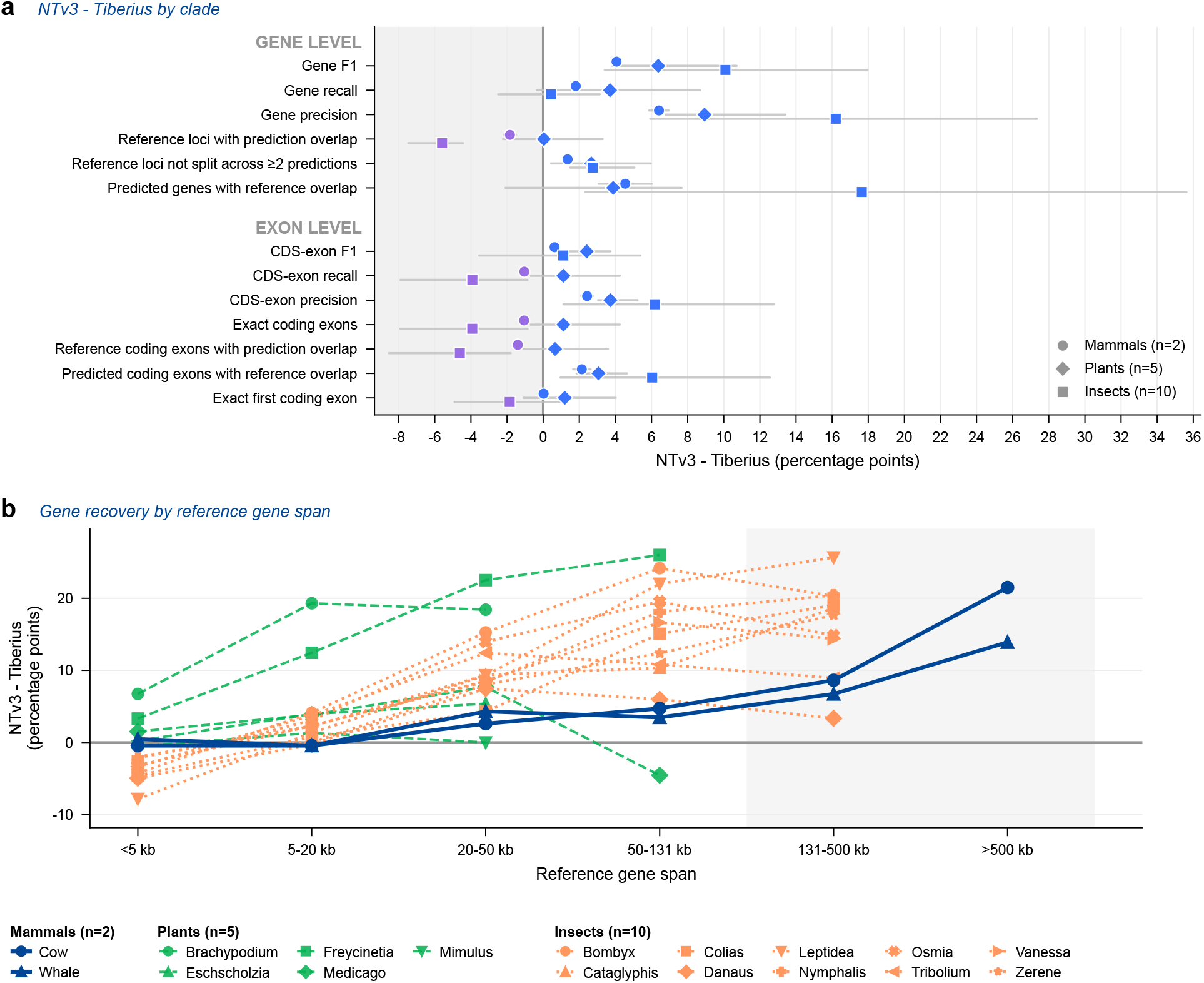
Clade- and gene-length dependence of annotation performance. Comparison of NTv3 with Tiberius across mammals, plants and insects. **a** Signed performance difference in percentage points for each annotation metric; positive values favor NTv3. Symbols show clade means across two mammals (circles), five plants (diamonds) and ten insects (squares), and gray intervals show the range across species within each clade. **b** Difference in the proportion of reference genes recovered by NTv3 and Tiberius as a function of locus span. Lines denote species and colors denote clades. Bins are shown only when both callers were evaluated on at least 20 reference genes. Shading marks loci longer than 131 kb; no plant species met the 20-locus threshold in this range.

**Supplementary Figure A.10.**
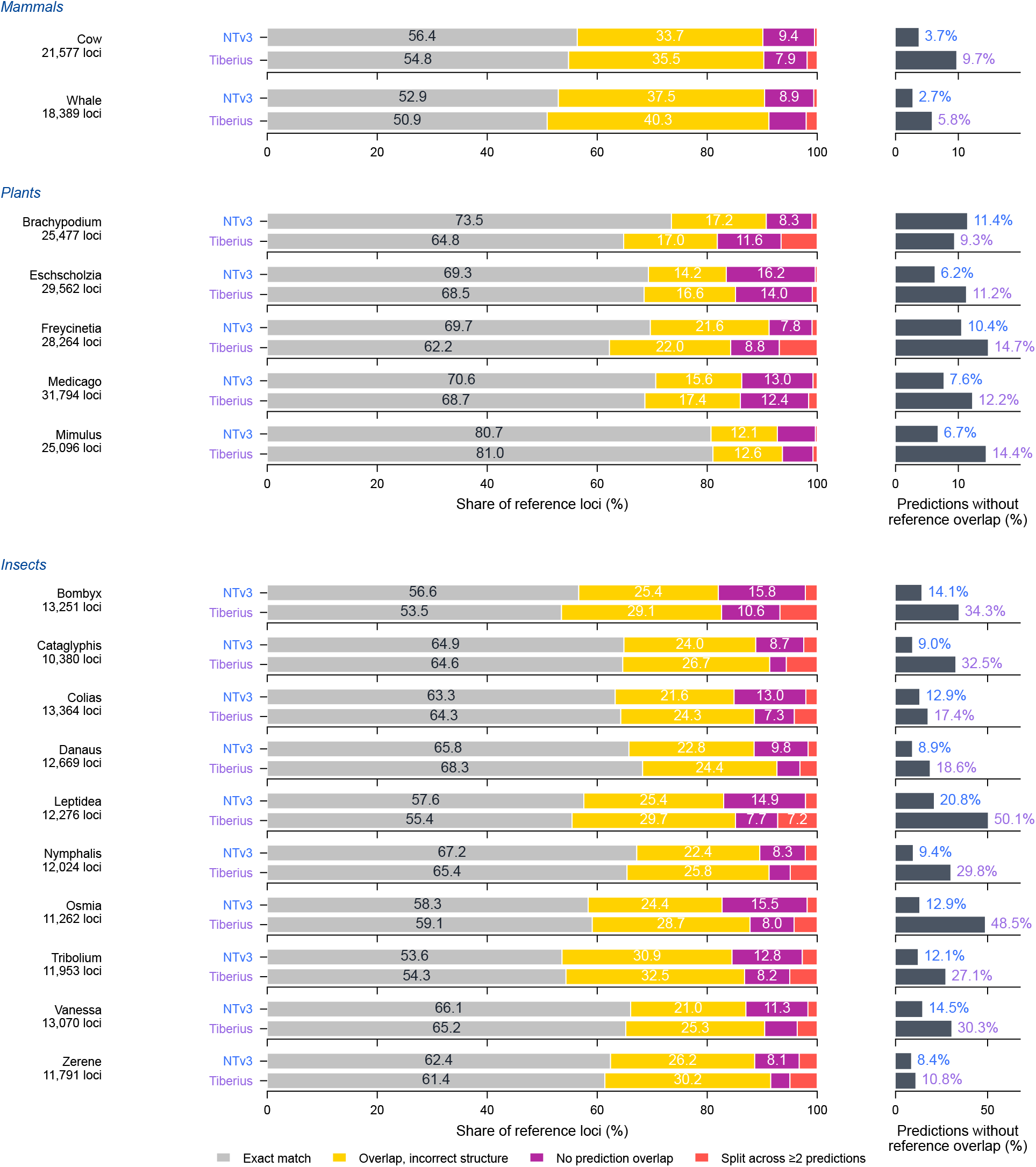
Locus-level annotation outcomes across mammals, plants and insects. Outcomes for two held-out mammals and five plants (**a**) and ten held-out insects (**b**). For each caller, reference loci are partitioned into four mutually exclusive categories: recovered exactly, overlapped by a prediction with an incorrect structure, not overlapped by any prediction, or split across two or more predictions. Right-hand panels show predictions without overlap to a reference locus as a proportion of all predicted gene models; these panels therefore use a different denominator and are scaled independently. Numbers beside species names give the reference-locus counts. All NTv3 results use the released retained fraction, *ρ* = 0.75 (Methods).

**Supplementary Figure A.11.**
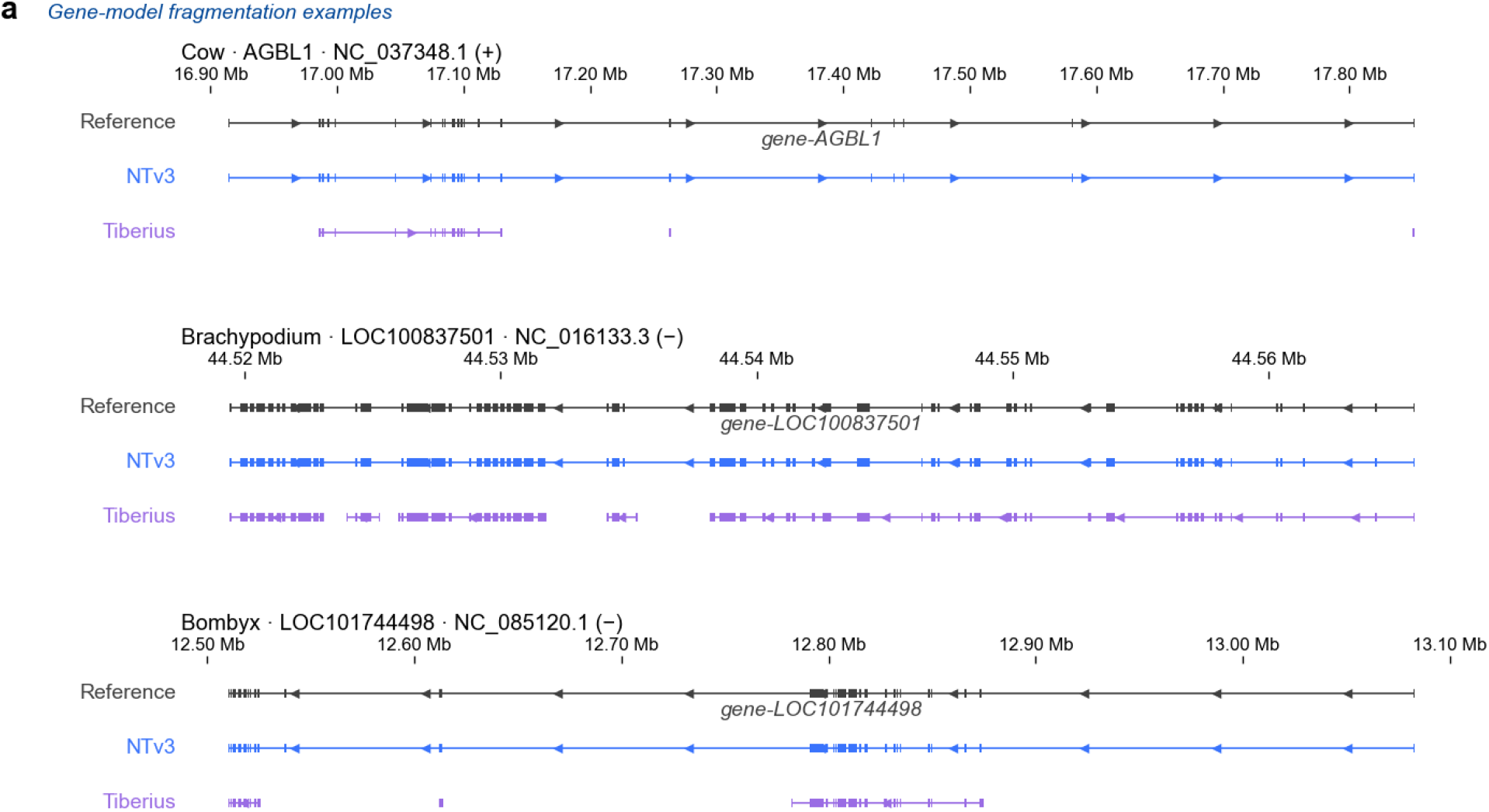
Examples of reduced gene-model fragmentation across clades. Reference, NTv3 and Tiberius gene models at one locus from each of three clades: cow *AGBL1* (936.3 kb; 23 coding exons), *Brachypodium LOC100837501* (46.3 kb; 64 coding exons) and *Bombyx LOC101744498* (572.2 kb; 53 coding exons). At these loci, Tiberius produced three, five and three gene models, respectively, whereas NTv3 produced one model and reproduced all coding exons. Examples were selected by the rule that one caller split the locus while the other produced a single exact model. Counts printed below each locus report the number of loci satisfying this rule in the displayed and reciprocal directions: 117 versus 9 in cow, 939 versus 5 in *Brachypodium*, and 244 versus 19 in *Bombyx*. The *Brachypodium* example illustrates fragmentation of a short, exon-dense locus.

**Supplementary Figure A.12.**
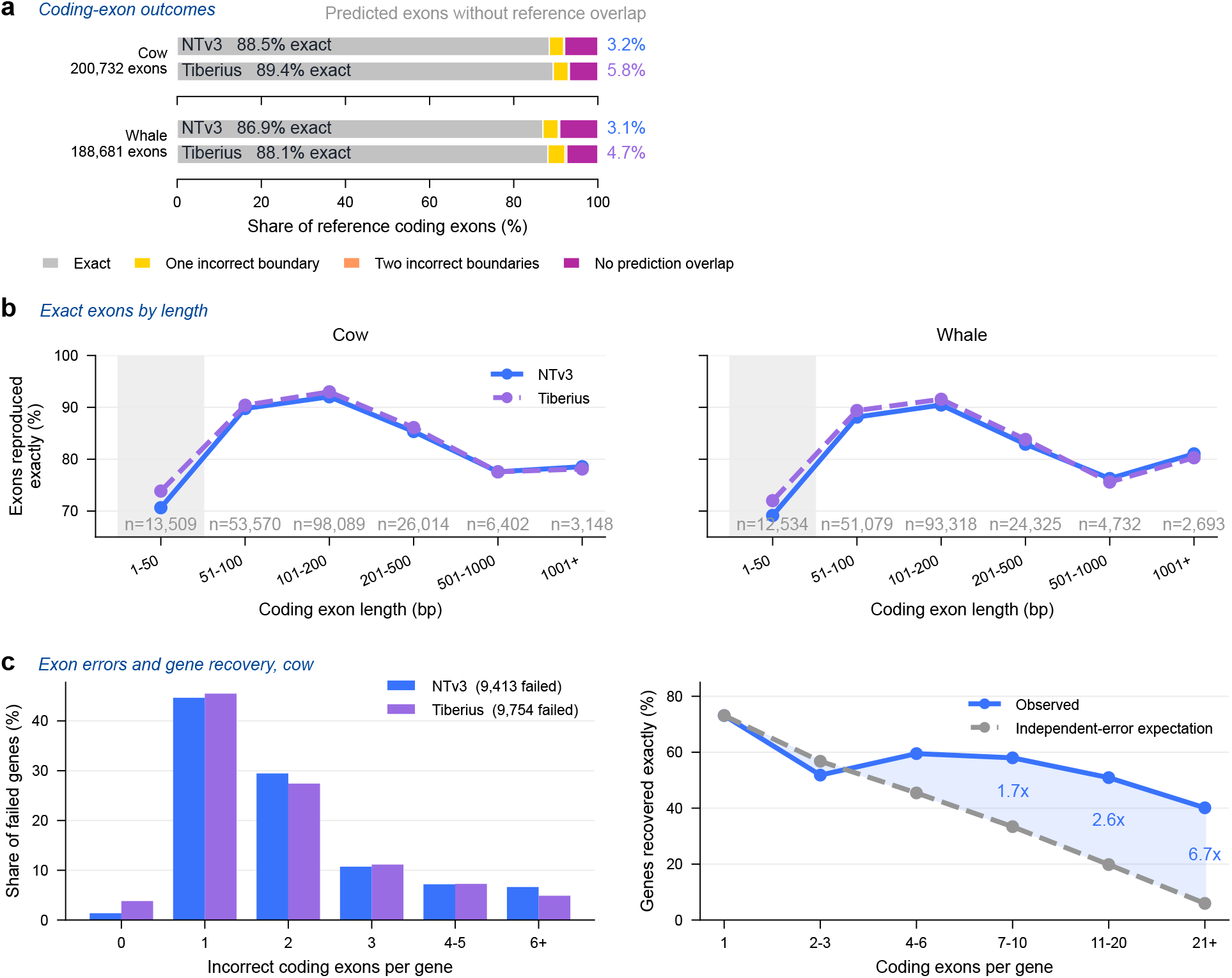
Exon-level error patterns and their relation to gene-level accuracy. Analyses of cow and beluga whale annotations. **a** Reference coding exons classified as reproduced exactly, having one incorrect boundary, having both boundaries incorrect, or having no overlapping prediction. Values at right give the proportion of predicted coding exons with no reference overlap. The evaluation comprises 200,732 cow exons and 188,681 beluga whale exons. **b** Proportion of coding exons reproduced exactly as a function of exon length. Shading denotes exons of at most 50 bp; sample sizes are shown below each bin. **c** Left, number of incorrectly predicted coding exons among failed cow gene models (9,413 for NTv3 and 9,754 for Tiberius). Right, observed exact gene recovery as a function of coding-exon count compared with the recovery expected if exon errors occurred independently. Ratios are shown where observed and expected recovery are both non-zero.

**Supplementary Figure A.13.**
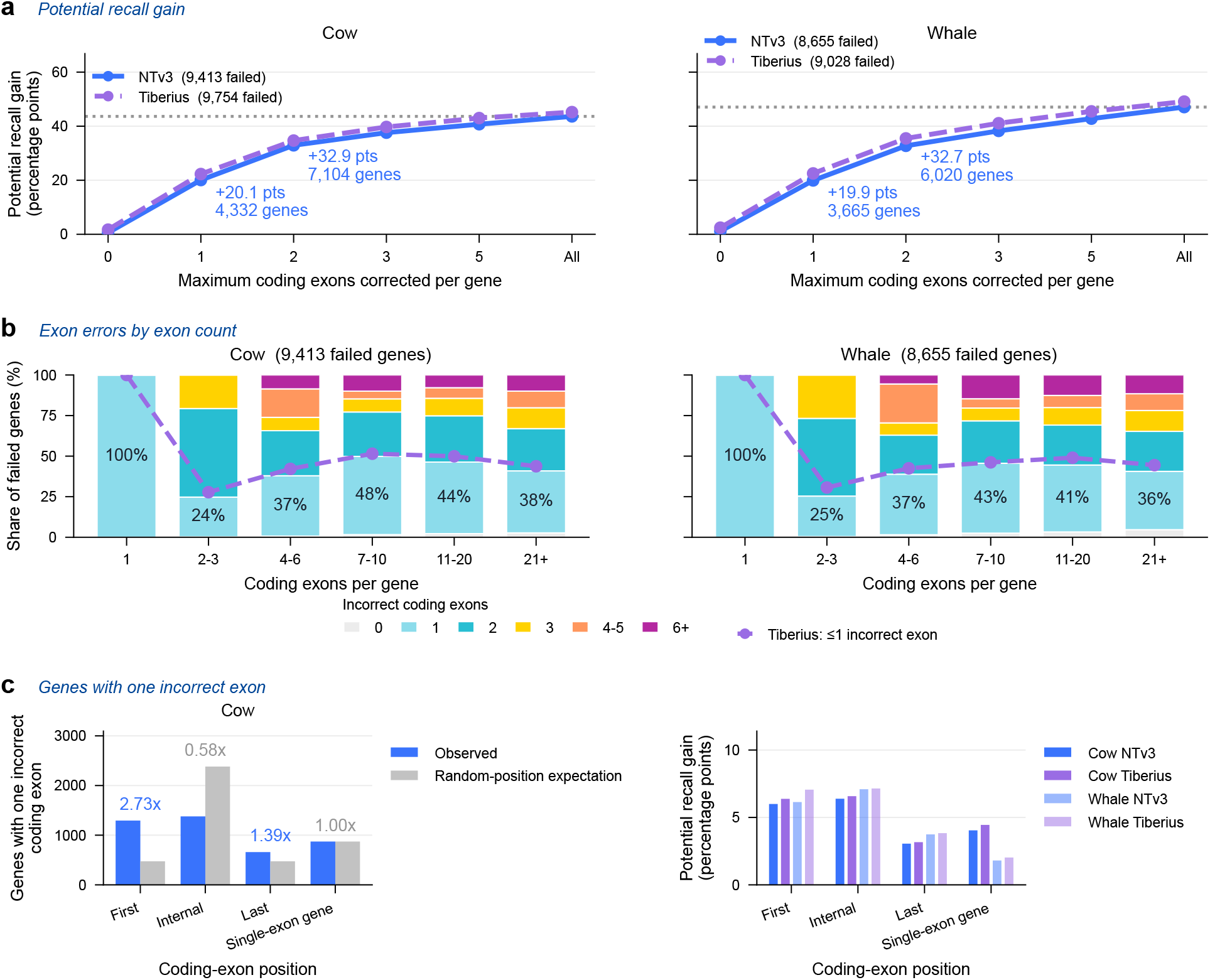
Exon errors underlying failed gene models. **a** Cumulative gene recall that would be obtained by correcting up to the indicated number of coding exons per failed gene model. The dotted line marks the maximum obtained if all failed models were corrected. **b** Distribution of the number of incorrect coding exons in failed NTv3 gene models, stratified by the number of coding exons in the reference gene. The dashed line gives the corresponding cumulative proportion of Tiberius failures at the upper boundary of the one-error category; percentages within the bars give this quantity for NTv3. **c** Left, observed position of the error among cow genes that fail because of exactly one coding exon, compared with a random-position expectation. Right, potential increase in gene recall from correcting each exon-position class alone, shown for both callers and both species.

**Supplementary Figure A.14.**
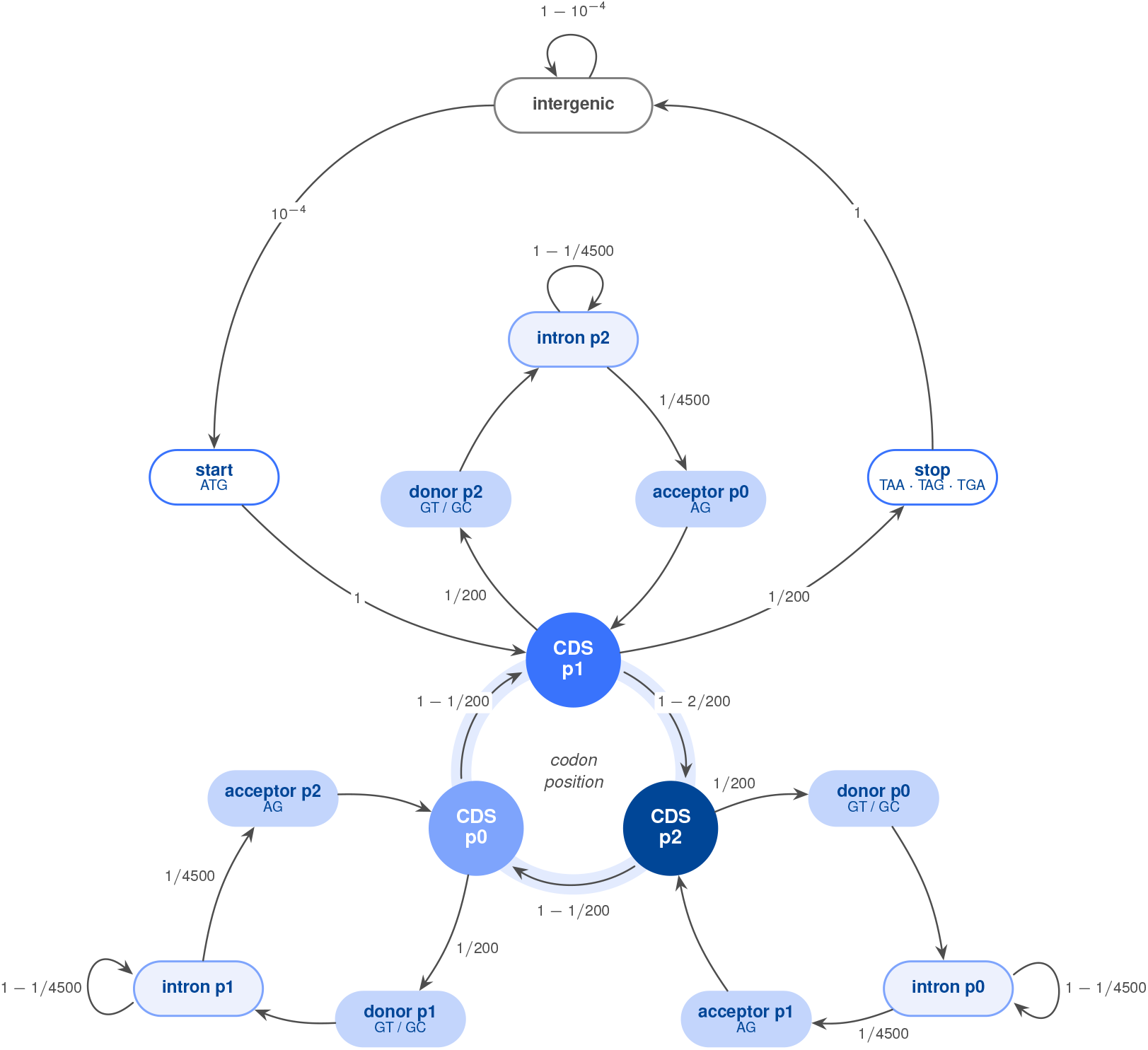
State grammar of the annotation decoder. The fifteen decoder states and every non-zero transition probability; omitted transitions are zero (Supplementary Note B.6). The three coding states form a codon-position cycle with no self-loop (center). Each intron detour leaves `CDS_p`*N* through `donor_p`(*N*+1) and returns to it through `acceptor_p`(*N*+2), phases modulo 3, so frame is preserved across an intron of any length. Genes are entered and left through `CDS_p1` only. Motifs under the start, stop and splice states are the sequence constraints applied to their emissions. Not drawn: the nine minor-spliceosome (U12) states, which mirror the donor, intron and acceptor states phase for phase and take 0.105% of each donor transition. Each strand is decoded separately.

**Supplementary Figure A.15.**
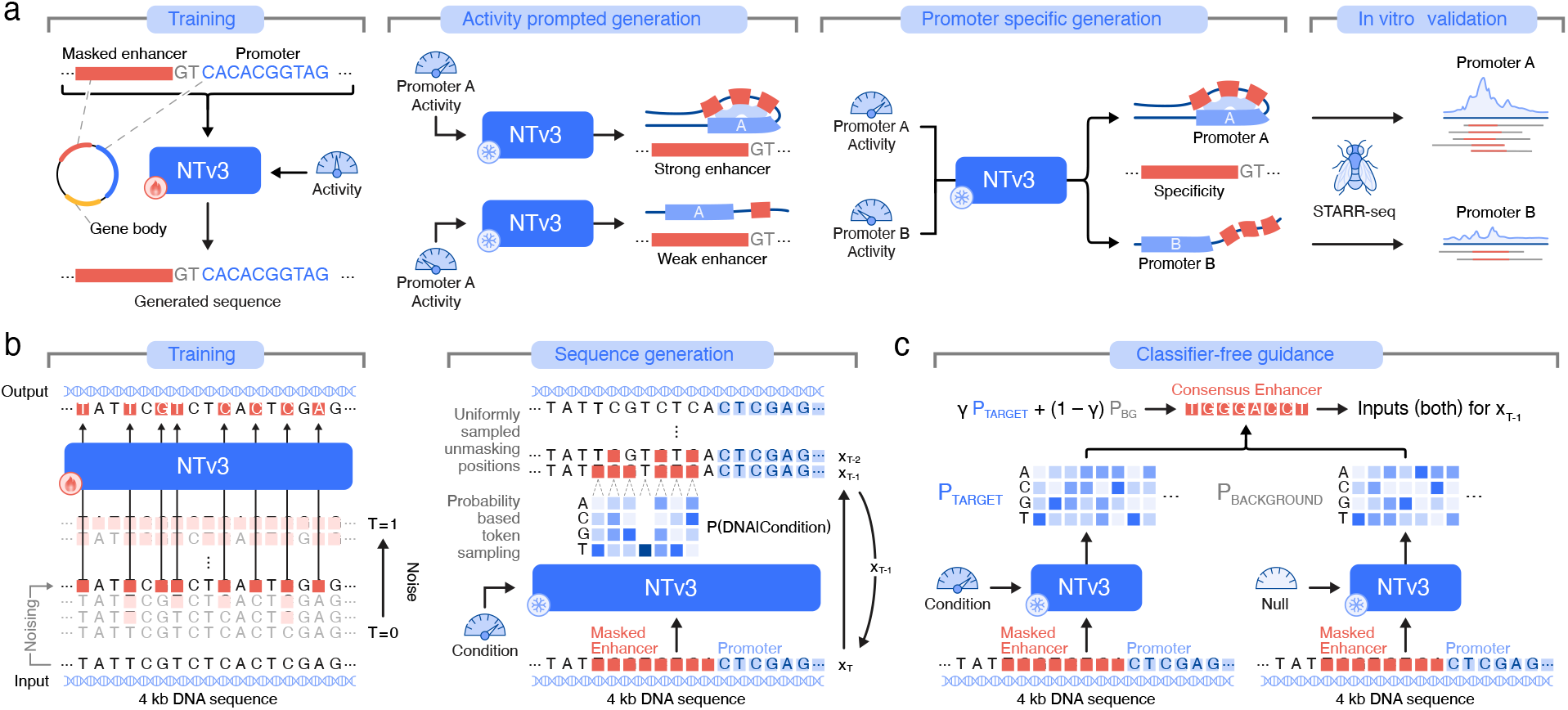
Detailed masked-diffusion workflow for *Drosophila* enhancer design. **a** Detailed schematic of the *Drosophila* S2-cell enhancer-design workflow summarized in Fig. 6a. A task-specific NTv3 fine-tune initialized from the post-trained NTv3 checkpoint supports activity-conditioned and promoter-specific enhancer design, with generated sequences subsequently evaluated by STARR-seq in S2 cells. **b** Masked diffusion language modeling (MDLM) enables generative fine-tuning of the NTv3 backbone. During training, variable fractions of the enhancer interval are masked; during generation, the masked interval is iteratively reconstructed conditional on the requested activity and reporter context. **c** Classifier-free guidance combines target-conditioned and background-conditioned denoising predictions to steer sampling toward the requested activity and promoter context.

**Supplementary Figure A.16.**
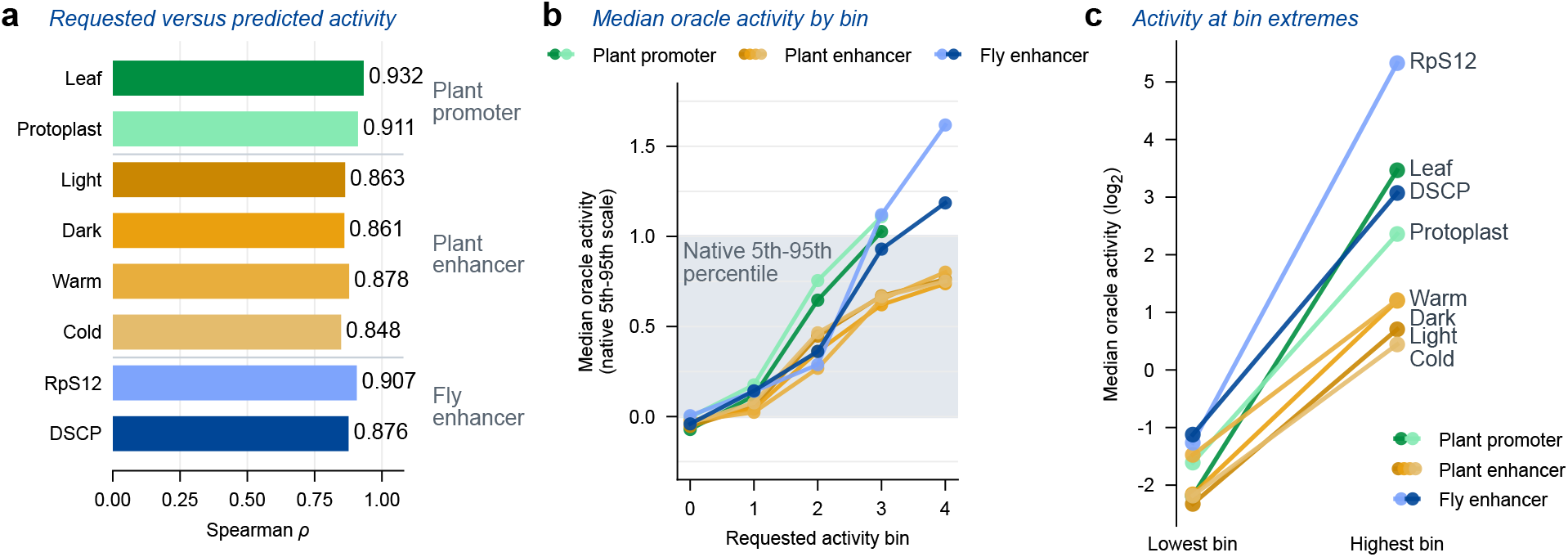
Control of oracle-predicted activity across design contexts. **a** Spearman correlation between requested activity bin and oracle-predicted activity for eight promoter or enhancer design contexts. **b** Median oracle-predicted activity across requested bins. Values are expressed relative to the corresponding native 5th–95th-percentile range where a native pool is available. **c** Median oracle-predicted activity in the lowest and highest generated bins for each context, shown in raw log_2_ activity units. The highest plant-promoter bin is excluded because it contains no training examples. The protoplast promoter condition shown here is distinct from the separate maize protoplast enhancer assay, for which no sequence pool was generated.

**Supplementary Figure A.17.**
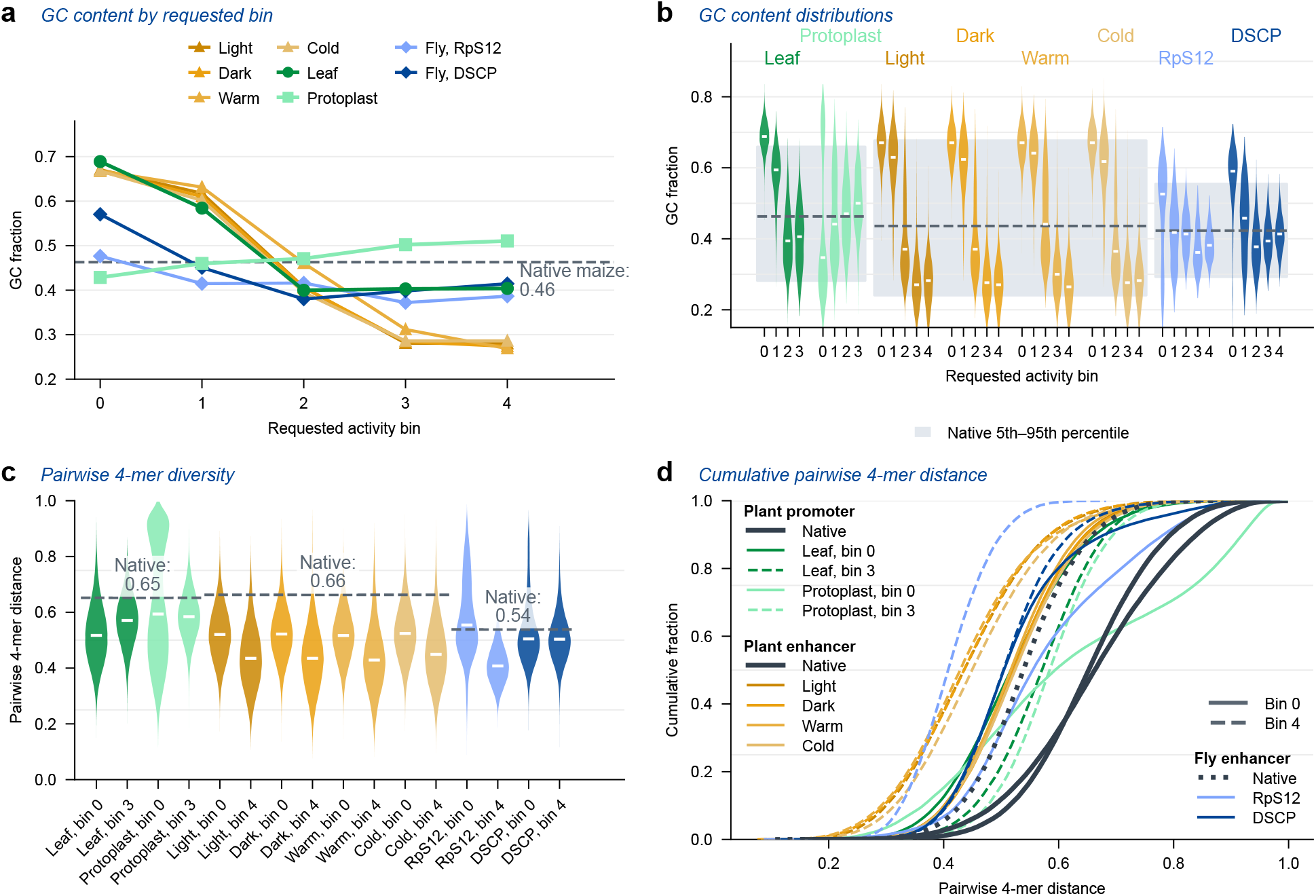
Sequence composition and diversity across requested activity levels. **a** Median GC content of generated sequences as a function of requested activity bin for each design context. The dashed line denotes the pooled native median. **b** GC-content distributions for the same pools; shaded regions show the corresponding native 5th–95th-percentile ranges. **c**,**d** Pairwise 4-mer distances within native and generated sequence pools, shown as distributions (**c**) and empirical cumulative distributions (**d**). Larger values indicate greater within-pool sequence diversity. Generated pools from the lowest and highest activity bins are shown in **c**; all displayed conditions are shown in **d**. No generated pool contained duplicate sequences.

**Supplementary Figure A.18.**
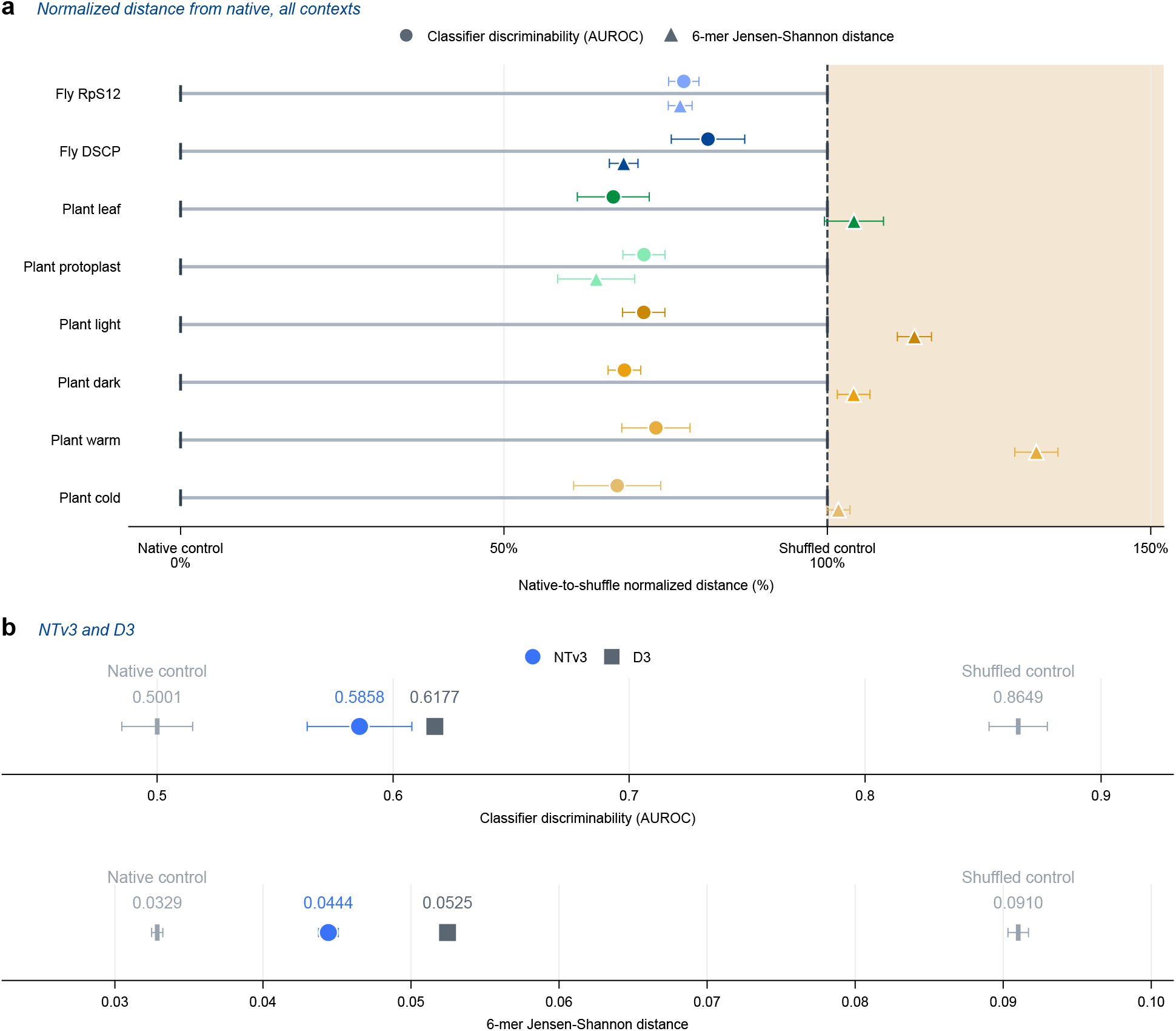
Generated-versus-native sequence similarity across design contexts. Benchmark underlying Fig. 6d and Fig. 6e. Each generated pool is compared with a held-out native sequence pool and with a dinucleotide-shuffled version of that pool. **a** Classifier discriminability (auROC; circles) and 6-mer Jensen–Shannon distance (triangles) for eight design contexts, normalized between the native-versus-native control (0%) and the native-versus-shuffled control (100%). Lower values indicate greater similarity to native sequence; the shaded region marks values exceeding the shuffled control. **b** The same metrics in their original units for the fly benchmark, comparing NTv3 and D3-Tran with native-versus-native and native-versus-shuffled controls. For auROC, 0.5 denotes chance discrimination; for Jensen–Shannon distance, 0 denotes identical 6-mer composition. Points show means and error bars show standard deviations across five independent draws for NTv3 and the native and shuffled controls.

**Supplementary Figure A.19.**
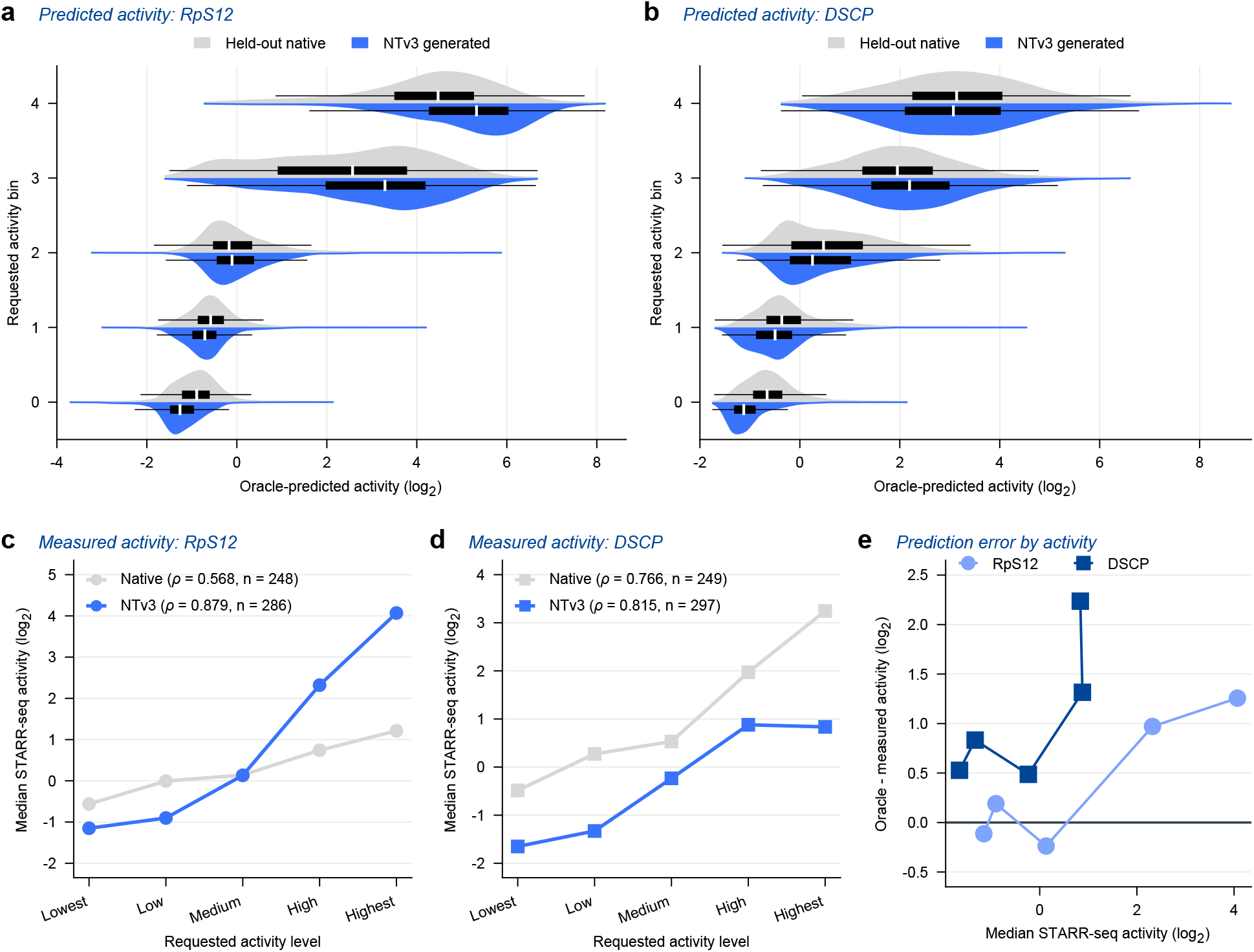
Oracle predictions and experimental validation of fly-enhancer activity. **a**,**b** Distributions of oracle-predicted activity for held-out native enhancers and NTv3-generated enhancers across requested activity bins in the RpS12 (**a**) and DSCP (**b**) promoter contexts. **c**,**d** Median STARR-seq activity measured in *Drosophila* S2 cells for native and generated enhancers across the same requested bins in the RpS12 (**c**) and DSCP (**d**) contexts. Spearman correlations and sample sizes are shown in the panels. **e** Difference between oracle-predicted and measured median activity as a function of measured STARR-seq activity. At the highest activity level, the oracle overestimates measured activity by 1.26 and 2.24 log_2_ units in the RpS12 and DSCP contexts, respectively.

**Supplementary Figure A.20.**
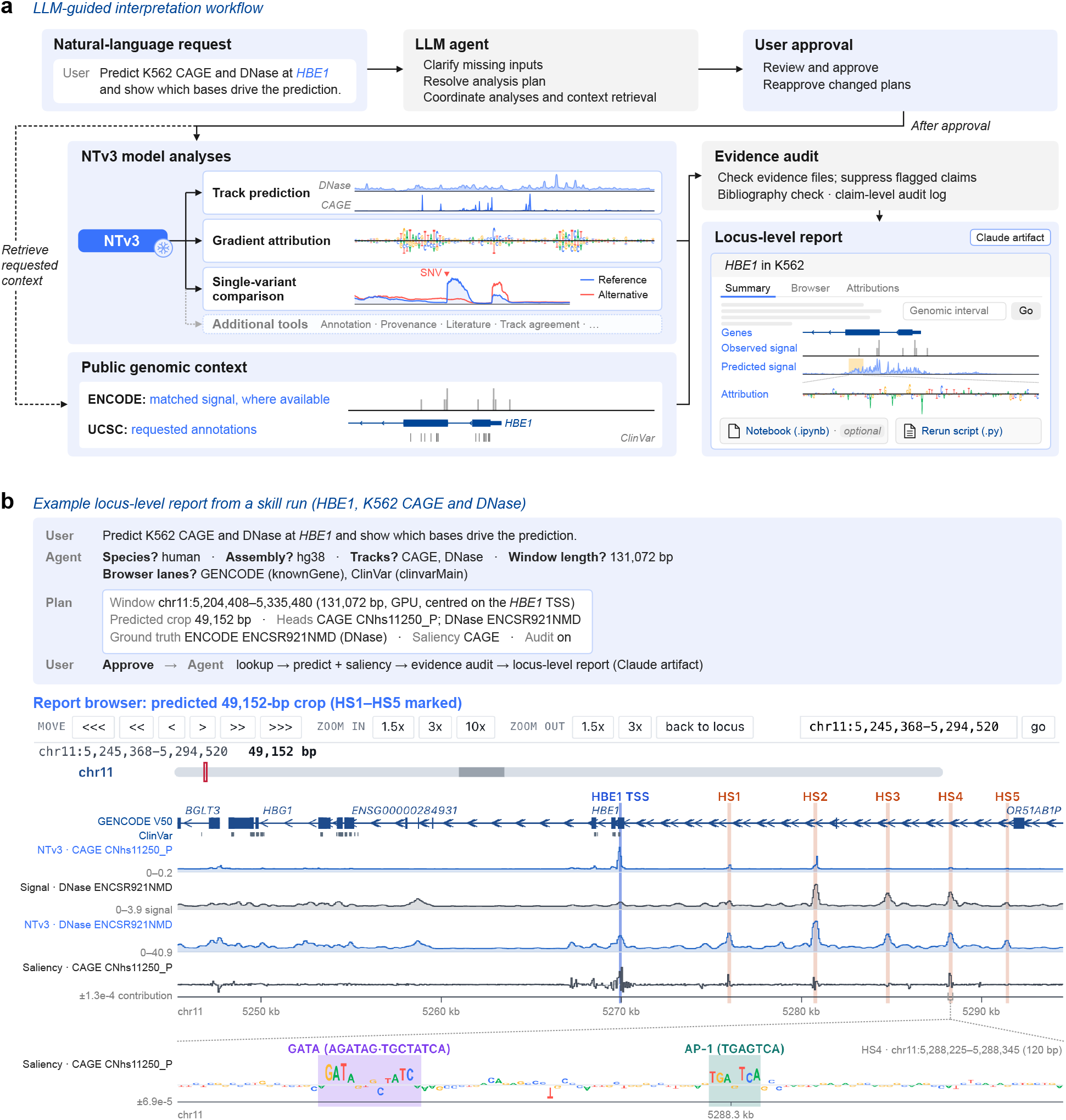
LLM-guided interpretation workflow and example locus-level report. **a** Overview of the agent-guided workflow. Following clarification of missing inputs and user approval of the analysis plan, the agent coordinates NTv3 predictions, gradient-based attribution and single-nucleotide variant comparisons, and retrieves matched ENCODE measurements, where available, and requested UCSC annotations. Model weights remain fixed during analysis. Results are assembled into a locus-level report with an interactive genome browser, a rerun script and an optional notebook. Report claims are screened for evidence-file references, bibliography entries and predefined overstatement criteria, with decisions recorded in an audit log. **b** Example report for the human *HBE1* locus in K562 cells (hg38). Top, clarification and approval of a 131,072-bp input window for CAGE and DNase prediction and CAGE attribution. Middle, genome-browser view of the central 49,152-bp prediction region, showing GENCODE and ClinVar [109] annotations, predicted CAGE and DNase signal, measured DNase signal (ENCODE ENCSR921NMD), and CAGE attributions. The *HBE1* transcription start site (TSS) and DNase hypersensitive elements HS1–HS5 are indicated. Bottom, a 120-bp attribution map within HS4, highlighting GATA- and AP-1-like sequences. Letter heights represent reference-base attribution scores.

**Supplementary Figure A.21.**
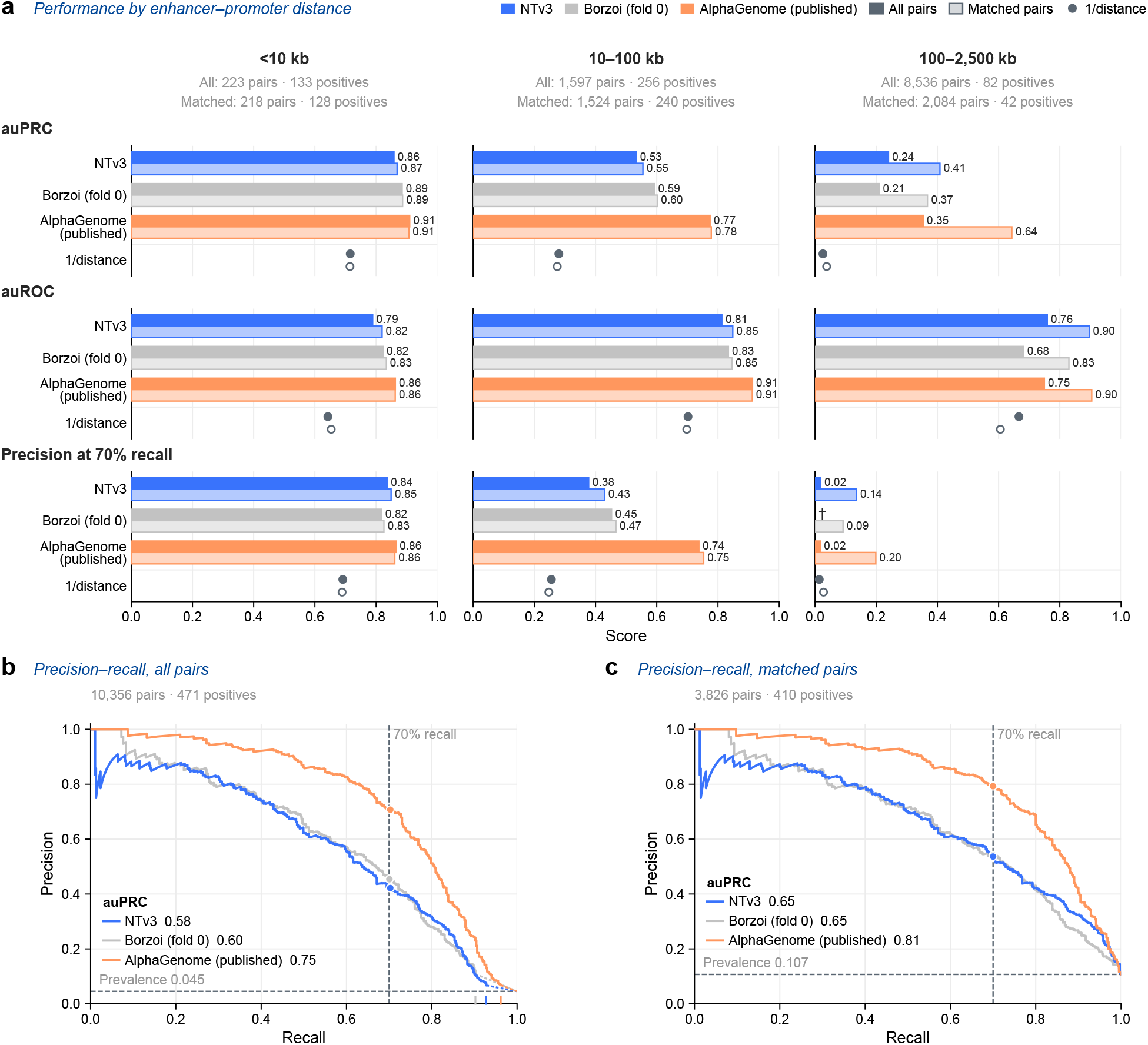
Enhancer–gene linking performance across distance bins. **a** auPRC, auROC and precision at 70% recall for NTv3, Borzoi fold 0 and published AlphaGenome scores on K562 CRISPRi enhancer–gene pairs from ENCODE-rE2G, stratified by enhancer-to-TSS distance. Solid bars show all 10,356 pairs (471 positives), with pairs a model did not score set to zero; tinted bars show the 3,826 matched pairs (410 positives) scored by all three models. Filled and open points show the reciprocal-distance (1/distance) baseline on all and matched pairs, respectively. Numbers above each column give pairs and experimentally supported positives. Values at the bar ends give the metric. *†* Precision at 70% recall is not defined: the model’s non-zero scores cover fewer than 70% of the positives in this bin (43 of 82), the remaining pairs being outside its input window and tied at zero. **b**,**c** Precision–recall curves for all pairs (b) and matched pairs (c), pooled across distance bins. Points mark precision at 70% recall (vertical dashed line); the horizontal dashed line indicates positive-class prevalence; legends give pooled auPRC. In b, dotted segments span the block of pairs tied at zero, and ticks on the x-axis mark the recall at which each model’s non-zero ranking ends (Borzoi 0.90, NTv3 0.93, AlphaGenome 0.96). The matched pairs in c contain no tied-at-zero scores.

### B. Supplementary Notes

These notes carry the implementation detail that the Methods reference but do not need to state in full: the species-conditioning mechanism, the token distribution behind the sequence-length curriculum, the training infrastructure and throughput optimizations, and the per-assay provenance of the post-training track and NTv3 Benchmark corpora.

#### B.1. Species Conditioning

NTv3 predicts functional tracks for fifteen genomes from a single set of weights. It does so with a species-conditioned output head rather than a species-conditioned trunk: the convolutional encoder, transformer tower and decoder see DNA sequence alone, and species identity enters only where the trunk representation is projected onto track space. The Methods (Section 4.1.1) give the form of that conditioning; this note records the implementation detail behind it.

Each conditioning variable has its own learned embedding table. For the released model the only variable is species, with an embedding dimension of *d*_cond_ = 32 over the fifteen training genomes. The embedding *e*_*s*_ enters the head along two independent paths, both of which are feature-wise linear modulations (FiLM) [110, 111].

##### Input-side modulation

The base-resolution trunk representation is layer-normalized and then scaled and shifted per channel, with the scale and shift produced from *e*_*s*_ by a linear projection. Writing *h*_*i*_ for the trunk representation at position *i* and *d* for its width,

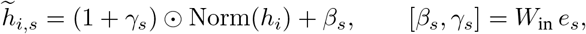

with 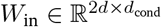. The modulated representation then passes through a single linear projection shared by every species.

##### Output-side modulation

A second zero-initialized projection maps *e*_*s*_ to a per-track additive bias applied to the raw logits, before the softplus that makes predictions non-negative. This gives each genome a learned offset per track without giving it its own projection matrix.

Both projections are initialized to zero, so at initialization the scale is one, the shift and bias are zero, and the conditioned head reduces exactly to an unconditioned normalized linear head. This is what permits post-training to warm-start from the pre-trained backbone without the randomly initialized conditioning perturbing the representation on the first step.

Because a single projection is shared across genomes, its output columns are shared *positionally*: column *t* is not a canonical assay, and the assay it corresponds to differs between genomes. Genomes with fewer tracks than the widest genome are zero-padded to the shared width and the padded columns are excluded from the loss by a per-genome mask, so the padding contributes no gradient.

The implementation also supports conditioning the trunk itself, in which the same FiLM mechanism modulates the transformer pre-normalizations and the decoder convolution blocks. This was disabled for the released model, and the convolutional encoder is never conditioned in either configuration.

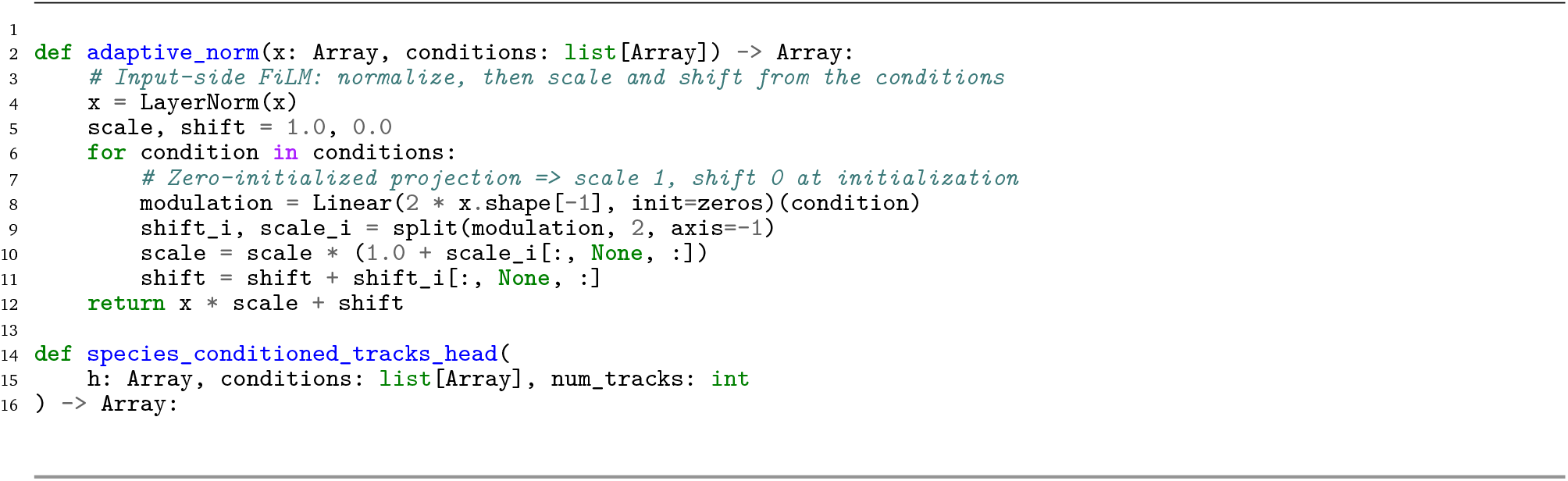
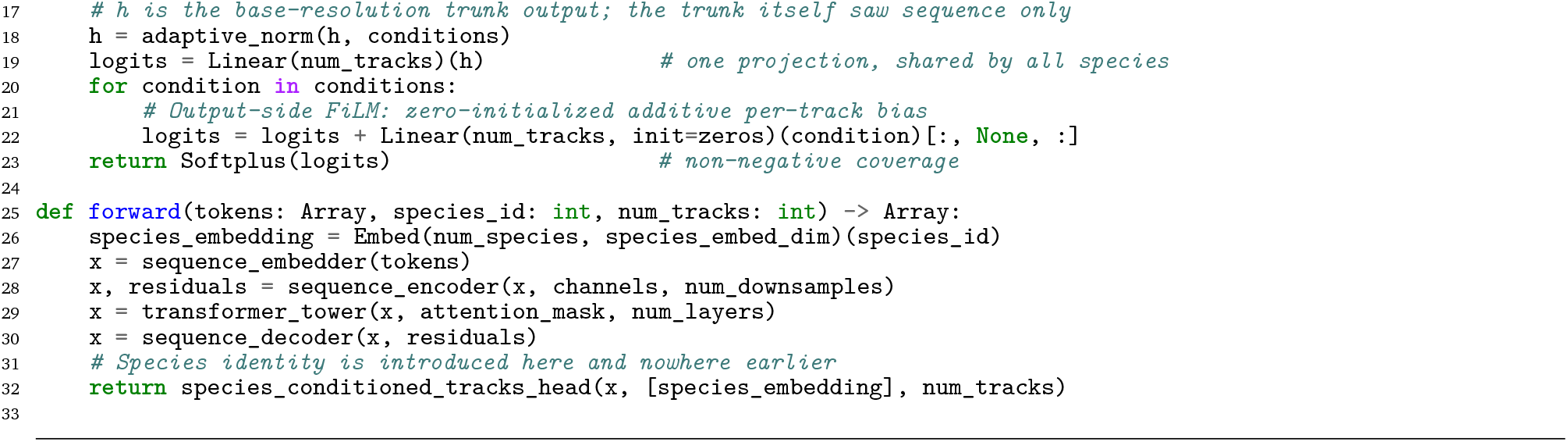

#### B.2. Sequence-Length Extension Token Distribution

After 8kb-training, we extended the model’s context length from 8,192 bp to 1 Mb through a staged curriculum. Approximately every 50B tokens, we introduced a new maximum sequence length, doubling it at each stage (16 kb, 32 kb, 64 kb, …, 1 Mb). Throughout this process, all shorter sequence lengths remained in the training mixture, allowing the model to acquire long-range dependencies gradually while retaining strong performance on short contexts.

As in 8kb-training, we created a distinct data source for each sequence length. For subsets composed of inherently short sequences (e.g., transcripts, metagenomes, and prokaryotic genomes), sequences were assigned to datasource_*i*_ if their length satisfied

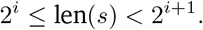

As the maximum context length grew through the stages, the set of active bins expanded from datasource_14_ up to datasource_20_.

Whole eukaryotic genomes (`ncbi_eukaryotic_genomes`) required a separate strategy, as they constitute the majority of tokens and are often extremely long, with many sequences exceeding one megabase. We introduce them earlier in training, at the 32-kb stage, to allow the model more exposure, particularly to smaller chunks that may be more representative of practical downstream applications. Sequences shorter than 32 kb within `ncbi_eukaryotic_genomes` account for 0.8T of its 6.98T nucleotides, leaving 6.2T for the remaining length extension phase. We chose not to include these shorter length whole genome sequences in the early stages of training to focus short context training on functionally enriched areas.

Starting at the 32 kb stage, each eukaryotic genome was subdivided into non-overlapping windows matching the stage’s target length, and all such windows were added to the corresponding datasource_*i*_. In other words, for datasource_*i*_ with target length 2^*i*^ bp, every 2^*i*^-bp chunk from genomes at least 2^*i*^ bp long was included. This ensured that the model saw whole-genome sequences progressively, integrating both shorter and longer contexts.

For example, during the 64 kb stage, datasource_16_ included (i) all non-eukaryotic sequences between 64 kb and 128 kb, and (ii) all 64 kb chunks extracted from eukaryotic genomes at least 64 kb in length, including megabase-scale chromosomes. This staged inclusion allowed the model to encounter whole-genome structures earlier in training rather than only in the final stage.

At each stage, the primary focus was to adjust the model to the newly introduced longest sequences. To maintain robust performance across all context lengths, we dynamically adjusted the sampling weights of each data source. The newly introduced longest sequence length was assigned approximately 40% of the sampling probability, while the remainder was rescaled among all shorter bins. To prevent shorter contexts from being underrepresented, we enforced a minimum sampling probability of 5% for every bin; if rescaling caused a bin to fall below this threshold, it was set to 5%, and the remaining weights were re-normalized. The evolution of these sampling distributions is detailed in Table 1. We provide below the basic implementation for generating this sequence length extension schedule, given an initial distribution over datasets, a number of stages to extend for, and the new weight with which to fold in the longest sequence. The pseudocode gives the schedule in its basic form; the deployed schedule additionally retires the shortest bins once the context has grown far beyond them, which is why the 1 kb and 2 kb bins reach zero from stage 5 onward in Table 1 rather than remaining at the 5% floor, and why the realized weights differ from those the code below produces.

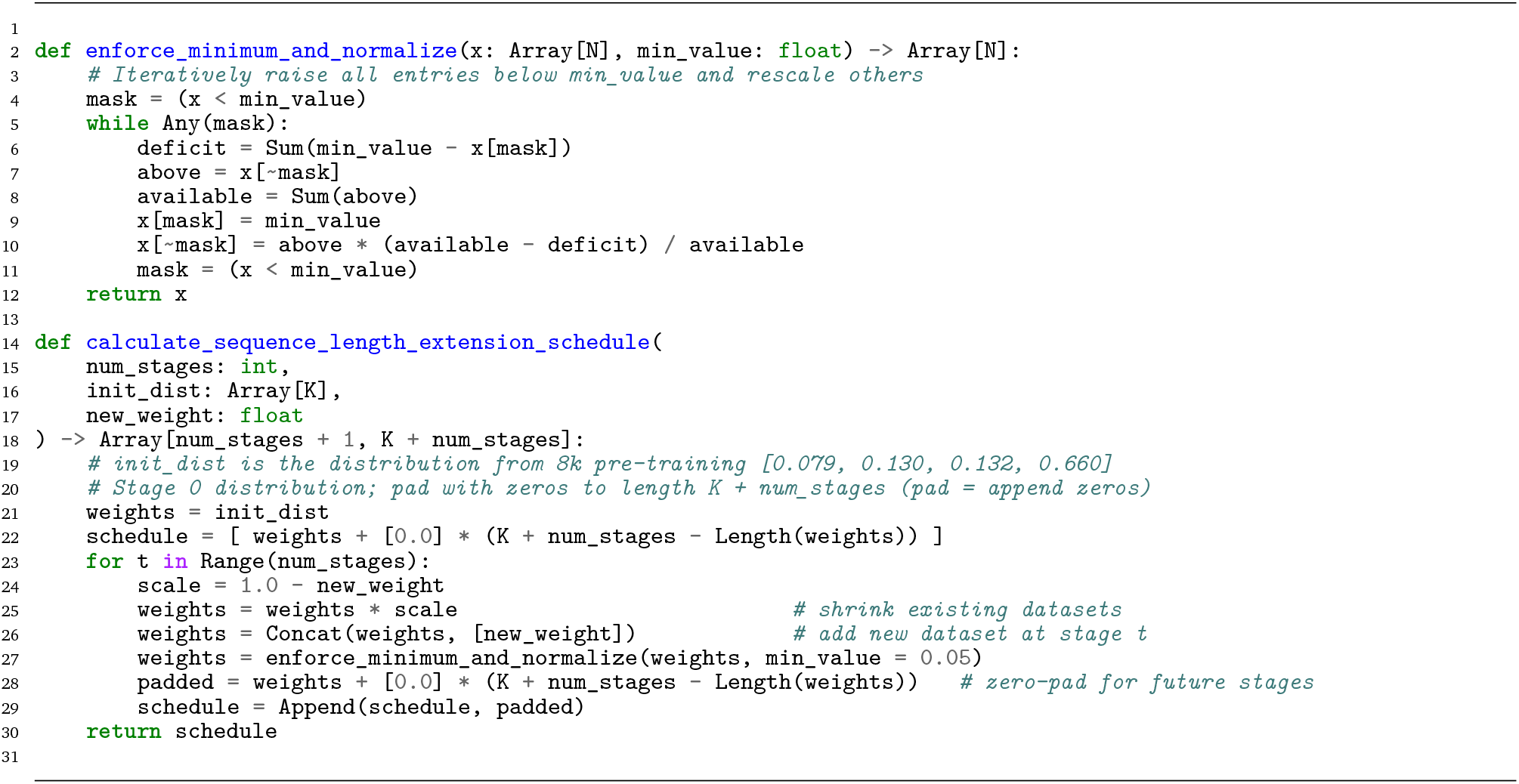

#### B.3. Training Infrastructure and Optimization

##### B.3.1. Data Loading for Pre-training

To efficiently manage large-scale genomic datasets comprising millions of individual FASTA files, we implemented a scalable data loading pipeline designed to maintain a low memory footprint and minimize disk I/O, independent of dataset size. The system enables direct, byte-level random access to genomic subsequences without loading full files or global indices into memory.

The approach consists of two stages: (1) a single-pass metadata indexing stage and (2) a byte-addressable data retrieval stage.

During the metadata indexing stage, each FASTA file is scanned once to extract attributes required for random access, including total sequence length, header byte length, and per-line sequence length. To eliminate the memory burden associated with tracking millions of FASTA files, we serialize all file paths and metadata into disk-backed, memory-mapped `array_record` databases, ensuring that metadata lookup does not scale RAM usage with dataset size. This design externalizes metadata management from system RAM and supports efficient, persistent lookup across arbitrarily large datasets.

In the retrieval stage, the complete corpus is treated as a virtual contiguous sequence that is partitioned into fixed-length segments used as atomic training units. Each segment is addressable by a unique global index. Given a segment index *i*, the corresponding source file is identified via a binary search over a precomputed array of cumulative segment counts. The global coordinates of the segment are then mapped to nucleotide offsets within the identified file. These offsets are translated into absolute byte positions using the metadata. A single read operation is issued to retrieve only the necessary byte range, which is then decoded directly into sequence data.

##### B.3.2. Data Loading for Genomic Tracks Post-training

The post-training data comprises species genomes and functional tracks. The species genomes are moderately sized and can be easily loaded for training. In contrast, the functional track dataset is large with data from 16,262 tracks.

The `tensorstore` Python package was used to store and load the functional track data. The data is stored in an S3 bucket as tensorstore arrays with datatype of float16. A separate tensorstore is used for each species. The data is compressed with zstd level 3 compression, which takes advantage of the sparsity of elements. To enable fast loading of all the tracks in the tensorstore, the data is compressed in chunks of 8192 bp by the full number of tracks used for that species.

Each target example for training the functional track head is substantially large. For instance, a single human sample comprising 7,362 functional tracks over a 393 kb window corresponds to approximately 6 GB of data in memory. To efficiently stream these large tensors during training, we employ the Grain Python framework with parallel data-loading across multiple worker processes. This design minimizes input latency and sustains high-throughput training by ensuring that model computation is rarely idle while awaiting sample retrieval.

##### B.3.3. Training Optimizations

Training was conducted using multi-node setups on our internal Kyber cluster and on TPUv6 hardware. Distributed training was used for scaling to long sequences, with the largest models requiring sequence parallelism across one or more nodes of 8 GPUs. We applied a series of optimizations to accelerate training while preserving performance. We provide an overview of these optimizations below.

###### Efficient Up-sampling Implementation

To accelerate the up-sampling path, we replaced conventional transposed convolutions (Conv1DTranspose) with a **repeat+conv** approach. In this implementation, a 1D convolution is first applied at the input resolution; the sequence is then up-sampled using nearest-neighbor repetition along the sequence length, duplicating each position by the up-sampling factor, and a second 1D convolution is applied to the repeated sequence to mix features locally. Nearest-neighbor repetition is computationally cheap because it involves only memory replication, and the subsequent convolution operates at stride 1. This design reduces the cost of up-sampling while maintaining the function of conventional transposed convolutions.

###### Sequence Parallelism

To scale our models to very long sequences, we implemented sequence parallelism, in which each batch is split along the sequence dimension across multiple devices. This approach allows each device to process a contiguous portion of the sequence, reducing memory requirements while enabling training on sequences up to 1 Mb.

Certain operations require additional communication between devices to maintain correctness. We express the split declaratively, by annotating the sequence axis of the activations with its sharding, and the compiler’s SPMD partitioner inserts the required collectives. For convolutional layers this is a collective permute that shares the boundary tokens at the edges of each local sequence partition. For attention layers the key and value tensors are all-gathered across devices while the query tensor remains sharded, so each device computes attention locally for its portion of the query while still attending to all keys and values across the full sequence.

This setup allowed us to train our largest models up to 1 Mb by splitting a single batch on the four GPUs used for post-training so we could benefit from the high interconnect between devices on the same machine without exceeding per-device memory limits.

##### B.3.4. Training Infrastructure

All models were trained on either Cloud TPUs (v6) through the TPU Research Cloud program or on-premises infrastructure. JAX let us target both accelerator types with only configuration changes to our training codebase. Our on-premises infrastructure is composed of machines with 8 H100 GPUs with 80 GB of memory connected via NVLink. The machines are connected with RoCE to allow for direct GPU-to-GPU communication across nodes. We run our experiments using AIchor, our internal ML training platform built on top of Kubernetes (https://aichor.ai/).

###### Pre-training Compute

The first pre-training phase accounted for the vast majority of compute: processing the full 5.6 trillion tokens took 4,119 H100 GPU-hours. By leveraging a 2-node (16 GPU) setup, the training wall-clock time was about eleven days (Table B.1). Sequence processing throughput was approximately 3.0M tokens/sec per node, or 6.0M tokens/sec in aggregate across the two nodes. In the final stages of sequence length extension we achieved lower throughput due to sequence parallelism. Despite this, the length extension stages remained short and therefore fast.

**Supplementary Table B.1.**
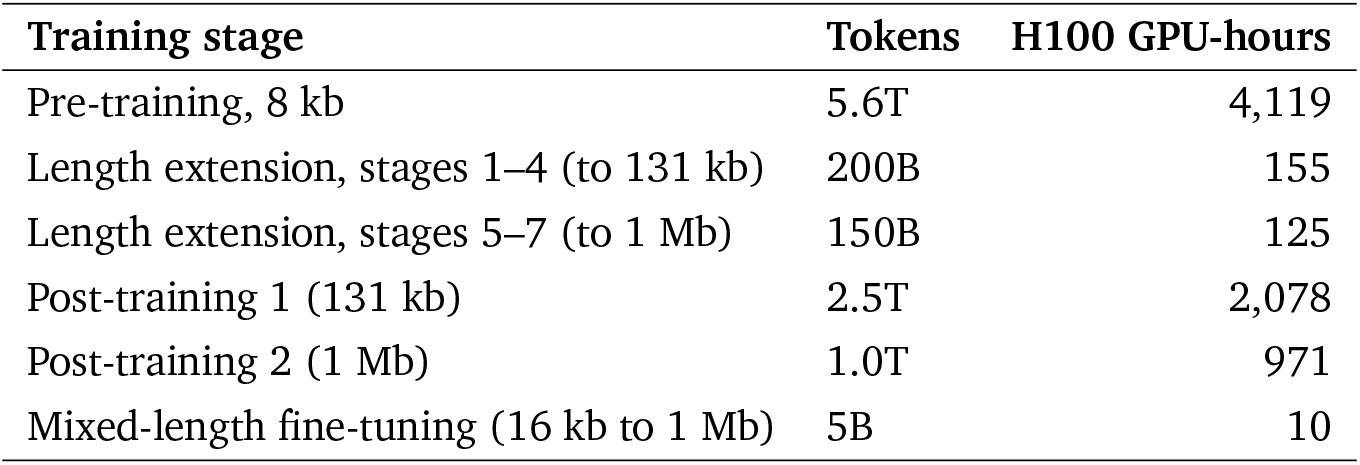
H100 GPU-hours required for each training stage of NTv3. Pre-training used 8 kb sequences, while length extension was split into stages 1–4 (up to 131 kb, full data parallelism) and stages 5–7 (262 kb to 1 Mb, requiring sequence parallelism). Post-training stage 1 used full data parallelism, whereas stage 2 required sequence parallelism across the four GPUs.

###### Post-training Compute

Post-training was split into two subphases to maximize efficiency and hardware usage. In the first subphase, models were trained on sequence lengths up to 131 kb using full data parallelism across four H100 GPUs of a single node: processing 2.5 trillion tokens took 2,078 GPU-hours, about three weeks of training wall-clock time, at approximately 1.3M tokens/sec (Table B.1). Using data parallelism enabled faster throughput due to reduced communication between GPUs. In the second subphase, the models were trained on longer sequences up to 1 Mb using sequence parallelism across the same four GPUs: processing 1 trillion tokens took 971 GPU-hours, about ten days, at approximately 1.1M tokens/sec. Using sequence parallelism reduced training throughput due to the higher communication demands.

#### B.4. Functional Track Data Sources and Processing

The post-training functional track corpus is summarized in Methods 4.3.1. This note gives the accessions, retrieval dates and processing pipeline for each species and assay type in that corpus.

##### B.4.1. Human and Mouse ENCODE Data

For both human and mouse, we used ENCODE v3 bigWig signal tracks corresponding to the exact sets included in Borzoi [5]. Data were retrieved directly from the ENCODE portal. When consortium-provided merged-replicate BigWigs were available, we used them preferentially; otherwise, we merged individual replicates ourselves to obtain unified signal tracks. All human ENCODE files were accessed on February 24, 2025, and all mouse ENCODE files on July 20, 2025.

##### B.4.2. Human and Mouse CAGE Data

CAGE datasets for both species were obtained from the FANTOM5 database, restricted to the standardized, reprocessed collections to ensure consistency across phases and assemblies. Specifically, we used the hg38 human CAGE data from https://fantom.gsc.riken.jp/5/datafiles/reprocessed/hg38_latest/basic/, and the mm10 mouse data from https://fantom.gsc.riken.jp/5/datafiles/reprocessed/mm10_latest/basic/. All files were accessed on July 24, 2025.

For both species, we used the `ctss.bed` files. Replicates were identified based on distinct `experiment_name` (e.g., CNhs11250) and `donor_id` (e.g., donor1, donor2). Each strand (+ and –) was processed independently. After download, we applied counts-per-million (CPM) normalization separately to each strand of each replicate. Replicates were then merged by averaging their normalized signal values, and the resulting processed files were converted to bigWig.

##### B.4.3. Human and Mouse GEO Data

For human and mouse functional tracks from GEO, we collected RNA-seq, ATAC-seq, DNase-seq, and ChIP-seq datasets and processed them using matching pipelines across species. GEO accessions were mapped to their corresponding SRA runs, and raw reads were retrieved and aligned using `bowtie2` [112] with default parameters. Human reads were aligned to GRCh38, and mouse reads to GRCm38/mm10. Resulting BAM files were coordinate-sorted using `samtools` [113]. Genome-wide coverage profiles were computed and normalized to CPM without additional filtering or duplicate removal. Only samples with complete metadata and successful processing were retained.

##### B.4.4. Drosophila Single-cell RNA- and ATAC-seq

For *Drosophila melanogaster*, we used the single-cell RNA-seq and ATAC-seq atlas of embryogenesis generated by Calderon et al. [70]. This dataset profiles nearly 1.5 million nuclei across the full 0–20 h developmental window and, through neural-network–based age inference, provides a continuous trajectory of chromatin accessibility and gene-expression dynamics with high temporal resolution. We extracted the published cell-type and developmental-age annotations and produced genome-wide RNA-seq and ATAC-seq coverage tracks by aggregating nuclei belonging to each annotated population. Following previous work [70], which treats NNLS mixture coefficients above 0.1 as high-confidence correspondence between RNA- and ATAC-derived cell states, we restricted our dataset to the 90 cell types exhibiting the strongest RNA–ATAC agreement. These high-quality, modality-matched tracks provide a temporally resolved view of transcriptional and regulatory activity throughout *Drosophila* embryogenesis.

##### B.4.5. Plant RNA-seq Tracks

The NCBI Sequence Read Archive (SRA) was queried for all RNA-seq experiments for six plant species: *Arabidopsis thaliana, Zea mays, Glycine max, Gossypium hirsutum, Triticum aestivum* and *Oryza sativa*. The metadata for the experiments was downloaded using Run Selector. Only experiments with `Treatment` and `Genotype` metadata were kept. To ensure the quality of the sequencing data experiments were filtered for `LibraryLayout` equal to `PAIRED`, `Bases` greater than or equal to 2.5 × 10^9^ and `AvgSpotLen` greater than or equal to 200 base pairs. Additional metadata in NCBI for the experiments was obtained by additionally querying for the biosample metadata associated with each experiment and for the bioproject data for each set of experiments. After the filtering for complete metadata and sequencing quality the number of experiments for each species was: *A. thaliana*: 8712, *Z. mays*: 3326, *G. max*: 2422, *G. hirsutum*: 902, *T. aestivum*: 2436 and *O. sativa*: 4410.

The NCBI experiments were organized into BioProjects, the experimental design of each BioProject was determined, and the experiments were filtered to select only a single biological/technical replicate per biological condition and to exclude mutants. Mutants were excluded since the genotype would differ from the genome DNA track for which the model would be trained to predict gene expression coverage. Additionally, the separate RNA sequencing of the mutant vs control indicated that the original experimenter expected the mutation to cause the gene expression to differ from the wild-type. To perform this filtering a datacard was created for each BioProject listing the BioProject Title and Description and each Experiment. For each BioSample, the datacard listed the tissue, treatment, Sample Name, Experiment ID, BioSample ID, cultivar, replication, genotype and dev_stage in YAML format. A system and user prompt were created to submit together with each BioProject data card individually to the OpenAI model o3. The prompt requested one replicate of each biological condition with mutants excluded. To sense-check the results the model was asked to also return a concise matrix summary of the BioProject study design, its reasoning for its selection of the experiments and a confidence score from 1–5, with a semantic interpretation of the scoring scheme provided in the prompt. All BioProjects whose analysis scored below 4 out of 5 (“data essentially complete, minimal inference to interpret any abbreviations”) were discarded. As an additional sense check, 10 processed soybean BioProjects were selected at random and were curated by an expert to ensure agreement with the experiments selected by the model. After this final filtering step, the numbers of RNA-seq experiments were: *A. thaliana*: 1639, *Z. mays*: 813, *G. max*: 576, *G. hirsutum*: 286, *T. aestivum*: 762 and *O. sativa*: 1356.

Each RNA-seq experiment was aligned to the reference genome and corresponding GTF annotation file of the species using STAR [114]. To generate the genome index, the following options were used:

~~~
STAR --runMode genomeGenerate \
   --genomeFastaFiles GENOME_FILE \
   --limitGenomeGenerateRAM 300000000000 \
   --sjdbGTFfile GTF_FILE \
   --sjdbOverhang 299 \
   --genomeSAindexNbases VALUE
~~~

where `VALUE` was set to the recommended min(14, log_2_(GenomeLength)/2 − 1).

FASTQ files were trimmed using Trim Galore [115] with the options “`–quality 20 –length 20`“. The trimmed FASTQ files were then aligned to the indexed genome using STAR with default parameters and the option “`–outSAMtype BAM SortedByCoordinate`“.

Coverage tracks were generated using bamCoverage [116] with the options “`–normalizeUsing CPM –binSize 1`“.

##### B.4.6. Plant ATAC- and ChIP-seq Tracks

Similarly to the RNA-seq track processing, ATAC- and ChIP-seq tracks and their associated metadata were queried from the NCBI SRA. A comparable set of filters was applied, including `LibraryLayout` equal to `PAIRED`, `Bases` greater than or equal to 6 × 10^9^, and `AvgSpotLen` greater than or equal to 300 base pairs. As with RNA-seq track processing, mutants were excluded by selecting only wild-type samples from the `Genotype` (or `Cultivar`) field. For *A. thaliana*, this included any entry containing the string “Columbia 0”; for *Z. mays*, the string “B73”; for *G. max*, the string “Williams 82”; for *G. hirsutum*, the string “G. hirsutum acc. J668”; for *T. aestivum*, the string `Chinese Spring`; and for *O. sativa*, the string “japonica.” In addition, any sample annotated as `wild type` or similar variations (e.g., `wt`) was also included. Given that the number of ATAC- and ChIP-seq experiments is much smaller than those of RNA-seq, we manually curated the missing metadata entries for all experiments that passed the initial filters to obtain reliable tissue, and for ChIP-seq experiments, transcription factor or histone mark annotations.

Then each ATAC- and ChIP-seq experiment was processed using the following pipeline:

1. FASTQC [117]: to assess the quality of raw sequencing reads.
2. Trim Galore [115] with the options `-q 20 –phred33 –stringency 3 –length 20 -e 0.1`: to trim adapter sequences and low-quality bases.
3. Bowtie2 [112] with the options `-X2000`: to align reads to the reference genome, allowing for paired-end mapping with a maximum fragment length of 2000 bp.
4. Python script: to count the number of reads mapping to nuclear versus organellar genomes, enabling assessment of contamination from mitochondrial or chloroplast DNA.
5. SAMtools [113] with the options `-f 2 -F 1804 -q 30`: to filter for properly mapped, paired reads with MAPQ ≥ 30, followed by `fixmate -r` to correct mate information, and a second filtering with `-f` `2 -F 1804` to remove orphan or inter-chromosomal reads.
6. Bash script: to compute library complexity, which estimates the number of unique fragments and evaluates sequencing depth sufficiency.
7. Picard’s MarkDuplicates [118]: to identify and remove PCR duplicates, preventing bias from overrepresented fragments.
8. Python script: to calculate fragment length distribution on deduplicated reads, which is particularly important for ATAC-seq to infer nucleosome positioning.

We note that the Python and Bash scripts mentioned above were obtained from the documentation of ENCODE’s ATAC-seq pipeline. After processing, the following quality control filters were applied: experiments with “Basic Statistics” and “Per Base Sequence Quality” marked as “Pass” in the FASTQC report; experiments with a Bowtie2 alignment rate ≥ 0.8; experiments with an organelle read fraction < 0.5; experiments meeting library complexity criteria (PBC1 ≥ 0.5, PBC2 ≥ 1, NRF ≥ 0.5); experiments with a duplication rate < 0.4; experiments showing the presence of both NFR and mono-nucleosome peaks in the fragment length distribution; and experiments passing a read-depth filter to ensure sufficient sequencing coverage, using species-specific thresholds scaled from the ENCODE human guideline of 25 million mapped reads according to genome size.

After this final filtering step, the numbers of ATAC-seq and ChIP-seq experiments were: *A. thaliana*: 260, *Z. mays*: 108, *G. max*: 14, *G. hirsutum*: 33, *T. aestivum*: 14 and *O. sativa*: 36.

For ATAC-seq experiments, a Tn5-shifted BAM file was generated to accurately represent insertion sites. Specifically, the BAM file was processed with alignmentSieve [116] using the `–ATACshift` option to adjust reads according to the Tn5 transposase cut site. The resulting BAM was then sorted and indexed using SAMtools to prepare it for downstream analysis. An insertion signal track with 1-bp resolution was generated without centering or extending the reads. Finally, bamCoverage [116] was used to create a normalized bigWig track (`–normalizeUsing CPM`) at 1-bp bin size, representing the Tn5 insertion profile across the genome. For ChIP-seq experiments, two types of coverage tracks were generated depending on the target. For transcription factors or other proteins producing narrow, well-defined peaks, reads were centered using bamCoverage (`–centerReads`) and CPM-normalized at 1-bp bin size. For histone modifications, the full fragment coverage was used without centering and similarly CPM-normalized at 1-bp bin size.

##### B.4.7. Domesticated Animals RNA-seq, ChIP-seq, and ATAC-seq Tracks

RNA-seq data for chicken, horse, pig, and sheep were queried from the NCBI SRA database and limited to published studies that are part of the Functional Annotation of Animal Genomes (FAANG) Consortium. Among these, we prioritized studies that included the same tissues or cell types across species to minimize potential variability in sampling conditions. Quality-control filters were similar to those applied to the plant RNA-seq datasets, including `LibraryLayout` equal to `PAIRED` and `AvgSpotLen` greater than or equal to 200 base pairs. After filtering, each RNA-seq experiment was processed using the same RNA-seq pipeline as that used for the plant RNA-seq experiments. Because many RNA-seq experiments for a given species included multiple experiments involving the same tissues, we limited these to a single experiment per tissue by selecting the experiment with the highest mapping rate, based on the output from STAR.

ChIP-seq tracks for chicken, horse, pig, and sheep were downloaded from the Visualisation FAANG portal (https://viz.faang.org/experiments/). The pre-processed ChIP-seq tracks were generated using an nf-core ChIP-seq sub-workflow and therefore adhere to standard bioinformatics best practices for this type of omics data. ATAC-seq data were processed similarly to the RNA-seq data by querying the NCBI SRA database and limiting the datasets to published studies that are part of the FAANG Consortium. We applied quality-control filters similar to those used for the plant ATAC-seq experiments and processed the data using the same ATAC-seq pipeline as that used for the plant ATAC-seq experiments. For the ATAC-seq experiments, after processing, we then also used the same post-processing quality-control filters as mentioned in the plants dataset above. As for the RNA-seq experiments, ATAC-seq experiments were also limited to a single experiment per tissue, by selecting the experiment with the highest mapping rate. The final numbers of RNA-seq, ChIP-seq, and ATAC-seq experiments were: chicken: 18, 15, 22; horse: 31, 36, 29; pig: 19, 36, 39; sheep: 56, 5, 2, respectively.

##### B.4.8. Zebrafish and Worm RNA-seq and ATAC-seq Tracks

RNA-seq data for zebrafish were obtained from a high-resolution gene expression time course of embryonic development in zebrafish [119]. Raw data were obtained from the NCBI SRA under BioProject accession number PRJEB12982. ATAC-seq data for zebrafish were obtained from an embryonic development study covering five developmental stages [120]. Raw data were obtained from the NCBI SRA under BioProject accession number PRJNA416977. RNA-seq data for *C. elegans* were obtained from a Waterston laboratory modENCODE study profiling the transcriptome across developmental stages under various experimental conditions. Only experiments performed under normal conditions were retained. Raw data were obtained from the NCBI SRA under BioProject accession number PRJNA33023. ATAC-seq data for *C. elegans* were obtained from a chromatin accessibility atlas covering developmental stages and aging [121]. Raw data were obtained from the NCBI SRA under BioProject accession number PRJNA471351. The same RNA-seq and ATAC-seq pipelines described above were used to process the data for both species. For the ATAC-seq experiments, the same post-processing quality-control filters described above for the plant ATAC-seq datasets were also applied. Finally, a single experiment per tissue or developmental stage was retained based on the highest mapping rate. The numbers of experiments retained were 18 RNA-seq and 5 ATAC-seq for zebrafish, and 38 RNA-seq and 4 ATAC-seq for *C. elegans*.

#### B.5. NTv3 Benchmark Data Sources and Processing

The NTv3 Benchmark tasks are defined in Methods 4.6. This note gives the accessions and processing pipeline for the species-specific tracks collected for it. The plant pipelines these sections refer to are described in Supplementary Note B.4 above.

##### B.5.1. Tomato ATAC- and RNA-seq Tracks

For tomato (*S. lycopersicum*), ATAC-seq and RNA-seq data were sourced from a study of regulatory regions during fruit development [122]. The two assays cover different tissue panels. The seven ATAC-seq experiments span leaf, anthesis fruit, fruit at 3, 5 and 10 days post-anthesis, and fruit at 3 and 7 days before anthesis. The thirteen RNA-seq experiments cover fruit only, at anthesis and at 5, 10 and 15 days post-anthesis, with several experiments per stage. Raw data were obtained from the NCBI SRA (BioProject PRJNA1228166). Data processing followed the same pipelines and quality filters as for chicken and plant experiments. ATAC-seq reads were Tn5-shifted using the `–ATACshift` option, and both ATAC-seq and RNA-seq tracks were generated as normalized CPM bigWig files at 1-bp resolution using `bamCoverage`.

##### B.5.2. Maize, Rice, and Thale Cress Ribo-seq Tracks

Plant protein abundance data were derived from Ribo-seq assays for three species: *Z. mays, O. sativa*, and

*A. thaliana*. The experiments on these plant species were conducted across multiple tissues and developmental stages. Raw data for all species were accessed via the NCBI SRA under the BioProject accession number PRJNA637713.

Each Ribo-seq experiment was subsequently processed using the following pipeline:

1. FASTQC [117]: to assess the quality of raw sequencing reads.
2. Trim Galore [115] with the options `-q 20 –phred33 –stringency 3 –length 20 -e 0.1`: to remove adapter sequences and low-quality bases from the reads.
3. Bowtie2 [112]: to deplete reads mapping to rRNA and tRNA, retaining only putative mRNA reads.
4. HISAT2 [123] with default parameters: to align ncRNA-depleted reads to the reference genome.
5. SAMtools [113] with the options `view -bS -q 30`: to filter for high-quality alignments.

After running the pipeline, only experiments that passed FASTQC quality filters, had an alignment rate ≥0.5, and contained more than 2 million reads were retained. As with the ATAC-seq and RNA-seq tracks, `bamCoverage` [116] was used to generate a normalized CPM bigWig track at 1-bp resolution for all Ribo-seq experiments that passed quality filters.

##### B.5.3. Cattle and Rat Functional Tracks

The RNA-seq and ATAC-seq data for *Bos taurus* were obtained from the NCBI SRA database and restricted to published studies that are part of the Functional Annotation of Animal Genomes (FAANG) Consortium and cover a broad range of tissues. The NCBI SRA BioProject accessions were PRJEB27455 and PRJEB14330 for RNA-seq, and PRJEB27111, PRJNA531214, PRJNA605780, and PRJNA665194 for ATAC-seq. ChIP-seq tracks, as for the other domesticated animal species used for post-training, were downloaded from the Visualisation FAANG portal, which provides pre-processed ChIP-seq tracks generated using an nf-core ChIP-seq sub-workflow. The final numbers of RNA-seq, ChIP-seq, and ATAC-seq experiments for cattle were 11, 87, and 11, respectively. The *Rattus norvegicus* RNA-seq data were obtained from a previously published study of regulatory dynamics in mammalian tissues [124]. The raw data were accessed through the NCBI SRA under BioProject accession PRJNA177791. The ATAC-seq data were obtained from a previously published comparative study [125], with the raw data accessed through the NCBI SRA under BioProject accession PRJNA670507. RNA-seq and ATAC-seq data were processed using the pipelines described above, and the same post-processing quality-control filters were applied to retain only high-quality experiments. When multiple replicates were available for the same tissue, only the experiment with the highest read count was retained. The final numbers of RNA-seq and ATAC-seq experiments for rat were 9 and 3, respectively.

##### B.5.4. Fungi RNA-seq and ATAC-seq Tracks

*S. cerevisiae* RNA-seq raw data were obtained from the NCBI Sequence Read Archive (SRA) under BioProject accession PRJEB35903. These data were generated from an ethanol-grown yeast cell-cycle experiment, with samples collected at 14, 20, 92, 102, 132, 142, and 162 minutes. *S. cerevisiae* ATAC-seq data were obtained from a previously published study [126] (BioProject PRJNA276699). The dataset is a 0.6 M NaCl osmotic-stress time course of five samples, collected at 0, 15, 30, 45, and 60 minutes after treatment, the 0-minute sample being the untreated control. *S. pombe* RNA-seq raw data were obtained from the NCBI SRA under BioProject accession PRJEB11998. These data include experiments performed under normal nitrogen conditions and during nitrogen starvation. *S. pombe* ATAC-seq data were obtained from a previously published study [126] (BioProject PRJNA276699). The *S. pombe* ATAC-seq experiments represent the same unstressed biological condition but were generated using different experimental preparation protocols, varying the number of cells used for library preparation (1, 5, or 20 million cells) and the duration of the spheroplasting treatment. RNA-seq and ATAC-seq data were processed using the pipelines described above, and the same post-processing quality-control filters were applied to retain only high-quality experiments. When multiple replicates were available for the same condition or time point, only the experiment with the highest read count was retained.

##### B.5.5. Protists RNA-seq and ATAC-seq Tracks

*Plasmodium falciparum* RNA-seq data were obtained from a previously published study of blood-stage development [127]. The data include the ring (10–15 hours post infection [hpi]), trophozoite (25–30 hpi), and schizont (40–45 hpi) developmental stages. Raw data were accessed through the NCBI SRA under BioProject accession number PRJNA765585. ATAC-seq data were obtained for four developmental stages (10, 20, 30, and 40 hpi), with raw data accessed through the NCBI SRA under BioProject accession number PRJNA524965. The *Dictyostelium discoideum* RNA-seq data were obtained from a previously published developmental study [128]. The experiments included developmental time points at 0, 4, 8, 12, 16, 20, and 24 hours. Raw data were accessed through the NCBI SRA under BioProject accession number PRJNA640642. RNA-seq and ATAC-seq experiments were processed using the same pipelines described above, and the same post-processing quality-control filters were applied to retain only high-quality experiments. For developmental stages with multiple replicates, only the experiment with the highest read count was retained.

#### B.6. Genome Annotation: Decoding, Protein-level Analyses and Runtime

##### B.6.1. Decoding Algorithm

###### States

The decoder runs over the fifteen classes the annotation head emits: `intergenic`; `CDS_p0..p2`, where *p* is the codon position of that base; `intron_p0..p2`, where *p* is the phase inherited from the upstream donor; `splice_donor_p0..p2`, the last base of a non-final coding exon; `splice_acceptor_p0..p2`, the first base of a non-initial coding exon; and `start_codon` and `stop_codon`, single bases at the two ends of the coding extent. Every state except `intergenic` and the three introns is a one-base state with no self-loop, so a multi-base run of one label in the reference decodes as that state followed by its successors. For the minor-spliceosome (U12) introns, the decoder adds nine states, `splice_donor_u12_p0..p2`, `intron_u12_ p0..p2` and `splice_acceptor_u12_p0..p2`, which mirror their canonical counterparts phase for phase. They have no output class of their own: each reads the head’s log-posterior for its canonical counterpart and is relabeled as that counterpart after decoding, so the decoded path uses the fifteen classes only.

###### Transitions

Supplementary Fig. A.14 gives every non-zero transition; each state’s outgoing probabilities sum to one and all omitted transitions are zero, hence −∞ in log space. Three consequences are worth naming. The coding states form a three-cycle with no self-loop, so codon position advances at every base. The intron detour returns to the phase it left (the only intron path from `CDS_pN` is `donor_p`(*N*+1) → `intron_p`(*N*+1) → `acceptor_p`(*N*+2) → `CDS_pN`, the donor and acceptor taking the two intervening codon positions) so frame runs unbroken across a junction of any length, and transitions between intron phases are forbidden rather than penalized. And `stop_codon` is reachable from `CDS_p1` alone, which is why that state spends the exon-exit prior twice, and which forces the coding extent to a whole number of codons: the frame constraint is topological, not a downstream check.

###### Emission constraints

The emission score of state *j* at position *t* is the head’s log-posterior plus an additive mask *m*_*t*_(*j*) that vetoes states the local sequence cannot support: log *e*_*t*_(*j*) = log *p*_*t*_(*j*) + *m*_*t*_(*j*). Offsets below are relative to the state’s own base.

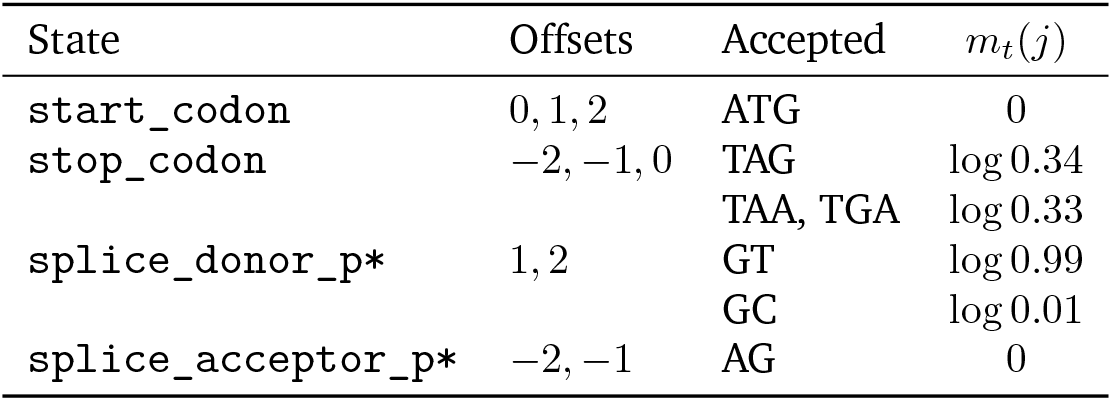

| State | Offsets | Accepted | $m_t(j)$ |
| --- | --- | --- | --- |
| start_codon | 0, 1, 2 | ATG | 0 |
| stop_codon | −2, −1, 0 | TAG<br>TAA, TGA | log 0.34<br>log 0.33 |
| splice_donor_p* | 1, 2 | GT<br>GC | log 0.99<br>log 0.01 |
| splice_acceptor_p* | −2, −1 | AG | 0 |

A position matching no accepted pattern is set to −∞, as is one whose context falls outside the window. Where any base of the context is `N`, the constraint is waived and the mask is 0, so assembly gaps neither force nor forbid a site. Unconstrained states carry no mask. Repeat soft-masking is not waived: lower-case bases are read as their upper-case codes and are constrained normally.

###### Viterbi decoding

Decoding maximizes the joint score of a state path over the lattice, in log space throughout. Writing *a*_*ij*_ for the transition log-probabilities of Supplementary Fig. A.14 and log *e*_*t*_(*j*) for the emission score of state *j* at position *t*, the recursion is

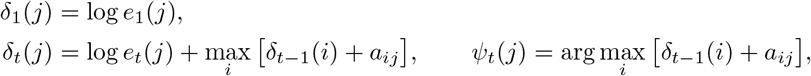

with *q*_*T*_ = arg max_*j*_ *δ*_*T*_ (*j*) and *q*_*t*_ = *ψ*_*t*+1_(*q*_*t*+1_). The initial distribution is uniform, so *δ*_1_ is the emission alone, and the final state is unconstrained. Tiberius’s HMM layer also runs the forward and backward recursions, so that gradients pass through it during end-to-end fine-tuning, although its parameters are frozen [28]. Our decoder is inference-only: posteriors come from the network and only the Viterbi recursion is evaluated.

###### Blocked evaluation

The recursion is a matrix–vector product in the (max, +) semiring, where (*u* ⊗ *A*)_*j*_ = max_*i*_(*u*_*i*_ + *A*_*ij*_) and (*A* ⊗ *B*)_*ij*_ = max_*k*_(*A*_*ik*_ + *B*_*kj*_). Because that product is associative it can be reassociated, which is what allows a sequence of *T* positions to be evaluated in three passes of bounded depth rather than *T* sequential steps. Positions 2, …, *T* are partitioned into *C* chunks of length *L* (the last padded and masked), giving a depth of *O*(*L* + *T* /*L*), minimized near 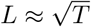; *L* defaults to 512.

###### Pass 1 — chunk transfer matrices, all chunks in parallel

For each chunk *c* we build *M*^(*c*)^ ∈ ℝ^*K*×*K*^, where 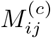 is the best score of a path that enters chunk *c* in state *i* and leaves it in state *j*, measured relative to the entry score. Starting from the semiring identity *I* (0 on the diagonal, −∞ elsewhere), the *L* positions of every chunk are advanced together:

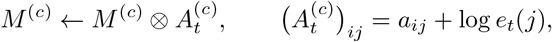

sequential in *L* but independent across the *C* chunks.

###### Pass 2 — entry scores, sequential across chunks

An exclusive scan carries the running score vector over whole chunks at a time, emitting each chunk’s entry score:

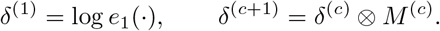

This is the only pass whose length is *C* rather than *L*, and it performs one *K* × *K* product per chunk instead of *L* per-position steps.

###### Pass 3 — replay with backpointers, all chunks in parallel

Each chunk is re-run from its now-known entry score *δ*^(*c*)^, this time recording *ψ*_*t*_(*j*) = arg max_*i*_[*δ*_*t*− 1_(*i*) + *a*_*ij*_] at every position. Because every chunk starts from a score vector fixed by pass 2, the chunks are again independent.

###### Backtrace

Each *ψ*_*t*_ is a map from states to states, and map composition is associative, so the backtrace is not a sequential walk: the reversed sequence *ψ*_*T*_, …, *ψ*_2_ is composed by an associative scan in *O*(log *T*) depth, and the whole path is then read off by applying each composed map to the terminal state *q*_*T*_ .

###### Numerical stability and exactness

After every step the scores are shifted by a single scalar per sequence and chunk, the maximum over the state axes. A uniform shift cancels in every subsequent comparison, so the decoded path is unchanged while running magnitudes stay bounded near zero; without it a float32 accumulation loses the optimal path at megabase scale. A floor at −10^30^ prevents an all-vetoed row from producing −∞ − (−∞). The blocked evaluation is exact rather than approximate because the per-chunk score combination is associative, and the chunk length is a speed parameter with no effect on the result: the decoded path matches a float64 sequential reference across sequence and chunk lengths, and the emitted GFF3 is byte-identical for chunk lengths from 64 to 100,000.

###### Windowing

The lattice spans one window’s prediction region, and windows are decoded independently: no lattice crosses a window boundary and the resulting state paths are concatenated without re-decoding. The stitched path is therefore not guaranteed to be globally optimal and may contain an illegal transition at a seam; nothing repairs seams, and the only post-decode steps are the two gene-level filters, which drop predicted genes with under 201 bp of CDS and those whose spliced CDS carries an internal in-frame stop.

##### B.6.2. Protein-level Analyses

###### Protein extraction

The CDS features of each gene were concatenated in ascending genomic order, reverse complemented for minus-strand genes, trimmed by the phase of the transcript’s first CDS and truncated to a whole number of codons, then translated with the standard genetic code, with the terminal stop removed. Proteins shorter than 20 residues were discarded. Protein identity was computed from a global alignment with BLASTP scoring, as identical residues divided by the length of the longer protein, so that a prediction is charged for residues it omits; recovery is reported at exact, ≥95%, ≥90% and ≥50% identity. Because translation requires per-contig sequence, the proteomes cover the staged contigs of each assembly, which hold 99.3% of cow and 98.9% of beluga whale reference loci; the same contig set was used for the reference and both callers.

###### BUSCO completeness

Supplementary Table B.2 gives the marker counts and completeness of each proteome. In cow, NTv3 recovers 98.1% of markers against 98.9% for the unfiltered reference proteome, and 96.0% against 97.7% in beluga whale; Tiberius recovers 97.0% and 95.6%. Both callers therefore sit below the curated annotation in both species, by 0.8 and 1.7 points for NTv3 and by 1.9 and 2.1 points for Tiberius, so the interpretable quantity is each caller’s shortfall against the same ceiling rather than an absolute completeness. NTv3’s lead over Tiberius comes mostly from fewer fragmented markers (23 against 82 in cow, 50 against 103 in beluga whale); it also leaves fewer markers missing in cow (152 against 190) but slightly more in beluga whale (320 against 303).

**Supplementary Table B.2.** BUSCO completeness of the held-out mammal proteomes. Marker counts against `mammalia_ odb10` (*n* = 9,226) and completeness (complete single-copy plus duplicated) as reported by BUSCO v5.7.1 in protein mode. The unfiltered reference proteome, built from the complete reference annotation before scoring-time filtering, is the ceiling; shortfall is in percentage points below it.

| Species | Proteome | Complete single-copy | Complete duplicated | Fragmented | Missing | Complete (%) | Shortfall (points) |
| --- | --- | --- | --- | --- | --- | --- | --- |
| Cow | NTv3 | 8,941 | 110 | 23 | 152 | 98.1 | 0.8 |
|  | Tiberius | 8,849 | 105 | 82 | 190 | 97.0 | 1.9 |
|  | Unfiltered reference | 8,983 | 138 | 51 | 54 | 98.9 | – |
| Beluga whale | NTv3 | 8,762 | 94 | 50 | 320 | 96.0 | 1.7 |
|  | Tiberius | 8,717 | 103 | 103 | 303 | 95.6 | 2.1 |
|  | Unfiltered reference | 8,922 | 91 | 77 | 136 | 97.7 | – |

##### B.6.3. Runtime

###### Inference precision

Training was performed in FP32. At inference the convolution, attention-projection and feed-forward matrix multiplications run in bfloat16 while the stored weights and the annotation head remain in FP32. On the whole cow assembly this runs 1.7 times faster than FP32 and changes gene- and CDS-level F1 by 0.03 (58.05 and 90.62 in bfloat16 against 58.08 and 90.65 in FP32). Assembly staging is excluded from the timing boundary as a property of our storage rather than of the tool, and repeat masking because it is a prerequisite shared by every caller.

###### Fixed cost and padding

Runtime is fitted on input windows rather than summarized as a single seconds-per-Mb value because each run carries a fixed cost of about 80 s, most of it compilation, small for a multi-gigabase genome but about half the runtime for a 40-Mb contig. The fit implies a marginal cost of 0.94 s per Mb of contiguous sequence, below the per-assembly means in the Results, which include the fixed cost. Each contig is padded to whole windows, so assembly size alone fits poorly (*r*^2^ = 0.85): on the cow assembly, contigs longer than 10 Mb ran at 0.93 s/Mb and those of 10 Mb or less at 16.4 s/Mb, at the same cost per window. Runtime comparisons on fragmented assemblies should therefore name the assembly rather than only the species.

##### B.7. Enhancer–Gene Linking across Distance Bins

To compare enhancer–gene ranking, we evaluated NTv3, Borzoi fold 0 and published AlphaGenome scores on K562 CRISPRi element–gene pairs from the ENCODE-rE2G benchmark [48], on all 10,356 pairs (unscored pairs set to zero) and on 3,826 matched pairs scored by all three models. We stratified pairs by enhancer-to-TSS distance because genomic distance is itself strongly predictive of regulatory interaction and because the number of experimentally supported positives decreases sharply in the most distal bin (Supplementary Fig. A.21a). All pairs comprised 223, 1,597 and 8,536 pairs with 133, 256 and 82 positives at < 10 kb, 10–100 kb and 100–2,500 kb; the matched set comprised 218, 1,524 and 2,084 pairs with 128, 240 and 42 positives.

At < 10 kb, all three models performed strongly, with auPRCs (all / matched pairs) of 0.86/0.87 for NTv3, 0.89/0.89 for Borzoi and 0.91/0.91 for AlphaGenome. At 10–100 kb, auPRC was 0.53/0.55, 0.59/0.60 and 0.77/0.78, respectively. In the 100–2,500 kb bin, NTv3 reached 0.24/0.41 compared with 0.21/0.37 for Borzoi and 0.35/0.64 for AlphaGenome. Precision at 70% recall showed the same broad pattern: 0.84/0.85, 0.82/0.83 and 0.86/0.86 at < 10 kb; 0.38/0.43, 0.45/0.47 and 0.74/0.75 at 10–100 kb; and 0.02/0.14, not defined/0.09 and 0.02/0.20 at 100–2,500 kb for NTv3, Borzoi and AlphaGenome, respectively. auROC remained high for all three models on matched pairs across distance bins (0.82–0.91; 0.68–0.91 on all pairs), whereas the precision–recall metrics separated them more clearly in the intermediate and distal regimes.

Because the number of positives is small at long range, we emphasize the threshold-free precision–recall curves together with a common 70% recall operating point rather than model-specific classification thresholds (Supplementary Fig. A.21b,c). NTv3 and Borzoi occupy a similar performance regime across the matched pairs, while AlphaGenome separates more clearly at intermediate and distal distances. Thus, NTv3 captures enhancer–gene regulatory signal across genomic distances, but the improved functional-track performance of NTv3 does not translate into a consistent advantage for enhancer–gene ranking.

### C. Supplementary Table Legends

**Supplementary Table C.1** | Details of functional tracks used in post-training per species.

**Supplementary Table C.2** | Details of species used for NTv3 annotation models training.

**Supplementary Table C.3** | Details of annotation performance and runtime.

**Supplementary Table C.4** | Details of NTv3 Benchmark tasks and test Pearson correlation of every evaluated model on each task, for each fine-tuning seed.

**Supplementary Table C.5** | Per-track test Pearson correlation of functional-track post-training initialized from the pre-trained model or randomly, for all 15 species.

**Supplementary Table C.6** | Per-track test Pearson correlation on held-out human and mouse windows of post-training on human and mouse only or on all 15 species.

**Supplementary Table C.7** | Per-track Pearson correlation of NTv3, Borzoi and AlphaGenome on Borzoi’s human tracks, on folds 3 and 4, at 1-bp (zero-shot) and 32-bp (fine-tuned) resolution.

**Supplementary Table C.8** | Per-track test Pearson correlation of NTv3 on AlphaGenome’s human single-base tasks, with AlphaGenome’s published assay-level means.

**Supplementary Table C.9** | Per-task performance of NTv3 and published baselines on the SeiPlant, Agronomics, DeepFARM and DanioDecima benchmarks.

**Supplementary Table C.10** | Per-tissue eQTL causality auROC and auPRC of NTv3 and Borzoi across 48 GTEx tissues, with tissue-weighted aggregates.

